# Spatial μProBe: a correlative multimodal imaging approach for spatial profiling of biological micro-environments

**DOI:** 10.1101/2025.04.23.650252

**Authors:** Francisco C. Marques, Neashan Mathavan, Nicholas Ohs, Christopher Goenczoel, Jack J. Kendall, Amit Singh, Dilara Yilmaz, Denise Günther, Gisela A. Kuhn, Friederike A. Schulte, Esther Wehrle, Ralph Müller

## Abstract

Despite fundamental advances in spatial omics, investigating cellular and molecular functions within their native environment remains a challenge in multiscale systems biology, especially in response to organ-level events. Here, we introduce Spatial µProBe (micro-ProBe), a multimodal imaging approach for spatial profiling of biological micro-environments. Spatial µProBe enables preprocessing, registration and correlative analysis of 2D and 3D imaging modalities, supported by an intuitive user interface. As an application, we investigated bone mechanobiology and characterised the cellular and molecular responses to mechanical loading during adaptation and regeneration, which continuously regulate the local microarchitecture. By integrating time-lapsed micro-computed tomography and end-point spatial transcriptomics, we profiled the local mechanical in vivo environment of thousands of musculoskeletal cells, revealing the spatiotemporal interplay between local mechanics and gene expression driving tissue development. Spatial µProBe marks a crucial advance in the characterisation of multiscale tissues and signalling, facilitating the exploration of targeted molecular therapies for pathological conditions.

## Introduction

Multimodal imaging (MI) approaches enable correlating complementary views of the same specimen across spatial scales^1^. Typical workflows involve imaging a region of interest (ROI) with two or more technologies, followed by image registration to align all datasets into a shared reference frame. Many established 2D and 3D imaging modalities have been successfully correlated, such as magnetic resonance imaging (MRI)^2–4^, computed tomography (micro-CT and CT)^2,3^, and light (LM)^3–5^, confocal (CM)^2,3^ and electron (EM)^5^ microscopy. The wide range of imaging capabilities, from structural to functional representations, and differences in spatial resolutions underline the discovery potential of integrating multiscale multimodal data. Indeed, integrating data from MI is becoming paramount in systems biology, with noteworthy examples in neuroscience (e.g., to study stroke^2^, connectivity maps^2^, and brain structure^4^), vascular biology (e.g., to investigate atherosclerosis^6^ and map vascular micro-environments^3^), or oncology (e.g. to characterise micro-environments in breast cancer^7^, or track tumour growth^3^), across preclinical^2,3^ and clinical^6,7^ applications. Some approaches^2,3^ have also combined in vivo and ex vivo imaging steps, aiming to represent time-dependent biological processes in their native environment. Yet, accessing and integrating high-resolution information at the cellular and molecular scales remains a challenge for achieving a holistic understanding of biological systems.

Notably, recent technological advances have established spatial transcriptomics (ST) methods that combine tissue imaging with a detailed profiling of gene expression within its native spatial context and tissue architecture. Typically, a slice of biological tissue is imaged and placed on a spatially barcoded array, and the mRNA of cells within each spot is sequenced^8^. For example, the 10x Genomics Visium system contains nearly 5,000 spots in one standard capture area, enabling the unbiased detection of almost 20,000 genes of the human and mouse transcriptomes at cellular resolution. This tool has contributed to democratising ST across many fields (e.g., oncology^9^, neuroscience^10^, immunology^11^ and developmental biology^12^), making it a pivotal technology to incorporate into MI approaches. To realize the potential of ST, standardised and reproducible methods are needed to integrate, analyse, and visualise associated large-scale datasets. Notable examples include MultI-modal Spatial Omics^13^ (MISO) and SpatialData^14^ (within the scverse^15^ ecosystem) which provide comprehensive workflows to process ST data and emphasise multimodal data integration. These methods established foundational tools to integrate 1D and 2D multi-omics data (e.g., single-cell RNA sequencing, microscopy imaging) with 2D ST datasets, consolidating ST as a valuable technology to reveal local and spatially-resolved biological insights^16^.

However, existing MI solutions often fall short in seamlessly integrating 3D tissue information, limiting their ability to accurately characterize complex biological processes. One example is bone mechanobiology, where organ-level mechanics influence the interplay and the local mechanical in vivo environment (L*iv*E) of cells in bone^17,18^. Mechanobiology is especially vital in developmental^19^, adaptive^20^ or regenerative^21^ phases, where a successful outcome hinges on the precise cellular coordination and responses to mechanical loading across L*iv*E profiles in highly heterogeneous mineralisation states. Likewise, ageing and bone degenerative conditions often compromise bone mechanoregulation^22^, which may contribute to bone fragility and an increased fracture risk, although the underlying mechanisms are not fully understood. Currently, only micro-finite element (micro-FE) analysis can reproduce 3D maps of the in vivo mechanical environment. These local mechanical signals have been successfully linked to tissue structural changes observed in time-lapsed in vivo micro-CT images during bone adaptation^20^ and regeneration^23^. While it is possible to visualise selected cell types in their in vivo environment^24^, an exhaustive profiling of all cells involved in bone mechanoregulation could be transformative for identifying relevant signalling pathways. ST is a promising method despite technical challenges associated with the calcified nature of bone. Since mechano-molecular responses are regulated at multiple levels, from transcription to protein degradation, an accurate characterisation of these pathways within and across L*iv*E profiles can reveal novel targets for treatments of bone disorders and promoting regeneration. Yet, this spatial mechanomics analysis remains largely unexplored, primarily because of the lack of correlative tools to integrate molecular, functional and anatomical information.

To address this gap, we present Spatial μProBe, a multimodal imaging approach for spatial profiling of biological micro-environments. Spatial μProBe streamlines 2D-3D registration of multimodal datasets, providing a seamless pipeline to preprocess histological sections using deep learning, register 2D and 3D multimodal datasets from the same specimen, and integrate the result for post-processing analysis and visualisation. These features are accessible through a user-friendly web browser interface that yields detailed quantitative outputs. We evaluate the registration performance of Spatial μProBe using gold-standard landmarking-based image registration and with simulated data. We demonstrate its potential by integrating 3D time-lapsed in vivo micro-CT images with (sub-)cellular information from end-point 2D histological and ST data during bone adaptation and regeneration, analysing the local mechanical in vivo environment of thousands of musculoskeletal cells. We show that Spatial μProBe can recapitulate the phases of bone fracture healing at the cellular and molecular levels in an established femoral osteotomy mouse model^21,25^ under physiological and cyclic mechanical loading. Finally, we show how Spatial μProBe can unlock gene mechanoregulation and characterize the spatiotemporal expression of individual genes analytically.

## Results

### Integration of multiscale multimodal datasets

Spatial μProBe is a tool designed for 2D-3D correlative multimodal image analysis (Fig. 1a). Our approach is grounded in three main principles: performance, accuracy and usability. We implemented an efficient pipeline based on semi-binary image registration that requires binary masks of the same tissue for each modality to be registered. A dataset with over ten consecutive 2D histological sections covering the entire field of view of a corresponding micro-CT scan can be analysed within a few hours. Applications relying on 2D histological sections can access and train deep-learning methods to support high-throughput image segmentation of relevant tissue classes based on well-established architectures^26^ or individual cells based on CellPose^27,28^ (Fig. 1b). For a dataset of histological sections with visually similar counterstains (e.g. Fast Green), semantic segmentation of pixels representing bone achieved a Dice score of 0.970 ± 0.004, virtually matching manually produced annotations (Supplementary Note 1). Next, 2D-3D image registration, which achieved sub-voxel accuracy in our validation studies, is accomplished through Iterative Assisted Registration (IterAR), by iteratively optimising the similarity between the 2D image and a 2D section obtained from the 3D image. Affine registration parameters comprising 3D rotations, translations and 2D in-plane stretching are identified until a successful result is achieved, verified by the Dice score and visual inspection between modalities. Image similarity functions can be adjusted at each iteration and are optimised using the Powell optimiser^29^ (Fig. 1c). Regarding usability, the pipeline can be run locally and accessed on a web browser and includes the tools needed to initialise a project, set metadata, preprocess 2D and 3D images to identify relevant ROIs, perform 2D-3D image registration and extract quantitative metrics from the registered data. Crucially, we also leverage recent developments in ST data analysis to incorporate super-resolution methods (iStar^30^) that can predict ST data at the resolution of the 2D-3D registered imaging modalities (e.g., in vivo micro-CT scan; Fig. 1d).

**Figure 1.**
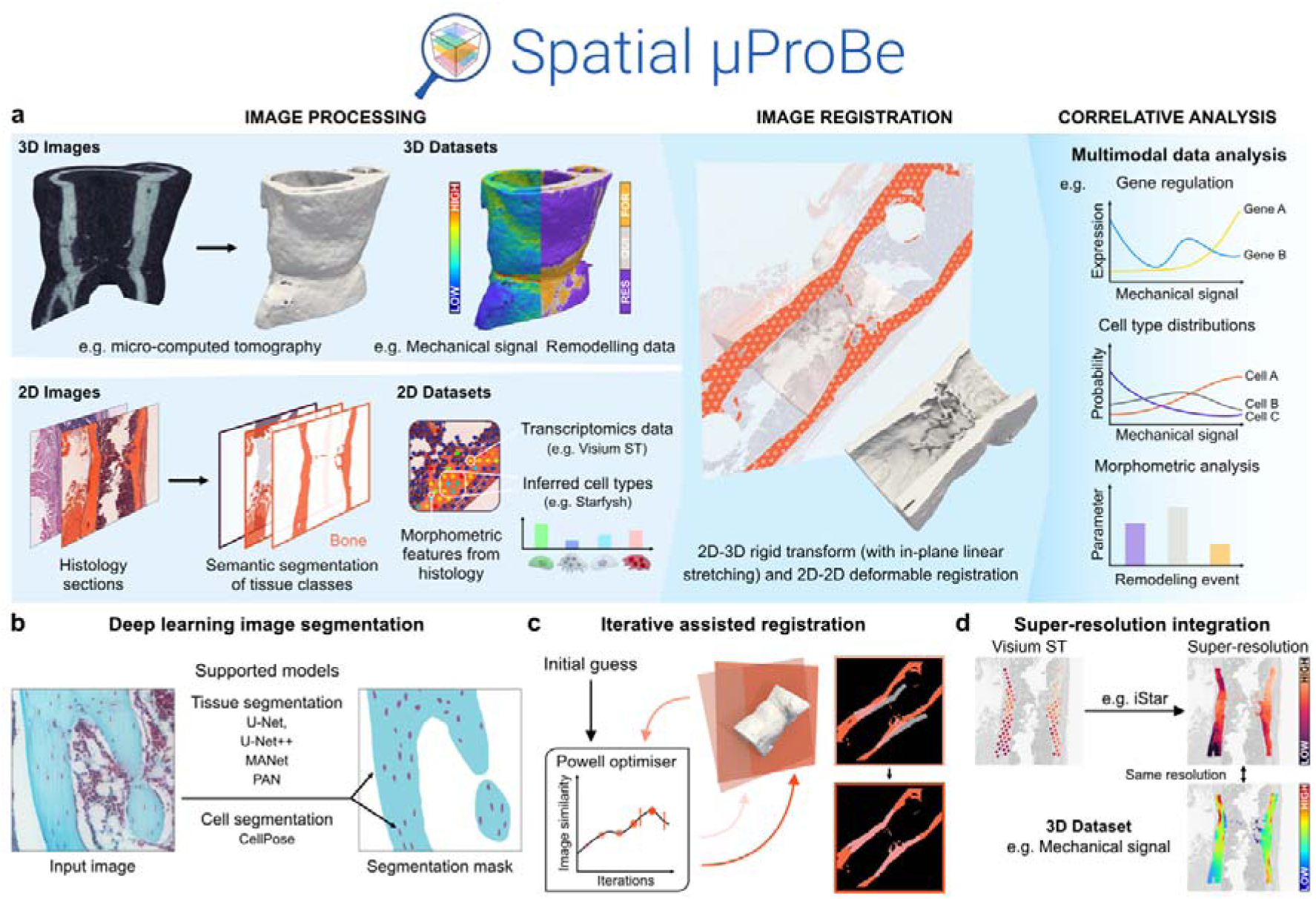
Profiling biological micro-environments with Spatial μProBe. a, The workflow of Spatial μProBe comprises three steps: image preprocessing, 2D-3D image registration and correlative analysis. The input data consists of 2D and 3D images, such as histological sections and micro-computed tomography (micro-CT) images, respectively. These datasets are segmented to generate binary masks for relevant tissues. Binary masked images are 2D-3D registered by searching for optimal transformation parameters that maximise the similarity between the input 2D image and a 2D section extracted from the input 3D image. After a successful registration, relevant quantities from 2D and 3D datasets can be correlated using the spatial mapping provided by Spatial μProBe. In bone mechanobiology, the mechanical signal computed with micro-finite element analysis may be correlated with remodelling events (Formation: FOR, Quiescence: QUI, Resorption: RES) determined from time-lapsed in vivo micro-CT images. b, Spatial μProBe integrates with established deep learning models for 2D semantic segmentation tasks of tissues and cells from histological data. For tissue segmentation, we integrate four architectures (U-Net, U-Net++, Multi-Attention-Network: MANet, Pyramid Attention Network: PAN) from segmentation-models-pytorch, while for cell segmentation we provide an interface for CellPose^27,28^. c, 2D-3D registration is performed using Iterative Assisted Registration (IterAR). IterAR uses the Powell method to optimise the transformation parameters that maximise a user-defined combination of image similarity functions, until a successful result is obtained. An initial guess can be provided to expedite the registration process. d, Post-processing of ST data (from Visium 10x Genomics) in Spatial μProBe supports super-resolution operations (e.g., using iStar^30^) to match the resolution of 3D imaging modalities (e.g., in vivo micro-CT data).

### Validation of 2D-3D registration

We first evaluated the accuracy of 2D-3D registration compared to a gold-standard landmarking-based registration approach. One operator (F.C.M.) identified 132 ± 23 pairs of landmarks per image between six histological sections (Fig. 2a-b, Supplementary Fig. 1a) of three 6^th^ mouse caudal vertebrae and their corresponding high-resolution ex vivo micro-CT scans (voxel-size: 2.1 µm). Matched landmark pairs defined a reference transformation, with an average baseline accuracy of 4.25 ± 0.77 µm (Fig. 2c). Sections were registered with Spatial μProBe at in vivo micro-CT resolution (10.5 µm), achieving an average accuracy of 5.15 ± 0.95 µm (Fig. 2d, no significance difference to the baseline accuracy, p=0.13, two-sided t-test), which is orders of magnitude below the typical size for 3D bone structures measured with micro-CT, such as trabecular thickness (Tb.Th) at 91.25 µm or cortical thickness (Ct.Th) at 161.25 µm. Angular deviations between the normal vectors of the ground truth and optimised planes were below 0.75° for all sections, which is a quarter of the threshold considered in similar evaluations^31^.

**Figure 2.**
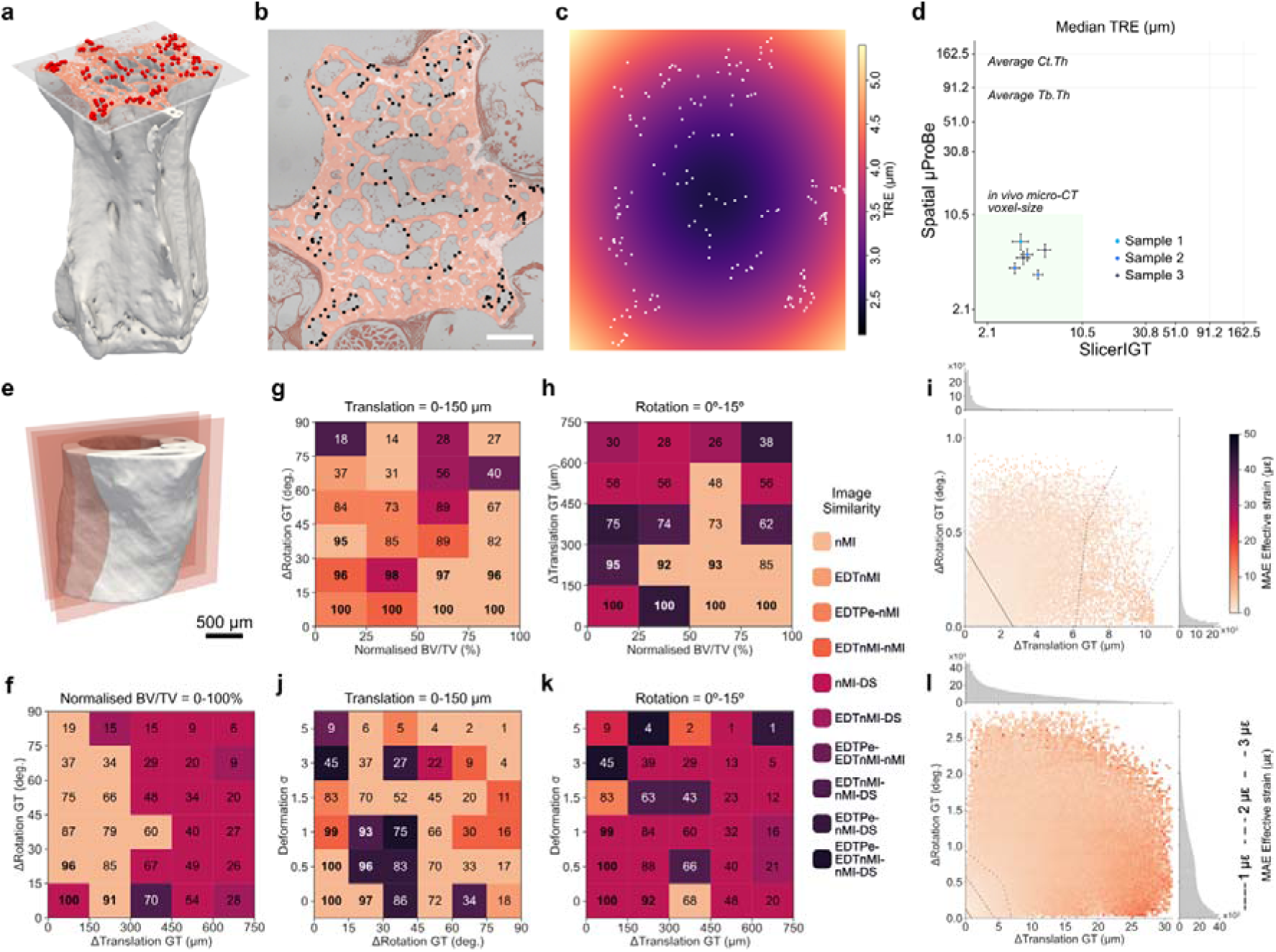
Validation of 2D-3D registration in Spatial μProBe. a, Example of a histological section registered to the ex vivo micro-computed tomography (micro-CT) image (voxel-size: 2.1 µm) using landmarking. Red dots indicate the landmarks (n=102) identified between both modalities (shown in b). b, Landmarks (n=102) positioned on the histological section (shown in a), used for landmarking registration and positioned across the entire section. c, Target Registration Error (TRE) map^5^ computed for all points of the 2D section (shown in a) after landmarking registration, achieving a maximum TRE of 5.5 µm. d, Comparison between the median TRE (and inter-quartile ranges) computed for the registered sections (n=6) using landmarking in SlicerIGT^55^ and Spatial μProBe. For each section, the median TRE was determined, and their average set a baseline accuracy of 4.25 ± 0.77 µm for the registration with Spatial μProBe. An average value for cortical thickness (Ct.Th) and trabecular thickness (Tb.Th) in the caudal vertebrae are indicated for comparison. e, Illustration of the generation of synthetic datasets of 2D sections from 3D micro-CT images along the longitudinal orientation. f, Convergence success rates (%) across all samples analysed, binned in intervals of 150 µm and 15° of translational and rotational initialisation offsets to the ground-truth (GT) transformation. The background colours of each bin indicate the simplest combination of image similarity functions that achieved the score indicated (remaining combinations not shown). Similarity functions: surface Dice similarity coefficient (DS), normalised Mutual Information (nMI), Pearson correlation coefficient (Pe), normalised Root Mean Squared Error (nRMSE), the nMI, nRMSE and Pe between the Euclidean distance transformed images (EDTnMI, EDTnRMSE and EDTPe, respectively). g, Convergence success rates (%) for the subset of sections initialised with a maximum of 150 µm translational offset to the GT transformation. Scores are binned in intervals of 15° of rotational initialisation offsets to the GT transformation in the y-axis, and with the normalised bone volume fraction (BV/TV) across the samples considered in the x-axis (one sample per column). Colour code of each bin as in sub-panel f). h, Convergence success rates (%) for sections initialised with a maximum of 15° rotational offset to the ground-truth transformation. Scores are binned in intervals of 150 µm translational offset to the ground-truth transformation in the y-axis, and with the normalised BV/TV across the samples considered in the x-axis (one sample per column). Colour code of each bin as in sub-panel f). i, Average mean absolute error (MAE) in effective strain (µ_ε_) values for successfully registered sections generated with rigid transformations (presented in f, g, h), plotted along the rotational and translational offsets of the final transformation. Dashed lines indicate contours delimiting areas with MAE below 1, 2 and 3 µε. Histograms at the top and right of the heatmap show the frequency of values in the heatmap (x10^3^). j, Convergence success rates (%) across varying deformation settings (σ) and rotational offsets, for sections initialised with a translation within 150 μm of the target location. Performance is optimal (96-100%) at low deformation (σ ≤ 0.5) and low rotation angles (0-15°). Success rates decline systematically with increasing deformation and rotation, reaching minimal values (2-9%) at high deformation (σ = 5) across all rotation angles. Colour code of each bin as in sub-panel f). k, Convergence success rates (%) across varying deformation settings (σ) and rotational offsets, for sections initialised within 15° of the target orientation. Colour code of each bin as in f. l, Average MAE in effective strain (µε) values for successfully registered sections generated with rigid transformations followed by elastic deformations (presented in j, k), plotted along the rotational and translational offsets of the final transformation. Dashed lines indicate contours delimiting areas with MAE below 1, 2 and 3 µε. Histograms at the top and right of the heatmap show the frequency of values in the heatmap (x10^3^).

To quantify the algorithm precision, we re-registered sections five times and quantified parameter dispersion using the median absolute deviation (MAD). For the caudal vertebra dataset (used for manual landmarking, *n* = 6), the mean ± standard deviation MAD was 1.27 ± 0.77 µm for translations, 0.19 ± 0.10° for rotations, and 0.10 ± 0.05% (X-axis) and 0.04 ± 0.02% (Y-axis) for uniform stretching. In contrast, for the mouse femur osteotomy dataset (*n* = 14), precision decreased to 14.14 ± 8.04 µm for translations, 2.34 ± 1.02° for rotations, and 0.99 ± 0.81% (X-axis) and 0.46 ± 0.39% (Y-axis) for 2D uniform stretching parameters. Reduced precision in the femur dataset likely reflects challenges posed by section deformations during registration.

Next, we assessed the convergence of the registration using simulated data. We generated datasets of 2D sections from 3D micro-CT images of mouse femora and 6^th^ caudal vertebrae of varying bone volume fraction (BV/TV), mimicking typical histological sections (Fig. 2e, Extended Fig. 1b, Supplementary Fig. 2, Supplementary Note 2). For each section, sequential trials were initialised between the default initialisation location and the final transformation, and with different combinations of image similarity functions guiding the registration step (see Methods and Supplementary Note 2). Our analysis of femur samples showed average convergence rates above 91% for sections initialised within 30° of the correct orientation or up to 300 µm of the target location, but not both simultaneously (Fig. 2f). We observed excellent convergence (96-100%) at low rotational deviations from the ground-truth (0-30°), for sections initialised within 150 µm of their target. Performance gradually deteriorated with higher angles, particularly in the 75-90° range, where success rates dropped to 14-27% (Fig. 2g). Perfect convergence (100%) was achieved at minimal translations (0-150 μm) across all BV/TV values. Success rates gradually declined as translational offsets increased, reaching 26-38% at the maximum considered (600-750 μm; Fig. 2h). Pairs of image similarity metrics based on the functions normalised Mutual Information (nMI), Surface Dice coefficient (DS), nMI and Pearson correlation between the Euclidean distance transformed images (EDTnMI and EDTPe) were sufficient to perform well within 150 μm translational offsets and 60° rotational offsets, whereas more complex combinations were required to maximise convergence rates for sections initialised further from their target or for samples with lower BV/TV (Fig. 2g-h). Either nMI or EDTnMI were consistently present in the best performing combinations under the most challenging initialisation conditions (Fig. 2f-h). BV/TV appeared to have a secondary influence on performance compared to the geometric transformations, though its effect was more pronounced under challenging initialisation conditions (higher rotational or translational offsets).

To put these results in perspective, we considered a histological dataset of a mouse femur (see Methods) and compared the initial guesses selected for each section with the final transformations after 2D-3D registration. We obtained a translational offset of 52.6 (32.6, 80.4) µm, and a rotational offset of 5.7° (3.8°, 9.5°); data indicated as median and interquartile range. These results are consistent with the excellent convergence observed in our in silico study, solidifying the performance of our 2D-3D registration approach.

Registration sensitivity to detect biologically relevant information was assessed with the mean absolute error (MAE) between mechanical signal values obtained with micro-FE as effective strain of the ground-truth and registered sections, across all successful attempts (Fig. 2i). This analysis achieved MAE values below 5 µε which is, at least, an order of magnitude smaller than typical values reported when quantifying bone mechanoregulation parameters^20^.

To further emulate realistic histological sections, we generated datasets with increasing magnitudes of non-linear deformations (Fig. 2d, Supplementary Fig. 2, Supplementary Note 2). This analysis followed the previous results for sections obtained with rigid transformations, with high convergence success for rotational deviations up to 30° and translational offsets up to 150 µm. Deformations with an average displacement of 40 µm (deformation sigma 3, Extended Fig. 1a) marked a transition to limited convergence success for cross-sectional and longitudinal orientations (Fig. 2j-k, Extended Fig. 1g-h). The MAE associated with effective strain values collected for successful attempts was below 50 µε (Fig. 2l, Extended Fig. 1i).

### L*iv*E profiling of bone cells during bone adaptation and regeneration

We applied Spatial μProBe to a femoral osteotomy sample from a previous fracture healing study^25^ and characterised the L*iv*E profile of osteocytes, the most abundant cells in bone. The dataset comprised weekly time-lapsed micro-CT images across seven weeks, and twelve serial histological sections produced at the end time-point stained for Safranin-O, receptor activator of nuclear factor kappa-Β ligand (RANKL) and Sclerostin. Bone tissue in each section was segmented using a human-in-a-loop approach (Supplementary Note 1). One operator (F.C.M.) registered the sections to the end time-point micro-CT image (Fig. 3a), achieving an average Dice score of 0.95 ± 0.02, which indicates a successful outcome consistent with high registration quality previously reported^32^. Using micro-FE analysis, we computed and mapped the 3D in vivo mechanical environment for each registered section (Fig. 3b). Distributions were statistically different (p<0.001, Conover test), except between sections 4 and 6. Individual cells in 2D histological sections were located in their L*iv*E environment (Fig. 3c), integrating cell staining information with the bone remodelling history and local mechanical signals, which defined statistically different L*iv*E profiles (Fig. 3d). These were consistent with the bone healing phases, with median mechanical signals increasing until weeks 3-4, when bridging occurred, followed by a reduction due to excess callus material that will be remodelled towards the original architecture.

**Figure 3.**
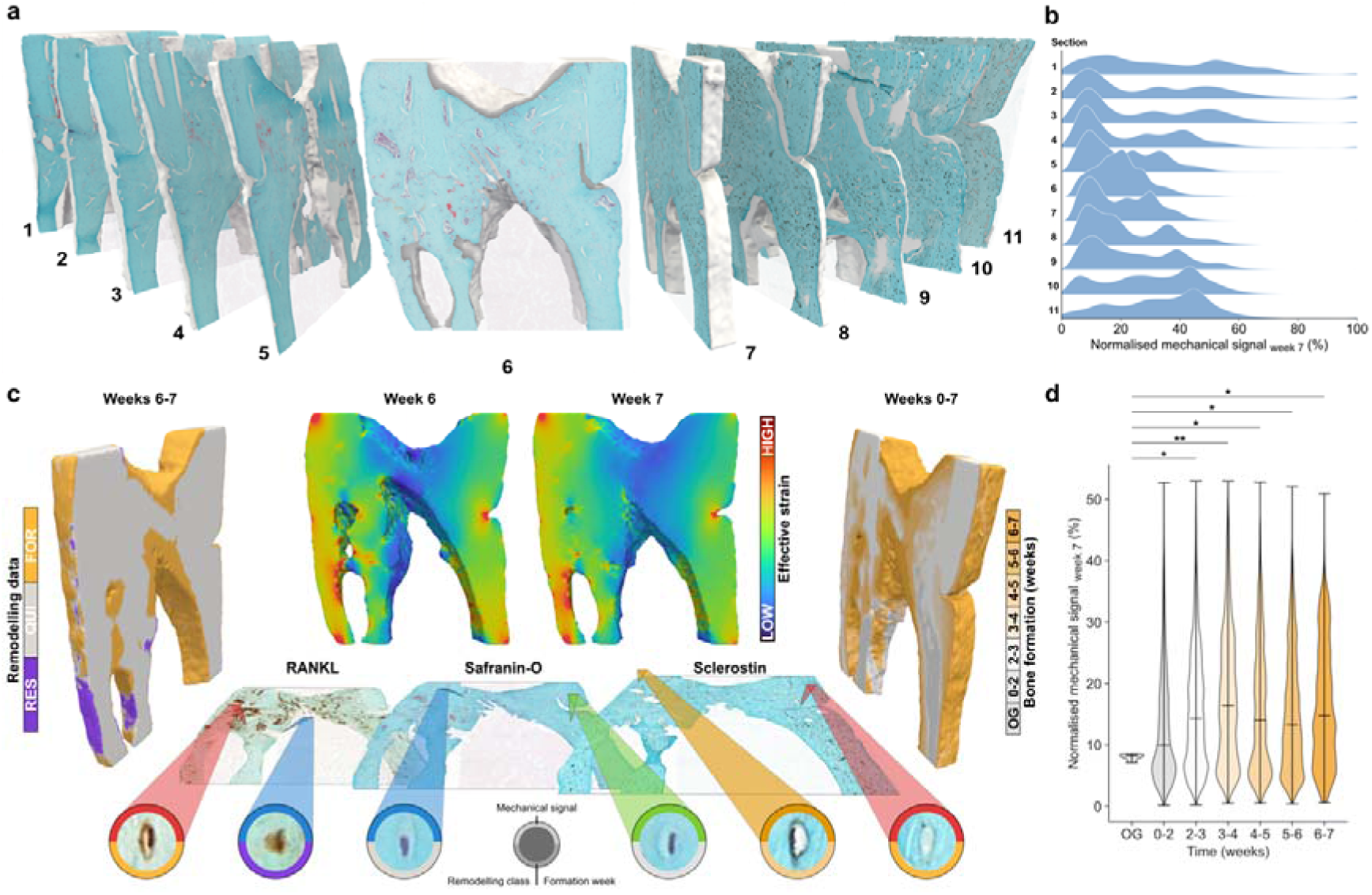
L*iv*E profiling of bone cells during bone regeneration. a, Example of consecutive 2D histological sections registered to the original 3D micro-computed tomography (micro-CT) image. Spatial μProBe successfully recovers the position of the histological sections in the micro-CT scan, maximising the amount of data extracted from each sample. 2D histological sections can be stained with different protocols (1-6: Safranin-O, 7-11: Sclerostin). b, Distributions computed with kernel density estimation (KDE) of local mechanical environments obtained from micro-finite element (micro-FE) analysis and extracted for each 2D section after 2D-3D registration. Due to heterogeneous mineralisation states and callus structure, each 2D section shows a different coverage of the full range of 3D mechanical signals. This effect can be offset with a holistic analysis of the sections. c, Spatial μProBe enables positioning individual cells in their local mechanical in vivo environment (L*iv*E) profile. Consecutive 2D histological sections are stained for relevant markers (e.g., receptor activator of nuclear factor kappa-Β ligand: RANKL, Safranin-O, Sclerostin), and individual cells segmented (e.g., manually or using deep learning segmentation models). After 2D-3D registration, each cell identified can be linked to their L*iv*E profile from 3D datasets (e.g., mechanical signal from micro-FE, bone remodelling or formation data from time-lapsed micro-CT images) and categorised accordingly. d, Distributions of local effective strain computed at week 5 with micro-FE normalised to the 99^th^ percentile of the 3D mechanical environment for bone formation regions identified with time-lapsed micro-CT for each interval of the study, highlighting varying L*iv*E profiles for cells in each week interval. Statistical comparisons shown only for comparisons between bone present originally at the start of the study (OG) and bone formed at subsequent intervals (**, p<0.01). Remaining comparisons among newly formed bone are all statistically significant (***, p<0.001 for comparisons between weeks 0-2 to 6-7 and **, p<0.01, for the comparison between weeks 3-4 and 5-6). Comparisons performed with post hoc pairwise multiple comparisons Conover’s test, using the Holm–Bonferroni method for adjusting p-values.

Nonetheless, bone mechanobiology stems from the interaction between several cell types across multiple systems (e.g., musculoskeletal, vascular, immunological). While Spatial μProBe can describe 3D L*iv*E profiles of individual cells at scale, established formalin-fixed paraffin-based (FFPE) techniques have limited capacity to stain markers specific to several cell types on the same section simultaneously. To counter this, we considered an ST dataset of another study evaluating bone healing of a femoral osteotomy^21^, under physiological and individualised cyclic loading^33^ and applied Starfysh^34^ to perform cell-type deconvolution of 15 cell types involved in bone regeneration (Supplementary data 1). One ST section per group (Control and Loaded) was 2D-3D registered with its end-point micro-CT, mapping mechanical signals from micro-FE to each image (Fig. 4a, Extended Fig. 2c). Based on time-lapsed in vivo micro-CT, voxels were categorised based on the week in which they were formed, grouping co-registered ST spots across time (Fig. 4b, Extended Fig. 2d). Cell-type relative proportions were super-resolved with iStar^30^ (Supplementary Note 3). Osteoblasts, osteoclasts, and mesenchymal stem cells, which are essential in bone mechanobiology, showed the highest relative proportions (Fig. 4c, Extended Fig. 2e, Supplementary Fig. 3a-b). This approach advances bone mechanoregulation analysis by leveraging generalised additive models (GAMs) to model continuous cell-type specific curves (Fig. 4d). Osteoclasts showed a baseline proportion of around 5% except in regions of low mechanical signals (< 10%) where it increased to over 10% (smooth term p-value < 0.001), coinciding with the interval where resorption was more frequently observed (Extended Fig. 2a-b, Supplementary Fig. 3c-d). Differences between relative proportions of osteoclastic and osteoblastic cell-markers were marginal under physiological loading but accentuated after cyclic mechanical loading (Fig. 4d-e, Extended Fig. 2f-g), with an increased proportion of osteoblastic cell-markers. The relative proportion of mesenchymal stem cells (MSCs) was also decreased in the latter loading scenario, concordant with the increased osteoblast differentiation induced by additional mechanical stimuli previously observed^35^. Osteocytic cell-markers were elevated in comparison to other cell-types (15% vs 5% for MSCs and osteoclasts, for example) under cyclic mechanical loading conditions, indicative of the increased mechanotransduction activity of these cells^36^. Across time, osteocytic markers under cyclic loading showed a sharp peak in bone formed after 2 weeks, when the callus was formed but not yet fully mineralised. This aligns with the need to direct osteoblasts and osteoclasts to deposit new bone and begin remodelling excessive material after bridging^37^. Osteoblastic markers were elevated (> 20%) in regions of higher mechanical signals (30% to 60%), where formation events were more frequently observed (Extended Fig. 2, smooth term p-value < 0.001). Interestingly, the relative proportion of osteoblastic markers was also elevated in regions of very low mechanical signals (< 15%), which coincided with regions where new bone formed and, therefore, was still mineralising (Fig. 4b).

**Figure 4.**
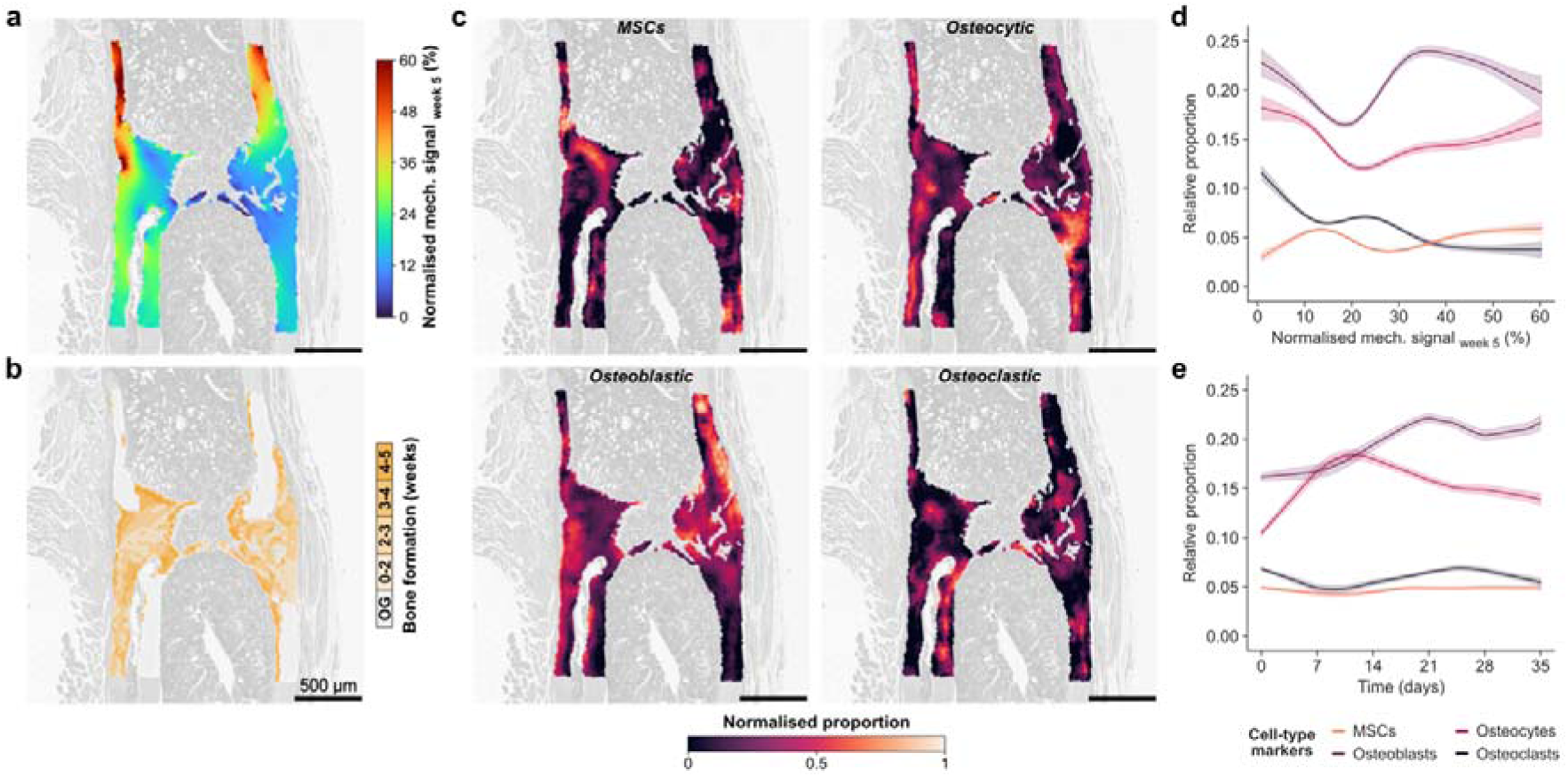
Cell-type deconvolution enables cell-type specific mechanoregulation analysis. a, Mechanical environment computed with micro-finite element (micro-FE) analysis on the micro-computed tomography (micro-CT) image and mapped to the 2D-3D registered spatial transcriptomics (ST) section of the Loaded sample. Normalised mechanical signal determined as effective strain values at week 5 and normalised to the 99^th^ percentile of the 3D mechanical environment. b, Bone formation regions identified with time-lapsed in vivo micro-CT and mapped to the 2D-3D registered ST section shown in figure a. “OG” describes the host bone, present at the start of the study. c, Selected examples of cell-type proportions determined with Starfysh^34^ and super-resolved with iStar^30^: mesenchymal stem cells (MSCs), osteocytic, osteoblastic and osteoclastic cells. Relative proportion values were normalised to the range 0-1 to support the visual comparison of individual cell-type distributions. d, Cell-level mechanoregulation analysis through the association between the relative proportion of the cell types presented in c with the mechanical environment shown in a, indicating non-monotonic changes in proportion across mechanical signals. Curves show the prediction obtained with generalised linear models (GAMs) with 25 splines and 5 effective degrees of freedom, and their 95% confidence interval. e, Association between the relative proportion presented in c with the time-points of bone formation shown in b, indicating differential distribution of cell populations across weeks. Curves obtained with GAMs with the same settings as in d.

### Time-lapsed differential gene expression analysis recapitulates the anabolic effect of mechanical loading

Individualised cyclic mechanical loading has been shown to accelerate the formation and mineralisation of the callus during healing^25^. We asked whether this time difference is detectable at the gene level and hypothesised that there would be a higher number of differentially expressed genes (DEGs) for comparisons between spots corresponding to bone formed in the same week than for pairings with positive time-offsets (i.e., bone formed first) for the Loaded sample. We compared the log2-fold change in gene expression between Control and Loaded samples, identifying seven clusters, including early upregulation, late upregulation or late downregulation (Supplementary Fig. 4a). The largest cluster of genes showed no change between conditions, indicating that only a limited subset is involved in mechanically-regulated responses. We first compared regions where bone was formed in the first three weeks and the last two weeks (i.e., before and after individualised mechanical loading) (Fig. 5a-c, Supplementary Fig. 4b-c). The comparison with the most differences occurred between bone formed in weeks 3-5 (Loaded sample) and early formation sites from the Control sample (weeks 0-3) (Fig. 5a). Key markers of bone formation such as *Col1a1*, *Col1a2* and *Bglap*, were strongly upregulated, while markers involved in endochondral ossification and mineralisation, like *Ccn2* and *Mgp*, were downregulated (Fig. 5b). The former set of genes remained upregulated for the comparison between weeks 3-5 for both Loaded and Control samples (Fig. 5c). We further improved the resolution of time comparisons to a nearly bi-weekly interval (Fig. 5d). In this setting, we observed a decrease in the number of DEGs between Control and Loaded samples with a positive offset (Fig. 5e), supporting our initial hypothesis, with the minimum being found for an offset of approximately 3 weeks. Expectedly, bone formation markers showed widespread expression in the Loaded sample, while markers involved in specialised functions showed more localised expressions, such as *Gja1* (essential for gap junctions), *Mmp9* (involved in bone matrix degradation) and *Cd34* (marker for hematopoietic stem cells; Fig. 5f, Supplementary Fig. 4d). These observations were also visible during bone adaptation in the mouse loading model of the 6^th^ caudal vertebra (Supplementary Fig. 6a-b). We identified 447 DEGs between regions of bone present at the start of the experiment for the Control and Loaded conditions, suggesting tissue-wide changes in gene expression with cyclic mechanical loading. In comparison, only 92 DEGs were observed between conditions for sites of bone formed during the experiment, where a limited number of markers associated with bone formation and mineralisation were upregulated (Supplementary Fig. 6c-e).

**Figure 5.**
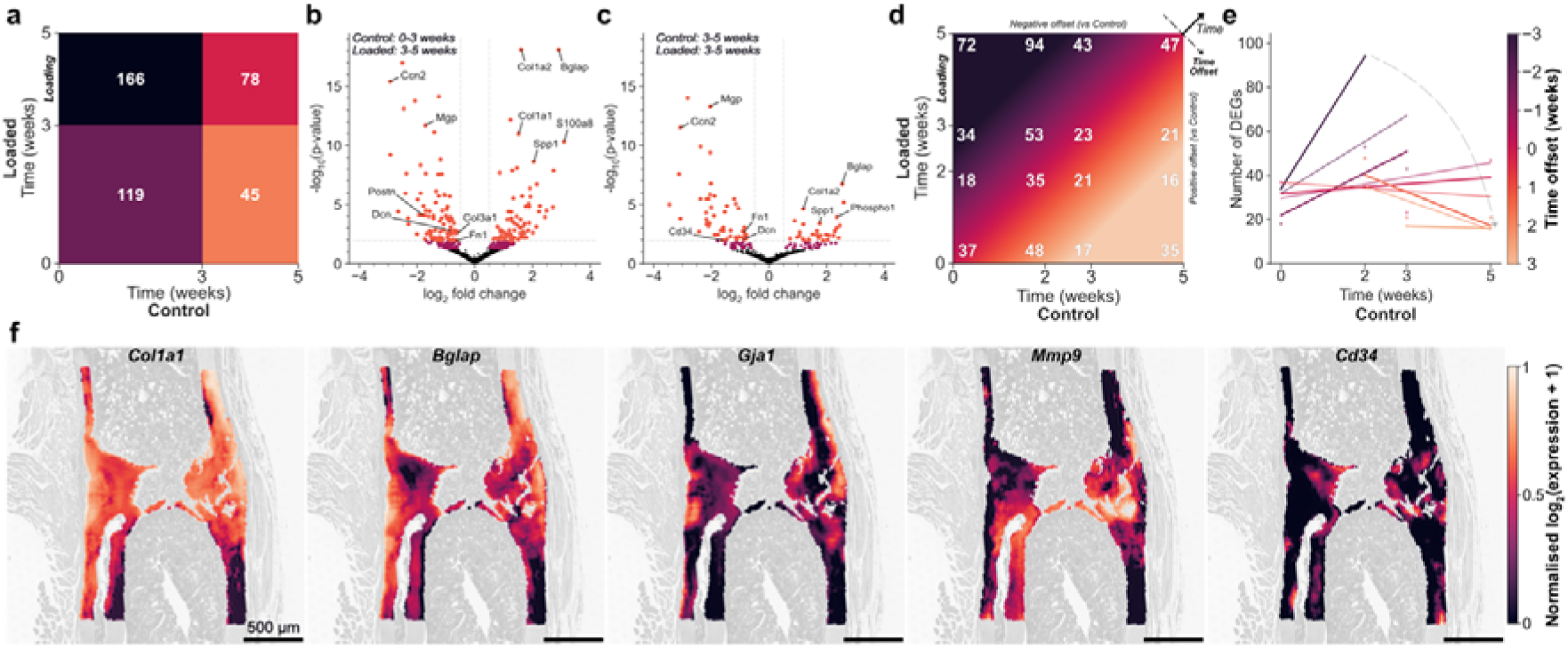
Differential gene expression analysis during bone fracture healing. a, Number of differentially expressed genes (DEGs) for pairwise comparisons between weeks 0-3 and 3-5 for Control and Loaded samples. Weekly individualised mechanical loading was applied to the Loaded sample between weeks 3-5. b, Volcano plot to visualize DEGs for the comparison between weeks 0-3 of the Control sample with weeks 3-5 of the Loaded sample. Significance criteria: absolute log2 fold change > 0.5 (vertical dashed lines); false discovery rate (FDR)-adjusted p-value < 0.0125 indicated in orange (after adjusting for multiple comparisons, horizontal dashed line) and FDR-adjusted p-value < 0.05 indicated in dark pink. Selected DEGs are annotated. c, Volcano plot to visualize DEGs for the comparison between weeks 3-5 of the Control sample with weeks 3-5 of the Loaded sample. Significance criteria and labelling follows the same rules indicated in b. d, Number of DEGs for pairwise comparisons between Control and Loaded samples for the spots in regions containing original bone (week 0), bone formed in weeks 0-2, bone formed in weeks 2-3 and bone formed in weeks 3-5. e, Linear modelling of the values presented in d. Values along the diagonals of d, corresponding to different time offsets between Control and Loaded samples, were plotted and fitted with linear models. Thick lines represent the values plotted in d, while thin lines denote the arithmetic mean of the values of consecutive time offsets, providing a visual representation of the intermediate data distribution between two offset values. A curved dashed arrow indicates the orientation of the time-offset axis labelled in d and in the colorbar. f, Super-resolved spatial gene expression maps of selected biologically-relevant markers at the fracture site of the Loaded sample. Osteoblast markers: *Col1a1* and *Bglap*; Gap junction marker: *Gja1*; Osteoclast marker: *Mmp9*; Hematopoietic stem cell marker: *Cd34*.

### Spatial μProBe enables time-lapsed gene mechanoregulation analysis during bone adaptation and regeneration

The Mechanostat Theory^18^ is a long-standing concept in bone mechanobiology, associating bone adaptation with tissue strains. This theory has been examined at the tissue-level^20^, revealing stable monotonic associations of bone surface velocity with L*iv*E mechanical signals. However, little is known about the (sub-)cellular responses that initiate tissue-level events. Therefore, we asked whether these monotonic responses hold at the gene level by determining continuous mechanoregulation curves for individual genes and hypothesised that gene expression is mechanoregulated during fracture healing, responding dynamically to mechanical cues over time.

First, we identified spatially variable genes (SVGs), as a proxy for genes more likely to vary with local mechanics. This revealed 580 and 742 SVGs for the Control and Loaded samples (Fig. 6a), respectively, with an overlap of 253 genes. However, only 208 genes had a mean count above 1 in the Loaded sample (in comparison to 263 in the Control sample), further supporting a selective refinement of gene expression with cyclic mechanical loading. During bone adaptation, this difference was even more accentuated, with the number of SVGs between cyclic and physiological loading decreasing from 706 to 72 (Supplementary Fig. 6f).

**Figure 6.**
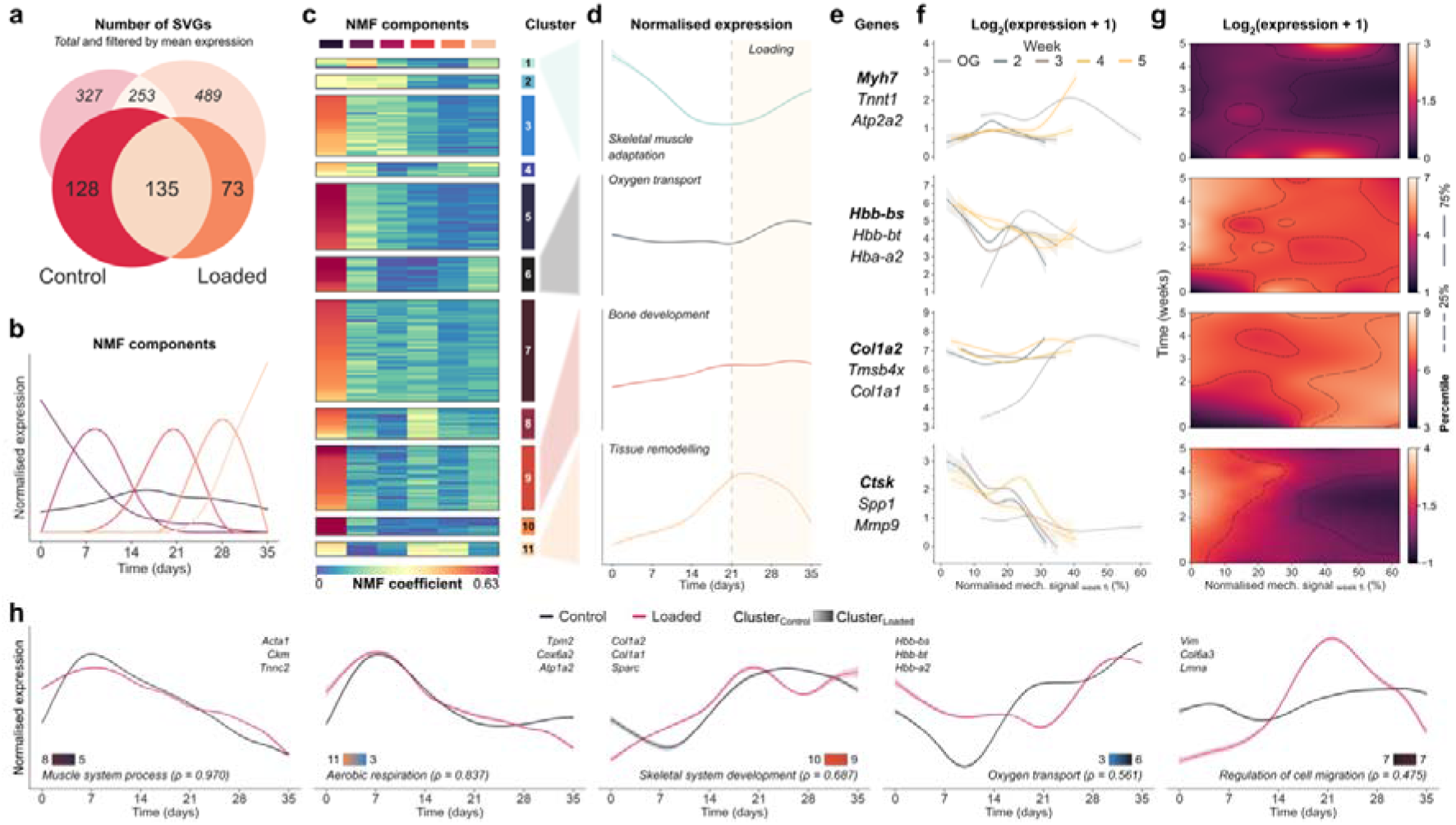
Time-lapsed and functional gene mechanoregulation analysis during bone fracture healing in a mechanically controlled environment. a, Number of spatially variable genes (SVGs) detected using SPARK-X^56^ for Control and Loaded samples. Only spots in bone tissue overlapping with the micro-computed tomography (micro-CT) registered image were considered. Values indicate the total number of genes detected (*italic*) and the number of genes with a mean unique molecular identifier (UMI) value per spot above 1 (UMI counts corrected with SCTransform^57^). Only filtered genes were considered in subsequent steps. b, Components obtained after non-negative matrix factorisation (NMF) analysis of the generalised additive model (GAM) curves associating gene expression and time for SVGs of the Loaded sample. GAM curves were normalised prior to NMF decomposition, to neutralise differences in expression magnitude between genes. c, Agglomerative clustering of all genes based on NMF coefficients. Clustering settings and results were optimised to maximise the silhouette score. d, Normalised expression for selected clusters reconstructed from NMF coefficients and components. Curves show the average expression and 95% confidence interval, obtained with bootstrapping (2500 iterations). Each curve is labelled with a functionally-relevant significant enrichment term identified with g:Profiler^58^ using the genes of the cluster, revealing how the expression of genes associated with those biological functions evolve over the 5-week experiment. e, Top 3 most expressed genes for the clusters represented in d. f, Time-lapsed gene mechanoregulation analysis for the genes highlighted in e in bold. A GAM (25 splines and 5 effective degrees of freedom) relating gene expression and local mechanical signals is computed for the set of points assigned to each time-point based on bone formation data obtained from time-lapsed micro-CT images after 2D-3D registration. Curves show the GAM prediction and 95% confidence interval. g, Time-lapsed gene mechanoregulation analysis for the genes highlighted in e in bold. A 2D GAM (25 splines and 25 effective degrees of freedom) is computed for all time-points simultaneously, yielding a continuous model of gene mechanoregulation across time and local mechanical signals. h, Comparison between the normalised expression of sets of genes between Control and Loaded samples. Gene sets were identified by comparing groups of genes (min. 3 elements) in clusters of one sample that were also aggregated in a cluster on the other sample. The top three most expressed genes are listed, as well as the cluster numbers (referring to c for the Loaded sample and Extended Fig. 3 for the Control sample). A functionally-relevant significant enrichment term identified with g:Profiler^58^ is indicated, determined using only the genes from each gene set. The Spearman correlation coefficient between the curves obtained for both samples is indicated as a proxy for the change in expression trends resulting from cyclic mechanical loading.

We assumed that SVGs were driving key physiological functions during healing, with genes involved in the same pathways showing similar responses over time. We modelled the expression of each SVG across time using GAMs and decomposed the predicted curves using non-negative matrix factorisation (NMF) into a subset of 6 components, while retaining over 99% of the variance in the data (Fig. 6b, Supplementary Fig. 5a). NMF components resembled the activation of different phases over time (Fig. 6b). Likewise, factorisation of expression curves as a function of mechanical signals, revealed similar activation windows (Supplementary Fig. 6g), hinting at mechanically-bounded regions that maximise the expression of specific genes, similar to band-pass mechano-sensing phenomena observed at single-cell scales^19^. Following this idea, we used agglomerative clustering to identify groups of genes with similar activations based on the magnitude of the factors for each NMF component (Fig. 6c, Extended Fig. 3a, Supplementary Fig. 6h). We identified key pathways involved in bone regeneration for genes in selected clusters, such as skeletal muscle adaptation, oxygen transport, bone development and tissue remodelling (Fig. 6d-e, Extended Fig. 3b-c). Our findings were corroborated by shifts in expression from week 3, when cyclic mechanical loading was applied to the osteotomy of the Loaded sample. Callus remodelling starting at this time-point was superseded by additional bone formation, aligning with the anabolic effect of mechanical loading. Hence, to inspect time-lapsed gene mechanoregulation, we selected individual markers (Fig. 6e, Extended Fig. 3c) and computed a GAM per time-point available (Fig. 6f, Extended Fig. 3d), which can be extended to a continuous 2D GAM that predicts gene expression across time and mechanical signals (Fig. 6g). This analysis revealed non-monotonic gene expression patterns, which we can also model analytically (Extended Fig. 3d, Extended Fig. 4, Supplementary Note 4), providing valuable quantitative inputs for in silico multiscale models of bone adaptation^38^ and regeneration^39^.

At last, we investigated whether sets of functionally-relevant genes present in clusters of the Control sample were conserved upon mechanical loading and whether their activation changed over time. Notably, several biological functions driven by small sets of genes were preserved, but their expression was influenced by mechanics to different degrees (Fig. 6h, Supplementary Fig. 5). Muscle-related processes (e.g., aerobic respiration) were unaltered (Spearman 11 = 0.837), while functions directly linked to bone mechanobiology showed larger variations, including sustained oxygen transport (with an increase upon applying cyclic mechanical loading at week 3), or the suppression of remodelling activity (indicated by reduced cell migration at week 3). Additional functions related to signal transduction and regulation of structural molecules showed steady activation in the Control sample, but increased fluctuations with cyclic loading (Supplementary Fig. 5). During bone adaptation, gene mechanoregulation followed a similar behaviour with collagen-related processes, essential for bone formation, being moderately affected by mechanical loading, while muscle-related activity remained consistent in both samples (Supplementary Fig. 6i-j).

## Discussion

Here, we presented Spatial μProBe, a correlative multimodal imaging approach for spatial profiling of biological micro-environments. We applied our tool to investigate bone mechanobiology, by integrating end-point 2D histological and ST data with 3D time-lapsed in vivo micro-CT data, revealing the L*iv*E profile of thousands of cells.

Spatial omics approaches have revolutionised the analysis of biological systems. However, highly dynamic biological processes undergo different phases over time, underscoring the need for MI approaches that can also resolve temporal scales. Recent tools like spatial-Mux-seq^40^ and moscot^41^ are already pushing these boundaries further, enabling spatiotemporal reconstructions of cell differentiation trajectories in the brain, liver and pancreas. In bone, SpatialTime^42^ has been successfully applied to characterise the activation of signalling pathways in intact mouse femora, assuming a linear development across space as a proxy for time. While existing MI approaches focusing on bone tissue were limited to structural characterisation^43,44^, Spatial μProBe can leverage time-lapsed in vivo micro-CT to delimit spatially accurate regions for weekly timepoints, resolving spatial and temporal scales concurrently. This approach enabled the analysis of complex bone regeneration scenarios, where the spatial linearity assumption of SpatialTime is not valid because of the highly heterogenous callus structures.

Accordingly, local mechanics play a crucial role in determining cell differentiation^45^ through key mechanosensitive pathways such as YAP/TAZ^46,47^, which drive tissue formation and regeneration. Using micro-FE, we reconstructed the 3D L*iv*E profiles of bone cells, revealing the effects of mechanical loading on gene expression measured with ST data. Supported by super-resolved ST data, we derived precise curves describing gene expression changes over time, during key functional processes involved in bone adaptation and regeneration. Our analytical framework builds upon the tissue-level understanding of bone mechanoregulation, as proposed in the Mechanostat Theory^18^ and quantified with time-lapsed micro-CT^20,22,48^ and extends it to the cellular and molecular scales. By integrating spatial omics with the native mechanobiological environment, we were able to quantitatively describe the mechanical regulation of gene expression, and reveal non-linear associations involving multiple gene groups that collectively shape the tissue-level response, thereby challenging the monotonic proposal described by Frost and aligning with observations at the single-cell scale^19^. While recent advances in ST technologies, such as Visium HD, allow sub-cellular transcriptome profiling, the combination of the standard Visium kit with super-resolution offers an attractive cost-effective alternative to generate high resolution ST data across experimental conditions. Such data enables the unique integration of molecular information with high-resolution tissue properties, such as mechanical strain, which are also computed at the cell scale.

Several correlative imaging approaches have been proposed for 2D-3D registration^2,5^, with some specifically developed for bone tissue^31,43^ using commercial software or focusing on limited ROIs. Powerful image analysis packages like ITK^49^, ANT^50^ and elastix^51^ are also compatible with 2D-3D registration tasks but we found no performance advantages per optimisation iteration. These also required additional complexity to setup and handle image data within our workflow or produced invalid results due to non-linear transformations while processing 2D bone histological sections in a 3D space. Therefore, we opted for a straightforward implementation based on binary image registration with affine transformations that is fast, accurate and compatible with other combinations of imaging modalities where the tissue of interest can be reliably segmented. As shown in our large-scale in silico validation analysis, a plausible initial guess is crucial for reliably performing 2D-3D registration. We aid this task in real applications by leveraging dedicated 3D viewers, like ParaView^52^, which allow browsing and digitally sectioning 3D structures conveniently. Given recent advances in large scale models for image analysis, like MatchAnything^53^, we anticipate it will be possible to expedite and automate this process, further improving the throughput of image registration.

Likewise, our ST analysis was limited to one section per condition. While we aimed to optimise sample sectioning to maximise the coverage of distinct L*iv*E profiles, integrating consecutive sections would improve the quality of ST data analysis^54^ and solidify the identification of SVGs and their expression across local mechanical signals. As ST technologies become more accessible, Spatial μProBe provides the tools for serial registration of ST data, which will enable a comprehensive mapping of L*iv*E gene mechanoregulation across anatomical sites and healthy or aged conditions, during bone adaptation and regeneration.

Ultimately, Spatial μProBe can bridge the gap between mechanical signalling and molecular biology during complex temporal and spatial events. Within the context of bone mechanobiology, decoding cellular and molecular responses to multiscale events like mechanical loading is vital for a holistic understanding of bone mechanoregulation. As mechanical interventions in preclinical models have demonstrated significant potential in inducing anabolic responses^21,25^, this knowledge is a crucial next step in advancing fracture healing therapies and may have broader implications for other bone-related conditions. Critically, we anticipate that the potential of Spatial μProBe extends far beyond bone mechanobiology, with valuable applications in other fields where multiscale molecular pathways play a critical role.

---

**Extended Figure 1.**
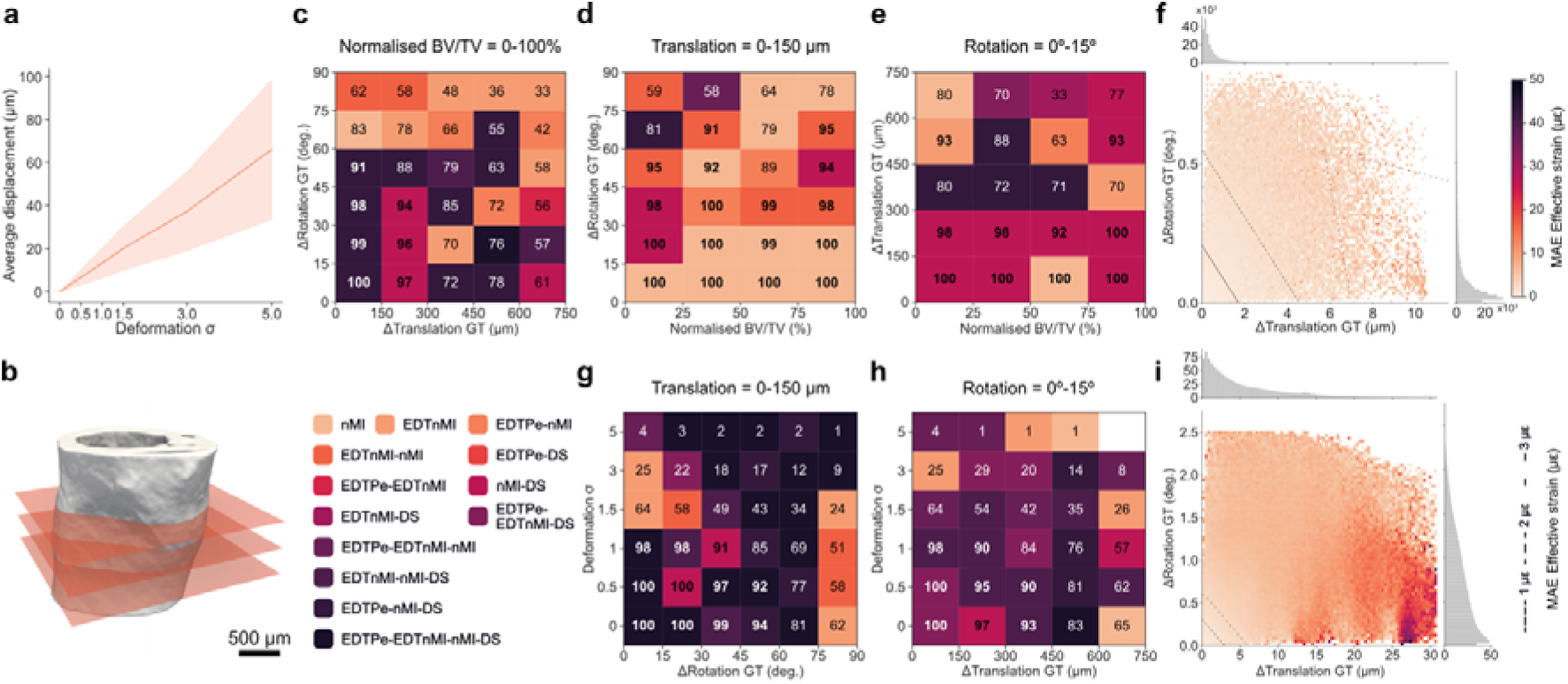
Validation of 2D-3D registration in Spatial μProBe for mouse femur sections obtained in cross-sectional orientation. a, Relationship between the deformation setting (σ) to simulate non-linear deformations on sections generated in silico and the average displacement (in µm) induced across the image. b, Illustration of the generation of synthetic datasets of 2D sections from 3D micro-computed tomography (micro-CT) images along the cross-sectional orientation. c, Convergence success rates (%) across all samples analysed (average of all values of bone volume fraction, BVTV), binned in intervals of 150 µm and 15° of translational and rotational initialisation offsets to the ground-truth (GT) transformation. The background colours of each bin indicate the simplest combination of image similarity functions that achieved the score indicated (remaining combinations not shown). d, Convergence success rates (%) for sections initialised with a maximum of 150 µm translational offset to the GT transformation. Scores are binned in intervals of 15° of rotational initialisation offsets to the GT transformation in the y-axis, and with the normalised BV/TV across the samples considered in the x-axis (one sample per column). Color code of each bin as in c. e, Convergence success rates (%) for sections initialised with a maximum of 15° rotational offset to the GT transformation. Scores are binned in intervals of 150 µm translational offset to the GT transformation in the y-axis, and with the normalised BV/TV across the samples considered in the x-axis (one sample per column). Color code of each bin as in c. f, Average mean absolute error (MAE) in effective strain values (µε) for successfully registered sections generated with rigid transformations (presented in f, g, h), plotted along the rotational and translational offsets of the final transformation. Dashed lines indicate contours delimiting areas with MAE below 1, 2 and 3 µε. Histograms at the top and right of the heatmap show the frequency of values in the heatmap (x10^3^). g, Convergence success rates (%) across varying deformation settings (σ) and rotational offsets, for sections initialised with a translation within 150 μm of the target location. Performance is optimal (96-100%) at low deformation (σ ≤ 0.5) and low rotation angles (0-15°). Success rates decline systematically with increasing deformation and rotation, reaching minimal values (2-9%) at high deformation (σ = 5) across all rotation angles. Color code of each bin as in c. h, Convergence success rates (%) across varying deformation settings (σ) and rotational offsets, for sections initialised within 15° of the target orientation. Color code of each bin as in f. i, Average MAE in effective strain (µε) values for successfully registered sections generated with rigid transformations followed by elastic deformations, plotted along the rotational and translational offsets of the final transformation. Dashed lines indicate contours delimiting areas with MAE below 1, 2 and 3 µε. Histograms at the top and right of the heatmap show the frequency of values in the heatmap (x10^3^).

**Extended Figure 2.**
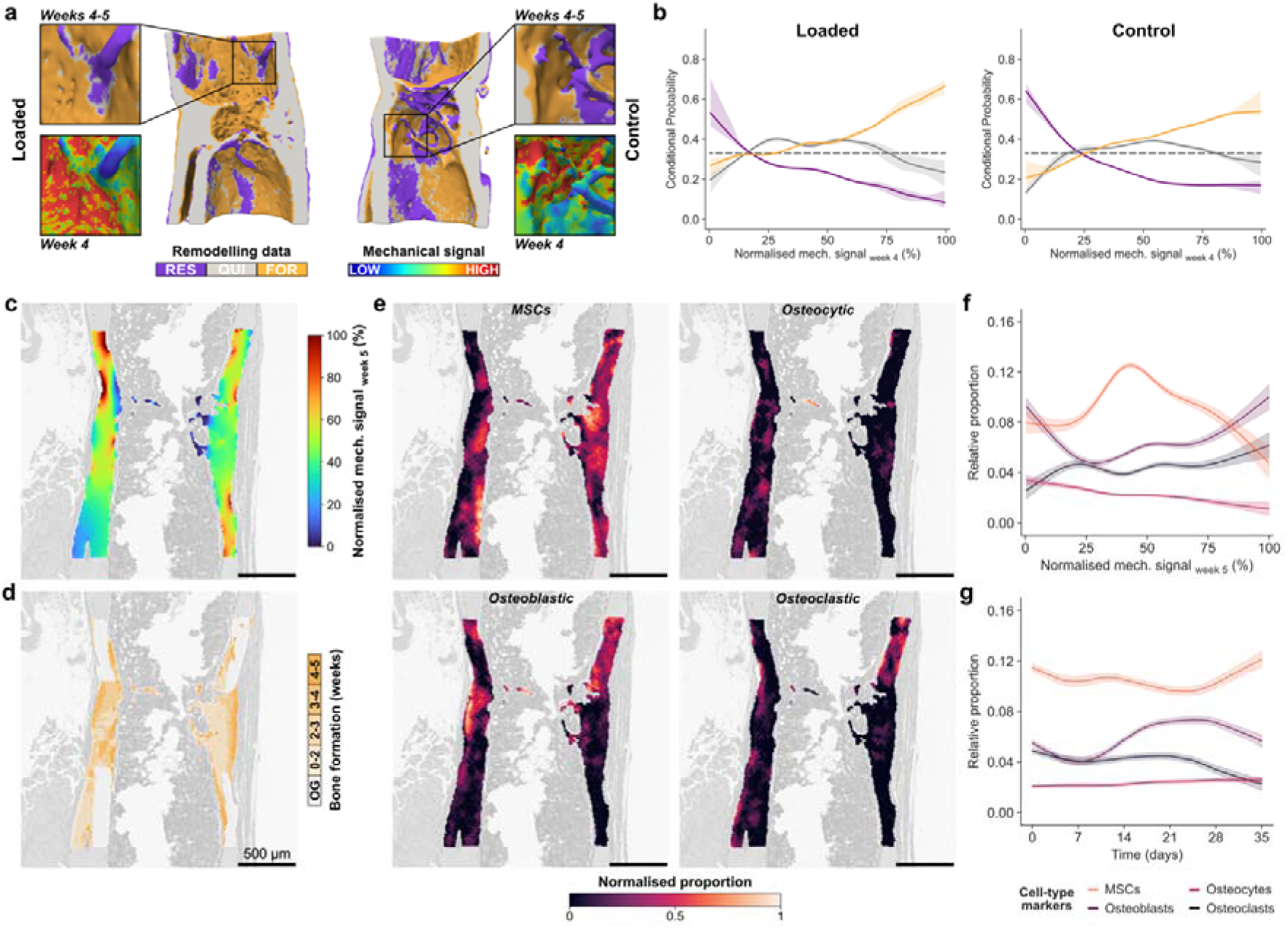
Bone mechanoregulation at the tissue and cell scales. a, Visualisation of (re)modelling events (formation: FOR, quiescence: QUI, resorption: RES) between weeks 4 and 5 determined from time-lapsed in vivo micro-computed tomography (micro-CT) images for the Control and Loaded samples. A selected region is highlighted, featuring the local mechanical signals computed with micro-finite element analysis (micro-FE), and indicating an association between high and low mechanical signals at week 4 with formation and resorption events over the subsequent week (weeks 4-5), respectively. b, Conditional probability curves connecting the surface mechanical environment as effective strain, normalised to the 99^th^ percentile, with surface (re)modelling events. The plots show the mean probability line and its corresponding 95% confidence interval, per (re)modelling event for the interval 4-5 weeks after bootstrapping (2500 iterations) of a LOWESS operation. The dashed line at 0.33 identifies the probability of a random event for a ternary classification case. c, Mechanical environment computed with micro-FE on the micro-CT image and mapped to the 2D-3D registered spatial transcriptomics (ST) section of the Control sample. Values are normalised to the 99^th^ percentile of the 3D mechanical environment. d, Bone formation regions identified with time-lapsed in vivo micro-CT and mapped to the 2D-3D registered ST section of the Control sample. “OG” describes the host bone, present at the start of the study. e, Selected examples of cell-type proportions determined with Starfysh^34^ and super-resolved with iStar^30^: mesenchymal stem cells (MSCs), osteocytic, osteoblastic and osteoclastic cells. Relative proportion values were normalised to the range 0-1 to support the visual comparison of individual cell-type distributions. f, Association between the relative proportion of the cell types presented in e with the mechanical environment shown in c, indicating non-monotonic changes in proportion across mechanical signals. Curves show the prediction obtained with generalised linear models (GAM) with 25 splines and 5 effective degrees of freedom, and their 95% confidence interval. g, Association between the relative proportion presented in e with the time-points of bone formation shown in b, indicating differential distribution of cell populations across weeks. Curves obtained with GAMs with the same settings as in f.

**Extended Figure 3.**
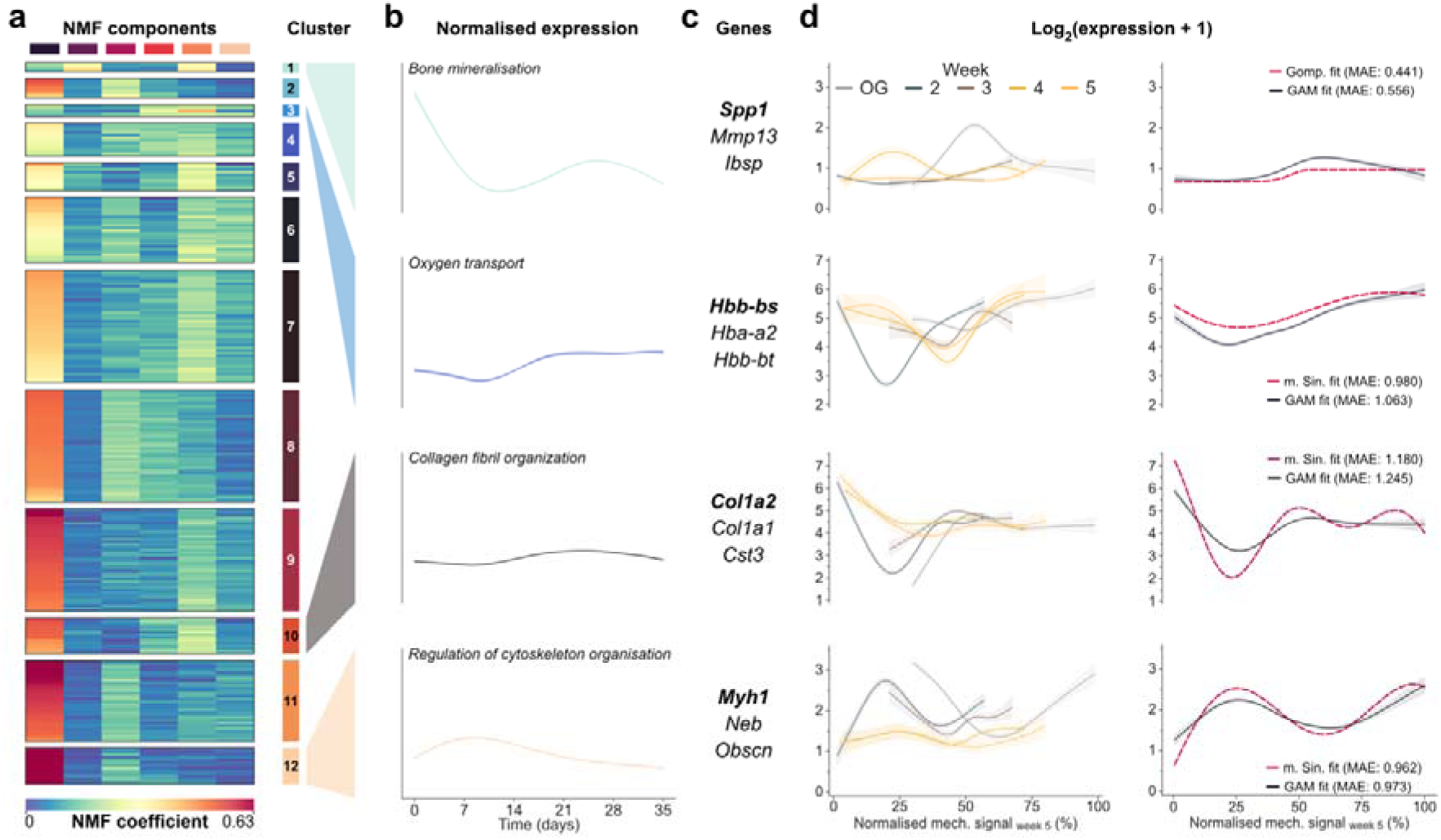
Time-lapsed and functional gene mechanoregulation analysis during bone fracture healing under physiological loading. a, Agglomerative clustering of spatially variable genes (SVGs) of the Control sample, based on non-negative matrix factorisation (NMF) coefficients after factorisation with the components displayed in Supplementary Fig. 5a. Clustering settings and results were optimised to maximise the silhouette score. b, Normalised expression for selected clusters reconstructed from NMF coefficients and components. Curves show the average expression and 95% confidence interval, obtained with bootstrapping (2500 iterations). Each curve is labelled with a functionally-relevant significant enrichment term identified with g:Profiler^58^ using the genes of the cluster, revealing how the expression of genes associated with those biological functions evolve over the duration of the experiment. c, Top 3 most expressed genes for the clusters represented in d. d, Time-lapsed gene mechanoregulation analysis for the genes highlighted in c in bold. A generalised additive model (GAM, 25 splines and 5 effective degrees of freedom) relating gene expression and local mechanical signals is computed for the set of points assigned to each time-point based on bone formation data obtained from time-lapsed micro-computed tomography (micro-CT) images after 2D-3D registration. Curves show the GAM prediction and its 95% confidence interval. The right column shows the resulting curve fits of mathematical functions to the same data, providing quantitative descriptors to model gene expression over time (see Supplementary Note 4). The mean absolute error (MAE) is indicated, for the fit of the function that best fitted the data and GAM curve.

**Extended Figure 4.**
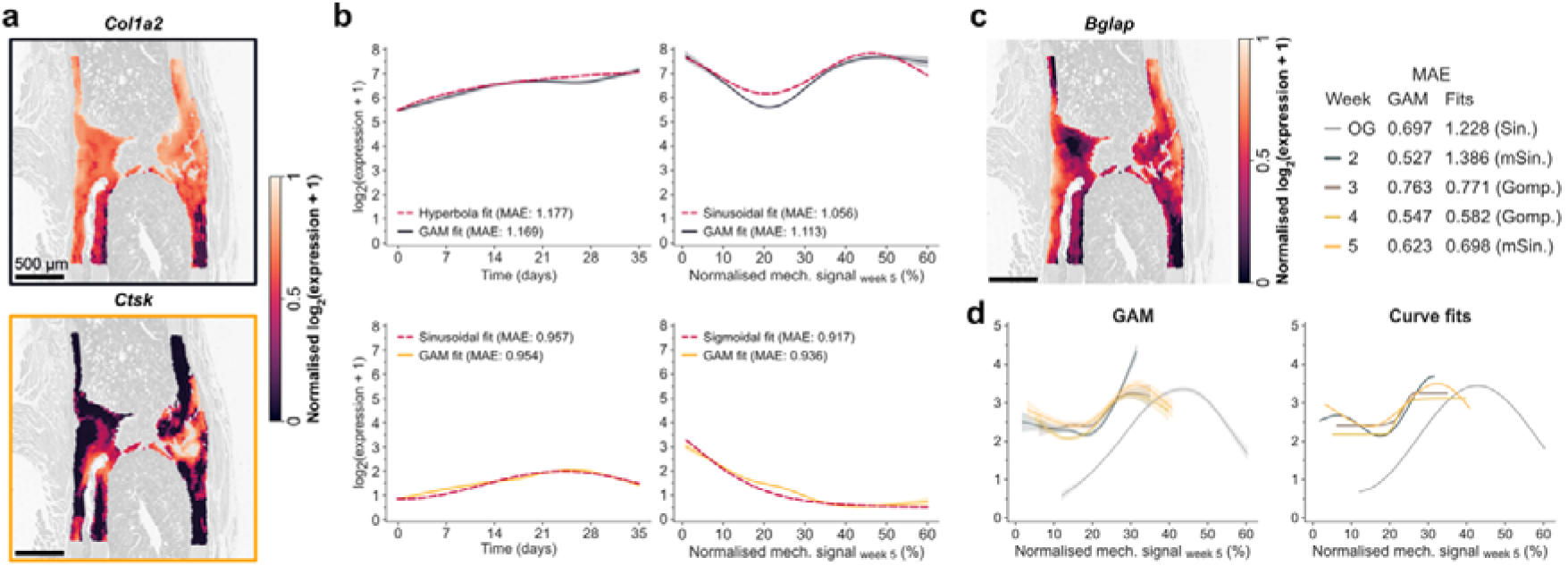
Analytical modelling of time-lapsed and functional gene mechanoregulation analysis during bone fracture healing. a, Super-resolved spatial gene expression maps of selected biologically-relevant markers at the fracture site of the Loaded sample: *Col1a1* and *Ctsk*. b, Curve fits of mathematical functions to the data associating gene expression data over time and local mechanical signals (see Supplementary Note 4). The top and bottom rows correspond to *Col1a1* and *Ctsk*, respectively. The mean absolute error (MAE) is indicated, for the curve fit of the function that best fitted the data and the generalised additive model (GAM) prediction. c, Super-resolved spatial gene expression map of *Bglap* at the fracture site of the Loaded sample. d, Curve fits of mathematical functions to the time-lapsed data associating gene expression data and local mechanical signals (see Supplementary Note 4). The left plot shows the GAM curve (25 splines and 5 effective degrees of freedom) relating gene expression and local mechanical signals computed for the set of points assigned to each time-point based on bone formation data obtained from time-lapsed micro-computed tomography (micro-CT) images after 2D-3D registration. Curves show the GAM prediction and its 95% confidence interval. The right plot shows the resulting curve fits of mathematical functions to the same data (see Supplementary Note 4). The MAE is indicated, for the fit of the function that best fitted the data and GAM curve.

## Methods

### Spatial μProBe workflow

Spatial μProBe was implemented in Python 3 and the graphical user interface was created with Solara (https://solara.dev/). A detailed documentation is available online. It is designed to be modular, allowing for easy integration of new imaging modalities and data analysis methods, while maintaining a user-friendly interface and an end-to-end workflow. There are three core steps in the primary workflow: image preprocessing, multimodal 2D-3D image registration and correlative analysis of multimodal data. Projects are organised in folders with a file tree structure that contains all the input data, associated metadata and output produced in the analysis.

### Image preprocessing

#### 2D images

Images are imported in their native format and their binary annotations in JSON format. Depending on the size of the dataset, users can prepare annotations manually or train and leverage deep-learning image segmentation models available in Spatial μProBe or elsewhere. For general tissue segmentation, we incorporated four well-established architectures available in Segmentation Models Pytorch (https://github.com/qubvel-org/segmentation_models.pytorch) into our workflow. Namely, we support UNet, UNet++, MANet, PAN and DeepLabV3+ architectures, which can be combined with a variety of backbones (VGG, ResNet, SE-ResNet, ResNeXt, SE-ResNeXt, SENet154, DenseNet, Inception, MobileNet, EfficientNet). Additionally, for specialised cell segmentation tasks, we also support models trained with Cellpose^27,28^. For convenience, we also provide a tool to convert binary annotations in image format into JSON format.

Two additional steps are implemented for performance considerations, with the aim of reducing the dimensions of the 2D image, which are typically histological or microscopy data imaged at very high resolution. First, images are rescaled to the lower resolution of the 3D image (e.g., micro-CT). Next, as 2D sections can cover much larger tissue areas than those available in the complementary modality, the user can define a region-of-interest (ROI) in each image focusing on the common region between the two imaging modalities, to be applied before the 2D-3D registration step.

#### 3D images

Spatial μProBe requires binary 3D images for 2D-3D image registration. We provide tools for importing, Gauss-filtering, component-labelling and applying thresholds to 3D images obtained in AIM format from Scanco micro-CT scanners (Scanco Medical AG, Brüttisellen, Switzerland). Users can also create compatible input files using the h5i module provided with Spatial μProBe.

### Multimodal 2D-3D image registration

A 2D-3D image registration step lies at the core of Spatial μProBe. Our approach relies on comparing the 2D binary image (based on the image annotation provided) with a 2D binary section obtained from the 3D image, for a given set of transformation parameters. The goal is to identify the 3D rigid transformation and 2D in-plane stretching parameters (along the vertical and horizontal axis) that produce the best overlap between these 2D images. The use of binary images makes the registration process independent of the modalities in use, and reduces the memory consumption of the image data, which is favourable for such complex optimisation tasks.

#### Initial guess estimation

First, 2D and 3D images are loaded from the corresponding pre-processed files. Pre-defined transformations around the axes of the 2D and 3D coordinate frames are available to facilitate a global alignment between both images. Next, users are expected to provide an initial guess for the transformation. To aid this operation, we provide tools to export the 3D binary image into a format compatible with ParaView 5.12^52^, which conveniently allows exploring and visualising the 3D image using the Section Tool to identify promising candidate parameters for further optimisation.

#### Iterative Assisted Registration

Next, we developed an approach entitled “Iterative Assisted Registration” (IterAR) that aims to facilitate the optimisation of the similarity between both binary images. This procedure searches the neighbourhood of the current transformation parameters for the values that maximise a set of similarity functions selected by the user, optimised using the Powell method. Currently, the following functions are supported: Dice similarity coefficient (Di), surface Dice similarity coefficient (DS), normalised Mutual Information (nMI), Pearson correlation coefficient (Pe), Mean Squared Error (MSE), the nMI and Pe between the Euclidean distance transformed images (EDTnMI and EDTPe, respectively). The range of the search neighbourhood can be adjusted to any value up to 50 units (no distinction is made in magnitude between degrees in rotation or voxels in translation). Users can select which combination of transformations (rotations, translations or uniform stretching) are allowed at each iteration and make any adjustments to the parameters before starting another iteration. Rotation angles correspond to Euler angles and refer to extrinsic rotations in the order “XYZ”. Through visual inspection of binary and grayscale image overlap, users can determine the end of the optimisation, supported by a notification indicating that the change in the optimisation metric displayed has not changed by more than 1% in the last five iterations.

### Correlative analysis

#### Fine-tuning with deformable 2D-2D image registration

To account for circumstances where uniform stretching cannot account for non-linear deformations in either registered modality, we provide a post-processing tool after 2D-3D registration for deformable image registration based on a BSpline transformation from SimpleITK^59^. This operation implies defining one of the modalities as the reference to which the other will be registered, and the density of the grid to be used in the optimisation, which determines the number of points available to deform the image to match the shape of the reference image.

#### Analysis of 2D histological sections

In Spatial μProBe, multimodal data evaluation can be performed in two ways: “dense” analysis of all pixels of the 2D image or “component-labelled”, focusing on individual elements (e.g., segmented cells) of the 2D image. Both options serve to determine the coordinates in 2D of the relevant pixels of the 2D image, to be used for multimodal integration into 3D space, with the latter also allowing the user to quantify 2D morphometric parameters of each component such as perimeter, area, minor and major axis length.

#### Analysis of Spatial Transcriptomics sections

For applications relying on ST data, Spatial μProBe is compatible with the data produced with Visium Spatial Gene Expression for FFPE assay (10x Genomics, Pleasanton, USA). Specifically, spot locations can be masked based on the tissue class annotations defined on the 2D histological section (including the class used for 2D-3D registration).

#### Correlative analysis of multimodal data

At last, Spatial μProBe provides convenient tools to correlate multimodal 2D and 3D datasets of the same sample. We provide three methods to achieve to perform this operation: “pixel-wise” fashion, where each pixel is linked to the value of the voxel where it is located in 3D; “radially”, where each pixel is described by measures of location (minimum and maximum, mean ± standard deviation, median and interquartile range) of the sphere of a user-defined radius centred in the voxel where the pixel is located in 3D; “surface-based”, where each pixel is linked to the value of the closest surface voxel of a given binary 3D mask, determined by the Euclidean distance. Outputs are exported in CSV format, in a table structured for post-processing analysis.

## Datasets

### Micro-computed tomography datasets

#### In vivo micro-CT imaging of the femur defect model

Micro-CT images of the mouse femur samples used for the ST analysis and in silico validation of the 2D-3D registration algorithm were obtained from two previous studies^21,25^. Briefly, the samples composing the ST dataset consisted of female 12-week-old bone cell reporter (BCR) mice (n = 2), while the images for the 2D-3D validation dataset were female 20-week-old C57BL/6J mice (n = 20). All mice received mid-diaphyseal femoral defects, supported with an external fixator (Mouse ExFix, RISystem, Davos, Switzerland) using four mounting pins. Three weeks after surgery, when defect sites typically bridge, the mice received either individualised cyclic mechanical loading via the external fixator (8 to 16 N, 10 Hz, 3000 cycles, 3x / week) or 0 N sham-loading. In vivo micro-CT imaging of the fracture site between the two inner screws of the external fixator was performed weekly for all mice. For the ST dataset, scans post-surgery were performed between weeks 0-5 (vivaCT 80, Scanco Medical AG, Brüttisellen, Switzerland; 10.5 μm resolution, 55 kVp, 145 µA, 350 ms integration time, 500 projections per 180°, 21 mm field of view (FOV), scan duration ca. 15 min), and for the 2D-3D validation dataset between weeks 0-7 (vivaCT 40, Scanco Medical AG, Brüttisellen, Switzerland; 10.5 μm; 2 stacks of 211 slices; 55 kVp, 145 μA, 350 ms integration time, 500 projections per 180°, 21 mm field of view (FOV), scan duration ca. 15 min). For the latter, all images from week 7 were sorted into groups based on the BV/TV percentiles (25^th^, 50^th^, 75^th^, 100^th^) of the dataset and one sample was randomly selected from each group, serving as an indirect proxy for the number of shape-specific image features (e.g. trabeculae) available. At 17 and 27 weeks, animals were sacrificed, and samples were processed to generate ST datasets and histological sections, respectively.

For each sample, reconstructed micro-CT images were registered sequentially^23^. Proximal and distal cortices were registered separately between consecutive time-points until bridging of the defect occurred. To ensure that complete cross sections of the femur were visible on proximal and distal sites across all time-points, registered scans were cropped to the largest consecutive interval of the 3D volume along the longitudinal axis determined across all time-lapsed images for each sample. Images were Gauss-filtered (sigma: 1.2, support: 1) and binary images were computed (threshold: 395 mgHA/cm^3^).

#### In vivo micro-CT imaging of the caudal vertebra model

Micro-CT images of the mouse vertebrae used for the ST analysis and in silico validation of the 2D-3D registration algorithm were obtained from a previous study^60^. Samples used for ST and for the 2D-3D validation dataset were BCR and C57BL/6J mice, respectively, as above for the femur model. Briefly, stainless steel pins were inserted into the fifth and seventh caudal vertebra of 12-week-old female mice (n = 37). After 3 weeks, mice received either sham (0 N, n = 18) or 8 N (n = 19) supra-physiological cyclic loading on their 6^th^ caudal vertebra with 10 Hz frequency for 5 min, three times per week over four weeks, with weekly in vivo micro-CT imaging of all mice using the vivaCT80 device (Scanco Medical AG) and the same settings described above. Reconstructed micro-CT images were registered consecutively^61^. Images were Gauss-filtered (sigma: 1.2, support: 1) and binary images were computed (threshold: 580 mgHA/cm^3^). Again, for the 2D-3D validation dataset, samples were sorted into groups based on the BV/TV percentiles (25^th^, 50^th^, 75^th^, 100^th^) of the dataset and one sample was randomly selected from each group.

#### Ex vivo micro-CT imaging of the caudal vertebra model

High-resolution micro-CT images used for the validation of the 2D-3D registration algorithm with manual landmarking were obtained from a previous study^62^. Briefly, 6^th^ caudal vertebrae (n = 3) were scanned at high-resolution, with a voxel-size of 2.1 μm (microCT50, Scanco Medical AG, Brüttisellen, Switzerland), and at 10.5 μm (microCT40, Scanco Medical AG, Brüttisellen, Switzerland), to simulate scans with in vivo quality. Scans at in vivo resolution were registered onto the corresponding high-resolution images. Images were Gauss-filtered (sigma: 1.2, support: 1) and binary images were computed (threshold: 580 mgHA/cm^3^).

### In silico datasets for 2D-3D registration validation

#### Datasets generated with rigid transformations

For each anatomical site (caudal vertebra and femur models), we used four micro-CT images (see section “Datasets”, regarding the sample selection process) and produced ten datasets per scan, five per orientation typically considered in histological sectioning (longitudinal and cross-sectional, producing sections parallel and perpendicular to the z-axis of the micro-CT images, respectively). For each dataset, an off-axis sectioning orientation was sampled based on user-defined ranges for translation, rotation and in-plane stretching values. Translation values were sampled uniformly within a range defined by 30 voxels into the limits of the bounding box of each micro-CT image, which approximately matches the usable range of histological sections produced from FFPE samples. Rotation values perpendicular to the z-axis (phi) could take any value between 0-180°, a rotation with respect to the z-axis (theta) could range between −15° to 15° for longitudinal datasets and 75°-105° for cross-sectional datasets, mimicking small variations in the sectioning angle resulting from different sample embedding orientations. For each dataset, 25 sections were extracted from the binary micro-CT image along the relevant axis (z-axis for cross-sectional, x-axis for longitudinal), using affine transforms of the sampled transformations and linear interpolation of the values on the 3D image.

#### Datasets generated with elastic transformations

To mimic typical structural deformations in histological sections, we produced additional datasets following the same workflow above that included an additional step that applied an elastic deformation with varying magnitude. Again, we generated ten datasets per scan, five per orientation (longitudinal and cross-sectional). Elastic deformations were created with a 2D deformation grid, defined by N x N points, where each point was randomly displaced by a value determined from a Gaussian distribution with a given sigma. Displacements were interpolated for all pixels of the image, producing a deformed image. Based on the pairwise distances between landmarks obtained previously, we performed a parametric study to evaluate which combination of Gaussian sigma and grid density values best matched these distributions. Specifically, we selected random coordinates in the image (similar to the arbitrary landmark selection process), computed an elastic deformation and calculated the distribution of pair-wise distances between these points, storing the parameters that achieved the highest correlation between the distributions of the landmarks and the ones generated by sampling deformation parameters. We assumed that the Gaussian sigma was the main parameter determining the magnitude of the deformations and hence, fitted a third-degree polynomial relating the value of sigma with the number of grid points that best matched the distribution of pair-wise distances between landmarks obtained for each histological section. From this curve, we defined a range of sigma values for the in silico datasets: 0 (no deformation), 0.5, 1, 1.5, 3 and 5. For each sigma, the number of grid points was determined from the polynomial function fitted before, producing visually realistic deformations of increasing magnitude for each section. Additionally, for each section, we intrinsically evaluated the reproducibility of the registration by producing five replicates for the same sigma value (i.e., despite different deformation grids, all sections should converge to the same ground truth position). Each dataset was composed of 10 sections, with each section being used to generate 25 variations (five sigma values and five replicates).

### Immunohistochemistry datasets

#### Dataset for manual landmarking

Caudal vertebra sections used for manual landmarking were obtained from a previous study^62^. Briefly, samples were decalcified in an EDTA solution in double distilled water (0.14 g/ml, pH 7.4, buffered with ammonium hydroxide) for 2 weeks at 4°C, rinsed for 24 h in PBS, dehydrated through increasing concentrations of ethanol followed by xylene and embedded in paraffin (formalin-fixed paraffin embedded sections, FFPE). Sections with a thickness of 9 μm were cut using a microtome (HM 355S, Microm International GmbH, Walldorf, Germany). The sections were rehydrated and stained with ready to use Haematoxylin and Eosin solutions (MHS1 and HT110116, Sigma Aldrich Inc., St Louise MO, USA). Following dehydration, the sections were mounted with Cytoseal 280 (Richard Allan Scientific, Kalamazoo, MI, USA). Brightfield images of the sections were taken using an inverted Zeiss microscope (Axiovert 200M, Stuttgart, Germany), with a pixel size of 1.25 µm. Six sections with the least number of visual artefacts were selected for analysis.

#### Dataset for LivE characterisation of bone cells

Femur sections used to characterise bone cells in their L*iv*E environment were obtained from a previous study^25^. On day 49, one sample was randomly selected, and the femur was excised, the femoral head was removed, placed in 4% neutrally buffered formalin for 24 h and subsequently decalcified in 12.5% EDTA for 10–14 days. The sample was embedded in paraffin and the complete fracture callus was cut in 10 μm longitudinal serial sections (consecutively along the perpendicular direction to the longitudinal axis). Every 10^th^ section was stained with Safranin-O, and additional sections in between were stained for Sclerostin and RANKL. Images were taken with Slide Scanner Pannoramic 250 (3D Histech, Budapest, Hungary) at 20× magnification, with a pixel size of 0.39 µm. A set of consecutive 12 sections with the least number of visual artefacts were selected for analysis (six stained for Safranin-O, five stained for Sclerostin and one stained for RANKL).

### Spatial transcriptomics datasets

#### Femur defect model

Femur sections used for the ST analysis were obtained from a previous study^21^. Briefly, all mice were euthanised 10 hours after the final cyclic mechanical loading session, five weeks after surgery and samples were processed for ST according to an existing protocol^63^. ST analysis was performed on five-micrometre thick longitudinal sections from explanted femurs (n = 1 section for Control and Loaded groups) using the Visium Spatial Gene Expression for FFPE assay (10x Genomics, Pleasanton, USA), with each section being placed on a 6.5 mm by 6.5 mm capture area of a Visium Spatial Gene Expression slide. Sections were probe hybridised with 20,551 genes targeted (10x Genomics, Visium Mouse Transcriptome Probe Set v1.0) and spatial transcriptomics libraries were prepared. Libraries were sequenced on an Illumina NovaSeq6000 System (Illumina, San Diego, USA) at a sequencing depth of approximately 75 to 120 million reads per sample. Demultiplexing and manual alignment of the sequencing data to the histological image were performed using the SpaceRanger analysis pipeline (10x Genomics, version 2.1.0).

#### Caudal vertebra model

Caudal vertebra sections used for the ST analysis were processed and sequenced in the same experiment described above, using WT mouse samples from a previous study^60^. Briefly, mice were euthanised 24 hours after the final cyclic mechanical loading session. ST analysis was performed on five-micrometre longitudinal sections from explanted vertebrae (n = 1 section for Control and Loaded groups). Both sections were placed on the same capture area, side-by-side.

### Micro-finite element analysis

Micro-CT images were analysed using micro-finite element (micro-FE) analysis to estimate local mechanical signals within bone. Image voxels were converted to 8-node hexahedral elements, with each model containing between 15 to 20 million elements. All micro-FE models assumed that bone tissue behaved within the linear elastic regime and considered a Poisson’s ratio of 0.3. Micro-FE simulations were computed using the parallel micro-FE solver ParOSol^64^ on the Euler cluster operated by the Scientific IT Services at ETH Zurich (on AMD EPYC processors, 2.6–3.3 GHz).

#### Femur defect model

Femur micro-FE models were assigned heterogeneous material properties, with bone density values between 395 mg HA/cm^3^ and 720 mg HA/cm^3^ being linearly converted to Young’s modulus, and setting a value of 2 MPa for marrow voxels (non-bone voxels surrounded by bone voxels), as previously established^23^. Control and loading in vivo scenarios were modelled with two loading conditions: axial compression and a combination of axial compression and bending, respectively. For axial compression, mesh nodes at the proximal end were fully constrained, while nodes at the distal end were displaced by 1% of the model height (longitudinal axis). The bending condition was implemented through affine displacements with a ±0.5-degree rotation around the x-axis at the proximal and distal ends. The simulations computed the Green-Lagrangian strain tensor, from which effective strain^65^ (EFF, in µε, dimensionless) values were determined and linearly scaled to match the in vivo loading condition of 16 N for the loaded group. A fixator stiffness of 16.8 N/mm was considered with a moment arm of 5 mm. A force of 10 N was used for the control group^23^.

#### Caudal vertebra model

Vertebrae micro-FE models based on the binary micro-CT images were assigned homogeneous material properties with a Young’s modulus value of 14.8 GPa for bone and 2 MPa for marrow, as previously established^20^. Two cylindrical discs were added at the proximal and distal ends of the vertebra model, mimicking the role of the intervertebral discs^20^. For the vertebrae models, only the axial compression loading condition was considered. The simulations computed strain energy density (SED, in MPa), from which EFF values were determined after linear rescaling to match the experimental conditions: 8 N for loaded mice and 4 N (physiological loading^66^) for the sham-loaded group (0 N).

#### Mechanoregulation analysis from time-lapsed micro-CT data

The mechanoregulation analysis of micro-CT data was based on the estimation of conditional probability of a bone remodelling event (formation, quiescence, resorption) to occur at each value of mechanical signal (EFF), as previously described^20^. The analysis was done for weekly intervals (i.e., weeks 4–5 considered the micro-CT images from weeks 4 and 5 and the micro-FE from week 4), from week 3 (i.e., time-point at which the defect bridged) and for the entire 4-week observation period for femur defect and caudal vertebra datasets, respectively.

### Validation of the 2D-3D registration algorithm

The validation of Spatial μProbe focused on three aspects: accuracy, precision and sensitivity. First, we evaluated the 2D-3D registration algorithm by comparing it to a gold-standard established with manual landmarking. Next, we assessed its precision with a reproducibility analysis and convergence success rate through a large-scale in silico study using unimodal synthetic data. Finally, we measured the error and sensitivity to detect biological differences by leveraging the results from the in silico study.

#### Evaluation of 2D-3D registration accuracy with manual landmarking

The accuracy of Spatial μProbe to register 2D histological sections into 3D micro-CT images of the same sample was evaluated by comparing the registration output with manual landmarking, which is regarded as a gold-standard approach to validate image registration algorithms^67^. First, the SlicerIGT^55^ extension available for Slicer3D^68^ was used to manually landmark and register six histological sections (see Datasets) to their corresponding high-resolution micro-CT scans (voxel-size: 2.1 µm). For each section, one operator (F.C.M) identified over 100 pairs of landmarks (minimum: 102, maximum: 166, average: 132) between the 2D image and the 3D micro-CT scan, from which a ground-truth similarity transformation was determined by minimising the root mean square error between pairs of landmarks. Landmarks were spread across the sections to improve global registration accuracy, following available recommendations^5^. Next, the same histological sections were registered to their corresponding in vivo micro-CT scan using Spatial μProbe. The evaluation relied on the Target Registration Error^5,69^ (TRE) and the angular deviation, defined as the angle between the normal vectors defined by planes of the registered section using the ground-truth transformation and the result obtained with Spatial μProbe.

Additionally, we considered the pairs of landmarks identified with SlicerIGT to estimate the magnitude of deformations associated with sample embedding and microtome sectioning. We computed the pairwise distance between the positions of all landmarks on the histological section and subtracted the pairwise distance between the positions of all landmarks on the micro-CT image (which is expected to be undeformed, since the scan is taken prior to tissue embedding and sectioning). The resulting distribution of differences between pairwise distances described the magnitude of local deformations of the histological section.

#### Evaluation of 2D-3D registration precision

To evaluate precision, we repeated the registration of the sections from two datasets five times (mouse vertebra model: the same six sections used for manual landmarking; mouse femur: 14 sections from an osteotomy experiment; see Datasets). For each section, we computed the median of the optimised registration parameters and determined the median absolute deviation (MAD) as a non-parametric dispersion metric. We reported the mean ± standard deviation MAD across each dataset. For translational parameters, the deviation of each section from the median was quantified as the Euclidean norm of the 3D displacement vector, scaled by the voxel size of the micro-CT images (10.5 µm). For rotational parameters, to account for the non-linear geometry of rotations, Euler angles were converted to rotation matrices and parameterised as rotation vectors (axis-angle representation). The Fréchet mean (central rotation) was computed on SO(3), with deviations quantified as geodesic distances (in degrees) from this mean. For 2D uniform stretching parameters, we first log-transformed the multiplicative scaling parameters to convert them into additive deviations. We computed the absolute deviations from the median in log space. MAD values in log space were converted to multiplicative scaling through an exponential operation, yielding a percentage relative to the median scaling factor.

#### Evaluation of 2D-3D registration convergence and sensitivity in silico

Next, we considered unimodal in silico experiments to characterise the convergence of the 2D-3D registration algorithm. Briefly, sections with known ground-truth positions were produced from 3D micro-CT images and registered back onto the source 3D image. The registration parameters were compared with the ground-truth values, highlighting the convergence rate of the registration for varying sections and initialisation conditions.

First, we identified a suitable set of image similarity functions: Dice similarity coefficient (Di), surface Dice similarity coefficient (DS), normalised Mutual Information (nMI), Pearson correlation coefficient (Pe), Jaccard Index, normalised Root Mean Squared Error (nRMSE), the nMI, nRMSE, Pe and the structural similarity index (SSIM) between the Euclidean distance transformed images (EDTnMI, EDTnRMSE, EDTSSIM and EDTPe, respectively). To reduce the number of linear combinations between similarity functions, we performed a preliminary analysis to evaluate cross-correlation among them in our application, from which four similarity functions were selected for subsequent experiments (see Supplementary Note 2). For each micro-CT image, we produced ten sections along the x-axis (longitudinal sections) and z-axis (cross-sectional sections) and computed the value of these similarity functions between each section and all other sections. Similarity values were aggregated based on relative displacement between pairs sections. Functions were selected based on smoothness, uniqueness of global minimum and least cross-correlation with other functions, converging on the set: EDTPe, EDTnMI, nMI and DS.

#### 2D-3D registration of in silico datasets

The next analysis was designed to investigate the effect of the following five factors on the registration outcome: the amount of bone material in the image (as a proxy for the amount of image features available in 2D sections), sectioning orientation (longitudinal or cross-sectional), presence of local deformations and their magnitude, image similarity functions guiding the optimisation step, and the initialisation of the registration transformation parameters. Registration trials were initialised with parameter sets obtained at five equidistant locations (0%, 25%, 50%, 75%, 100%) along the linear segment connecting the origin (centre of the image) and the ground-truth transformation. Rotations were additionally initialised at 62.5% and 87.5% values to increase the density of values in the results for this transformation.

Next, we registered the sections generated in silico with rigid and elastic transformations. Trials were carried out for each sample (four micro-CT images), each sectioning orientation (longitudinal and cross-sectional), each replicate dataset (five datasets of 25 images), each linear combination of similarity functions (15 cases) and each initialisation location (grid of 35 points). Sections were registered to the source 3D images in an unsupervised way to evaluate the ability of the registration algorithm to recover their original location. In all registration trials, we fixed the search range of the Powell method used to optimise image similarity to 25 for rotation and translation values, and 0.025 for in-plane stretching parameters when registering images from elastic datasets. A trial was considered successful if the 99th percentile of the Dense Registration Error^31^ (DRE; similar to the Target Registration Error but evaluated across all points of the image) between the ground-truth transformation and the optimised transformation was below a numerical threshold of 10.5. To align results with the imaging resolution, trials from datasets generated with rigid transformations were filtered to ensure median DRE values below one voxel (10.5 µm), while results from datasets produced with elastic transformation were filtered below three voxels (31.5 µm). Results were aggregated per sample and sectioning orientation based on the Euclidean norm of absolute differences in translation and the geodesic distance (smallest angle in 3D space) between the ground-truth and optimised rotation parameters after conversion of Euler angles (XYZ order) to rotation matrices. Rotation and translation values were binned in intervals of 15° and 150 or 250 μm for the femur and vertebra samples, respectively. For each bin, the configuration of similarity functions with the highest convergence rate was identified, with precedence being given to simpler combinations in case of a tie. The in silico simulations were performed on the Euler cluster operated by the Scientific IT Services at ETH Zurich (on AMD EPYC processors, 2.6–3.3 GHz).

Finally, to assess the effects of deviations in registration parameters from the ground-truth on the ability and sensitivity to retrieve relevant biological information after 2D-3D registration, we considered all successful iterations from the in silico validation study, stratified by mouse model and sectioning orientation. We considered the output of micro-FE analysis as an example dataset to be used in multimodal integration and measured the mean absolute error (MAE) between the mechanical signal, as effective strain values of the sections obtained with ground-truth and optimised transformations. These values were visualised in a heatmap, with the average MAE displayed for each pair of translational and rotational offsets to the ground-truth values, filtered at the 99.99% percentile.

### Multimodal data analysis for bone mechanobiology research Preprocessing multimodal data

For each 2D section analysed, we considered the region of interest defined for 2D-3D registration and extracted the following datasets from time-lapsed 3D micro-CT images: mechanical signal values from micro-FE analysis, bone formation over all weeks of the experiments and bone remodelling data in the last week before the end of the in vivo experiments.

#### Mechanical signal from micro-FE analysis

Mechanical signal values were collected in a voxel-wise fashion for each pixel on the 2D images mapped onto the 3D image.

#### Longitudinal bone formation from time-lapsed micro-CT data

Binary micro-CT images from each week were superimposed, and all bone voxels were encoded based on the weeks in which they were present. Voxels were grouped based on the earliest time-point in which they were formed and remained present until the last time-point (i.e. the longest continuous interval with respect to the last time-point). Additionally, to estimate the day at which each voxel was formed, we extended the concept of SpatialTime^70^ by computing the Euclidean distance transform of the binary image at each time-point and assigning voxels formed each week with a value based on their distance to the surface. For a micro-CT image at a given time-point, surface voxels were assigned the day the micro-CT scan was taken, and the remaining voxels between the surface voxels of the current and previous time-point were assigned a value in days between the two weeks based on their Euclidean distance to the surface. The bone formation week and day were collected in a voxel-wise fashion for each pixel on the 2D images mapped onto the 3D image.

#### Late-stage bone remodelling from time-lapsed micro-CT data

The binary micro-CT images from the last two time-points of each dataset were superimposed, revealing sites of bone formation, resorption and quiescence. Since bone resorption values are not present in 2D sections of the last time-point, we classified quiescent voxels adjacent to resorption volumes as resorption voxels. Specifically, we computed the average thickness of formed and resorbed volumes based on the values of the Euclidean distance transform of the surface values of the image containing all remodelling volumes. Next, for the voxel assigned to each pixel in the 2D section, we checked the events assigned to all voxels in a sphere with the radius matching the average remodelling thickness determined before and computed their weighted average (with resorption, quiescence and formation being assigned 3, 2 and 1), producing a visually realistic classification of remodelling events for all pixels in the 2D section.

### L*iv*E characterisation of bone cells

We used Spatial μProBe to register a dataset of consecutive longitudinal histological sections of a mouse femur onto the corresponding micro-CT image. Transformations were initialised using parameters obtained from ParaView after visually identifying an approximate section from the 3D image. IterAR was used to optimise the image similarity based on the combination of four similarity functions: Di, DS, nMI and EDTnMI. Translation and rotation transformations were initially applied until a visually realistic alignment was achieved, followed by iterations with in-plane stretching to correct for small deformations. An additional deformable registration step was performed between the resulting images, assuming the shape of the 2D section obtained from the micro-CT image as the reference shape. The coordinates of the 2D pixels were mapped to 3D and the values of the mechanical signal (EFF), bone formation and bone remodelling data were collected. A kernel density estimation (KDE) plot was used to visualise the distribution of EFF values for each section. Additionally, cells and pixels in the 2D section were categorised according to their L*iv*E environment, considering the magnitude of mechanical signal and type of remodelling events in their vicinity.

### Analysis of Spatial Transcriptomics data

#### Estimation of cell-type proportions using Starfysh

We applied Starfysh^34^ on the femur defect and caudal vertebra ST datasets. First, we identified 15 major cell types in musculoskeletal and neighbouring tissues: osteoblasts, osteoclasts, osteocytes, mesenchymal stem cells, hematopoietic stem cells, endothelial cells, adipocytes, chondrocytes, myocytes, fibroblasts, macrophages, T-cells, B-cells, neutrophils and pericytes. For each cell type, we identified 10-20 marker genes (see Supplementary Data 1). We provided this input to Starfysh and set the value of “number of genes to keep” (n_genes) to 12,000. The pipeline included the default Archetypal analysis to expand the input list of marker genes, and each model was trained for 500 epochs.

#### Differential gene expression analysis

We used DESeq2^71^ to perform differential gene expression analysis between Control and Loaded ST datasets. Specifically, we compared groups of spots based on bone formation time-points (categories: original bone, bone formed in weeks 0-2, bone formed in weeks 2-3, bone formed in weeks 3-5) between both datasets. We considered the intersection of the top 20% expressed genes on both datasets and used the package scran to compute the scaling normalization of gene expression counts through deconvolving size factors. In addition, null hypothesis testing was conducted using the likelihood ratio test, with a parametric fit. Shrunken log fold change factors were determined using the “apeglm” fit. Differentially expressed genes (DEGs) were selected using an FDR-adjusted p-value cutoff <0.05 and an absolute log2-fold change >0.5.

For comparisons between Control and Loaded datasets for the same time-points, we also determined the log2-fold change across all genes and clustered the results using agglomerative clustering, supported by the graph of k-neighbours of the data. The number of neighbours and the Power parameter for the Minkowski metric for the graph of k-neighbours, and the number of clusters for the agglomerative clustering algorithm were optimised to maximise the silhouette score of the clustering result, producing clusters with at least 10 elements. This optimisation suggested six clusters, with the k-neighbours graph being created based on 5 neighbours with a Power parameter of 1.25. Time-lapsed curves of log_2_ fold-changes for each cluster were summarised with quadratic functions.

#### Identification of spatially variable genes

We applied SPARK-X^56^ to identify spatially variable genes in all datasets. To highlight genes that were spatially variable within the region available for multimodal integration, we pre-filtered ST spots and only analysed spots in bone tissue and inside the 3D micro-CT image, after 2D-3D registration.

#### Super-resolution of ST data

We used iStar^30^ to produce super-resolved ST data at the same resolution as the micro-CT datasets, using the spatially variable genes identified in the previous step. From this set, we only considered genes with a mean count above 1 (counts of unique molecular identifier, UMI, above 1) across all the spots in bone and successfully mapped to the micro-CT image, to minimise the effect of noise from lowly expressed genes on the model training. Next, given that the magnitude and relative spatial variation in gene expression is vital for subsequent steps, for each model run, we overfitted the ST dataset by sweeping through the entire dataset at every epoch and training for 2500 epochs. As input, we provided gene expression counts corrected with SCTransform^57^. Since the model predictions are bounded to the range [0, 1], super-resolved gene expression maps were linearly scaled based on the original range of values. ST histological sections were passed as input in their native resolution and the super-resolved maps were rescaled to the micro-CT resolution. Gene expression values were collected for each gene for all relevant bone pixels of the ST histological section. A super-resolution transformation was also performed on the output of other analysis with ST datasets (i.e., cell-type proportion estimation with Starfysh). We evaluated the performance of iStar on the bone fracture healing dataset (Supplementary Note 3).

#### Mechanical and temporal regulation of gene expression

To investigate mechanoregulation of gene expression, we considered the genes super-resolved in the previous step. Gene expression was modelled as a function of mechanical signal (EFF), time (in days), and both using generalised additive models (GAMs). The number of splines was set to 25 and an optimal penalty factor (lambda) was optimised for each gene to produce a model with 5 and 25 effective degrees of freedom, for 1D and 2D models, respectively. The mechanical signal values for GAMs associated with gene expression for each time-point were additionally filtered at their 99^th^ percentile to reduce the influence of data sparsity beyond this point in the model. Next, we applied non-negative matrix factorisation (NMF) to the GAMs from all genes, using six components (the smallest value that retained over 99% of the explained variance in the data). Components were filtered to only retain the major sources of variation, without loss of explainability (less than 0.1% change in explained variance). GAM curves were normalised prior to NMF analysis, such that all genes had a total expression of 1, to neutralise differences in expression magnitude while conserving differences in the shape of the GAM curve. NMF factors were standardised across all genes and clustered with agglomerative clustering, with parameter settings maximising the silhouette score, as above (see “Differential gene expression analysis”), under the constraints that clusters should have a minimum and maximum of 3 and 50 elements, respectively, to ensure sufficient cluster size for all clusters while maintaining a fair level of granularity of the dataset. An additional step was performed to account for elements assigned to a sub-optimal cluster: sample-specific silhouette scores were computed and those with a negative score were assigned to the closest cluster, based on the shortest Euclidean distance to the median of the neighbouring clusters. This operation was performed iteratively until no further changes in cluster assignments were possible and was verified with improvements in the average silhouette at each stage. The mean curve and its 95% confidence interval of each cluster was estimated using bootstrapping, with 2500 resampling iterations.

#### Functional enrichment analysis

The biological and functional characterisation of the clusters determined for each dataset was performed using g:Profiler^58^, using a Benjamini-Hochberg FDR threshold of 0.05, with a focus on the results from Gene Ontology, KEGG Pathways and WikiPathways. Additionally, this approach was also applied to groups of genes (min. 3 elements) in clusters of one sample that were also aggregated in a cluster on the other sample (i.e. Control and Loaded conditions for ST datasets of the mouse femur and 6^th^ caudal vertebra), to investigate shifts in gene expression resulting from cyclic mechanical loading.

#### Analytical characterisation of gene mechanoregulation

To provide an interpretable analytical description for the mechanical regulation of gene expression, we provide a framework for fitting mathematical basis functions of different complexity to gene expression data as a function of effective strain. Specifically, the following functions are available (with y representing gene expression and x the mechanical signal):

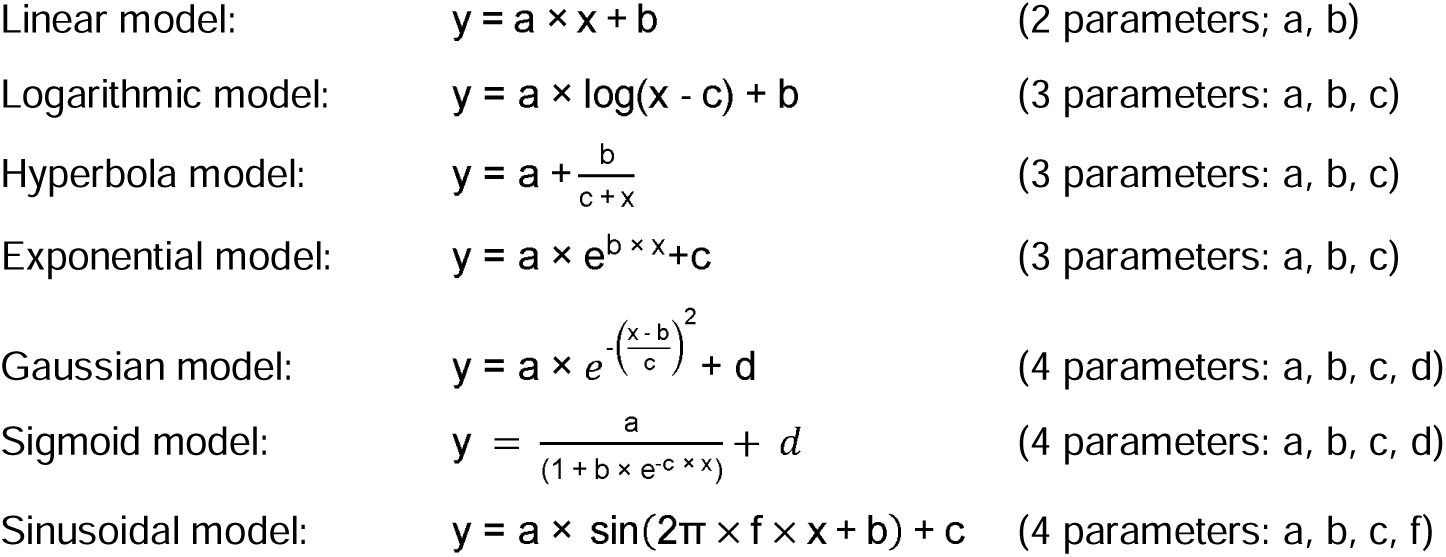

Modulated sinusoidal model:

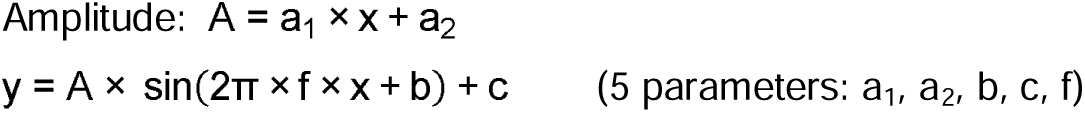

Additionally, we also considered well-established mathematical descriptions in biology:

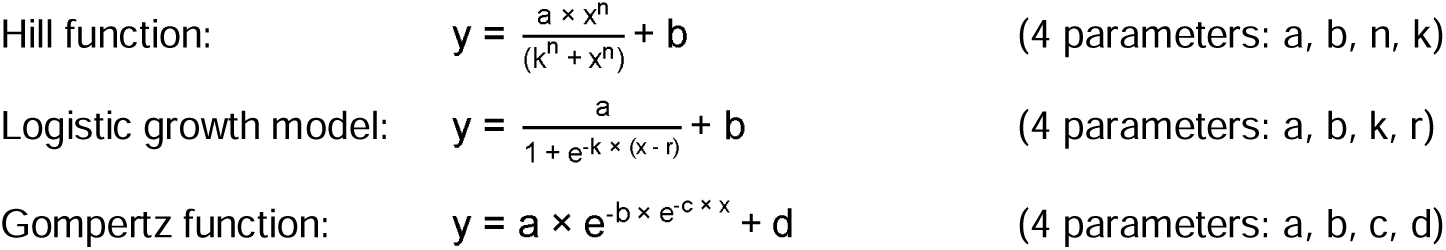

For this application, all functions were fitted per time-point to the original datapoints containing the value of the mechanical signal and gene expression. To maximise the chances of achieving a successful curve fit, each function was fitted 500 times using the method of Sequential Least Squares Programming (SLSQP), with the optimisation being initialised with parameters generated from a normal distribution. The mean and scale of the normal distribution were defined in a grid of 25 values, with the mean ranging between 0 and 0.8 (in steps of 0.2) and the scale ranging geometrically between 0 and 0.8. The curve fit optimisation considered the Huber loss, with a delta value of 1. Since several functions can be fitted to the same set of points, we propose a voting mechanism to sort the curve fitting results, that aims to provide the simplest set of functions that best fits the data for each time-point. First, the Bayesian information criterion (BIC) was determined for all iterations. Next, successful iterations were filtered per function and only those within 1% of the minimum Huber loss achieved for each function were kept. For each time-point, functions were filtered based on the median Huber loss, with those within 1% of the lowest median value being considered for subsequent steps. Finally, we considered all the combinations of the set of functions available per time-point and identified the set of functions with the lowest cumulative BIC value across all time-points.

### Statistical analysis

Mean and median values reported are accompanied by their standard deviation and interquartile range, respectively. For all statistical analysis, normality was first assessed with the Shapiro-Wilk test, followed by equality of variances using Levene’s test. Statistical tests for subsequent comparisons were chosen accordingly. Significance was set at p < 0.05 in all experiments, unless indicated otherwise. In Figure 3, distributions of mechanical signal values per registered section (Fig. 3b) and for bone formed at different time-points (Fig. 3d) were compared with Kruskal-Wallis test, followed by post hoc pairwise multiple comparisons Conover’s test, using the Holm–Bonferroni method for adjusting p-values.

### Visualisations

Visualisations of 3D images were produced using ParaView 5.12^52^. Images were first converted to VTI file format, and the “Threshold” tool was applied to identify the object in each 3D image. A smooth filter was applied to the surface of each object. Renderings were produced using the ray-tracing tool (“Ambient Samples”: 1, “Samples Per Pixel”: 10, “Progressive Passes”: 10, “Light Scale”: 0.8).

## Ethics declaration

All mouse experiments were performed in accordance with relevant national regulations (Swiss Animal Welfare Act, TSchG, and Swiss Animal Welfare Ordinance, TSchV) and authorised by the Zürich Cantonal Veterinary Office (approved license numbers: ZH176/2010, ZH09/2018, ZH229/2019, ZH36/2022; Kantonales Veterinäramt Zürich, Zurich, Switzerland).

## Author contributions

Conceptualization was done by F.C. Marques and R. Müller. Methodology development was carried out by F.C. Marques and R. Müller. Software implementation was performed by F.C. Marques, N. Ohs, C. Goenczoel, and J.J. Kendall. Validation was conducted by F.C. Marques and A. Singh. Formal analysis was done by F.C. Marques. Investigation was carried out by N. Mathavan, D. Yilmaz, D. Günther, G.A. Kuhn, and E. Wehrle. Data curation and visualization were handled by F.C. Marques and R. Müller. Resources were provided by R. Müller. The project was supervised by F.A. Schulte, E. Wehrle and R. Müller. Project administration was managed by R. Müller. Funding acquisition was led by R. Müller with contributions from N. Mathavan and E. Wehrle. The original draft was written by F.C. Marques and R. Müller, with review and editing by F.C. Marques, N. Mathavan, N. Ohs, C. Goenczoel, J.J. Kendall, G.A. Kuhn, F.A. Schulte, E. Wehrle and R. Müller.

## Acknowledgements

We acknowledge the ETH Phenomics Centre (EPIC) of ETH Zürich, and particularly S. Freedrich, for assistance with the in vivo experiments. Spatial transcriptomics was performed at the Functional Genomics Centre Zürich (FGCZ) of the University of Zürich and ETH Zürich. Tissue processing and paraffin embedding were performed at ScopeM at ETH Zürich. Sectioning, staining, and imaging were performed at ScopeM at ETH Zürich and at the Institute of Pathology and Molecular Pathology at the Universitätsspital Zürich (USZ). We thank Christopher M. O’Neill and Bryant Schroeder for the early support with software for affine transformations, and Cael McLennan for the support in the manual annotation of histological sections. We thank Dr. Andreas Trüssel for collecting part of the data used for validation of the 2D-3D registration algorithm using landmarking. High-performance computing data processing and analyses were performed on the ETH Euler cluster.

## Funding

This work was supported by European Research Council - Horizon 2020 grant ERC-2016-ADG-741883, European Union Marie Skłodowska-Curie Actions - Horizon 2020 grant 101029062, MechanoHealing-MSCA-IF-2020, and the Swiss National Science Foundation (COST Action GEMSTONE IZCOZ0_198152/1, SLIHI4BONE 213520).

## Notes

https://doi.org/10.5281/zenodo.15747876

### Competing Interest Statement

The authors have declared no competing interest.

### Summary of Updates

- Updates to figure 1, and formatting issues to figures 4 and 6. - Editing of text in Introduction, Discussion and certain Results sections - Improved flow between sections - Add missing citations and fix citation issues - Add author to author list.

