## SupplementaryMaterial for "Spatial μProBe: a correlative multimodal imaging approach for spatial profiling of biological micro-environments"

Supplementary Figures

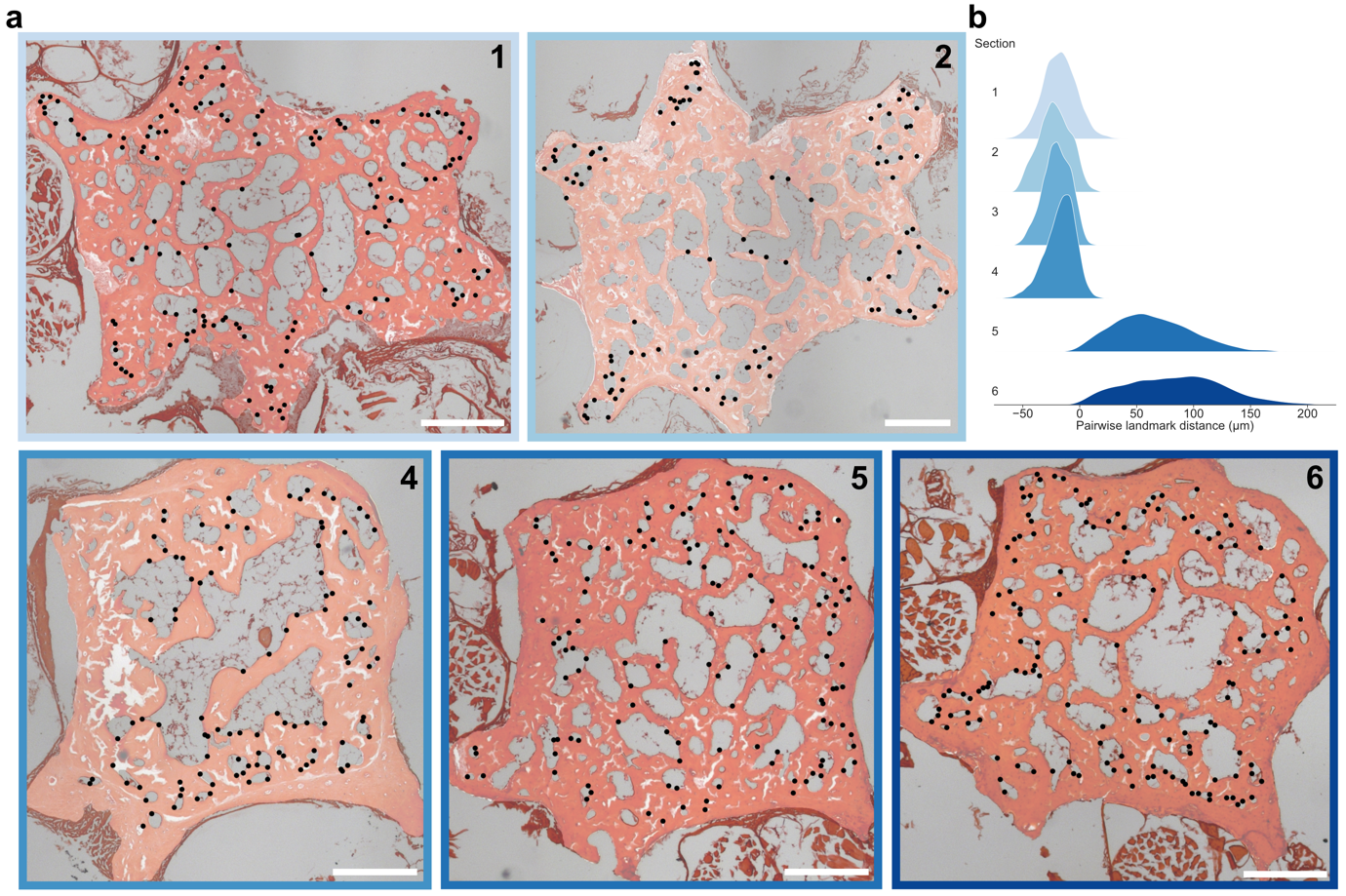

Supplementary Figure 1 | Landmarking of histological sections with SlicerIGT.

a, Sections and landmarks identified in black for the ground-truth registration with high resolution ex vivo micro-computed tomography (micro-CT). Section 3 is not present, since it is displayed in Fig. 2a. Scale bar: 200 µm. Section 1: 140 landmarks; section 2: 102 landmarks; section 4: 104 landmarks; section 5: 135 landmarks; section 6: 146 landmarks.

b, Kernel density estimation (KDE) plot of pairwise distances between the landmarks identified.

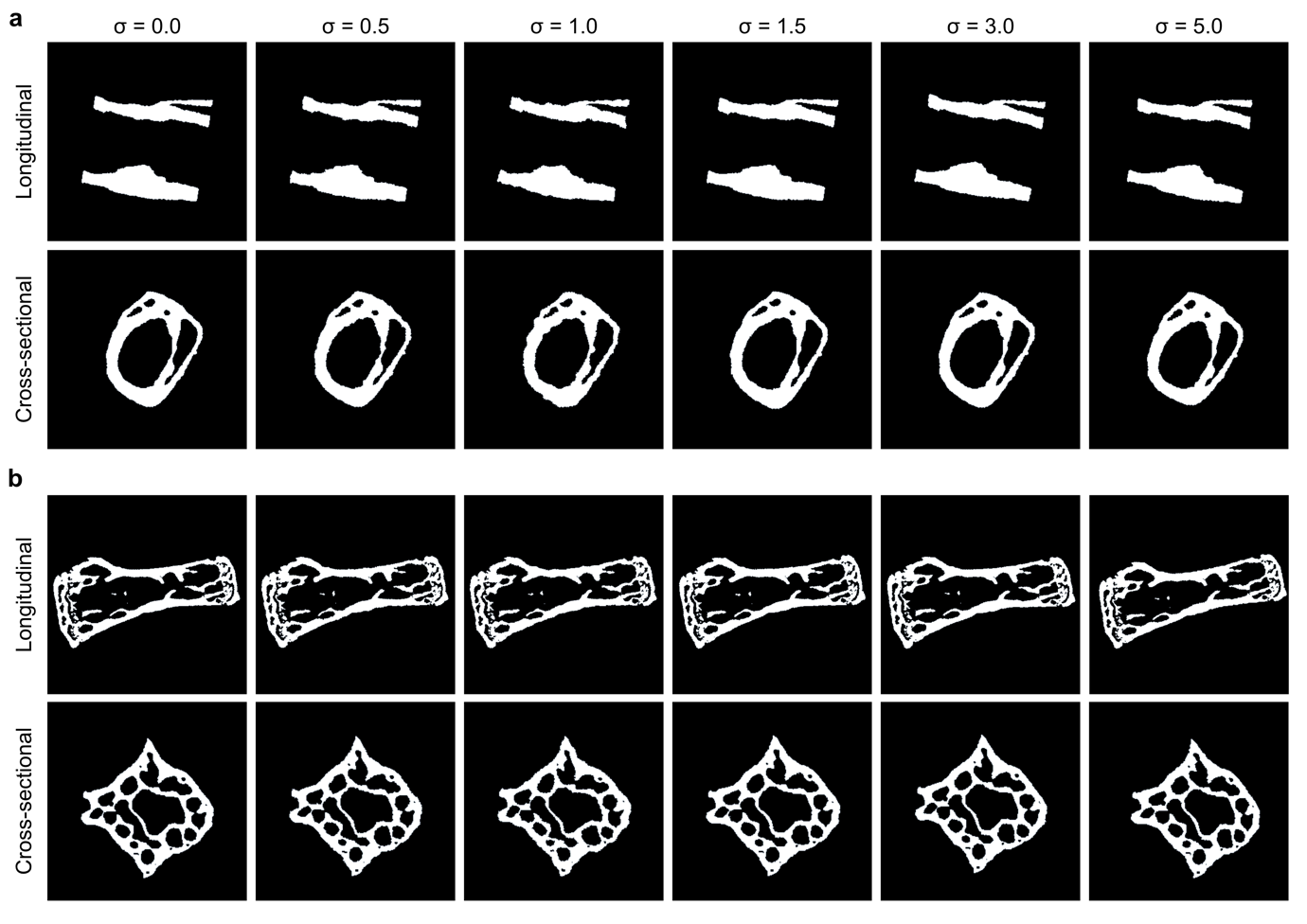

Supplementary Figure 2 | Representative examples of synthetic datasets of 2D sections.

a, Sections generated in silico from 3D micro-CT images across longitudinal and cross-sectional orientations (including off-axis rotations), for a mouse femur sample. Columns illustrate the influence of the deformation σ parameter, used to recreate non-linear deformations in the 2D sections generated from rigid transformations.

b, Sections generated across longitudinal and cross-sectional orientations (including off-axis rotations), for a mouse 6^th^ caudal vertebra sample. Columns represent the same quantity as in a.

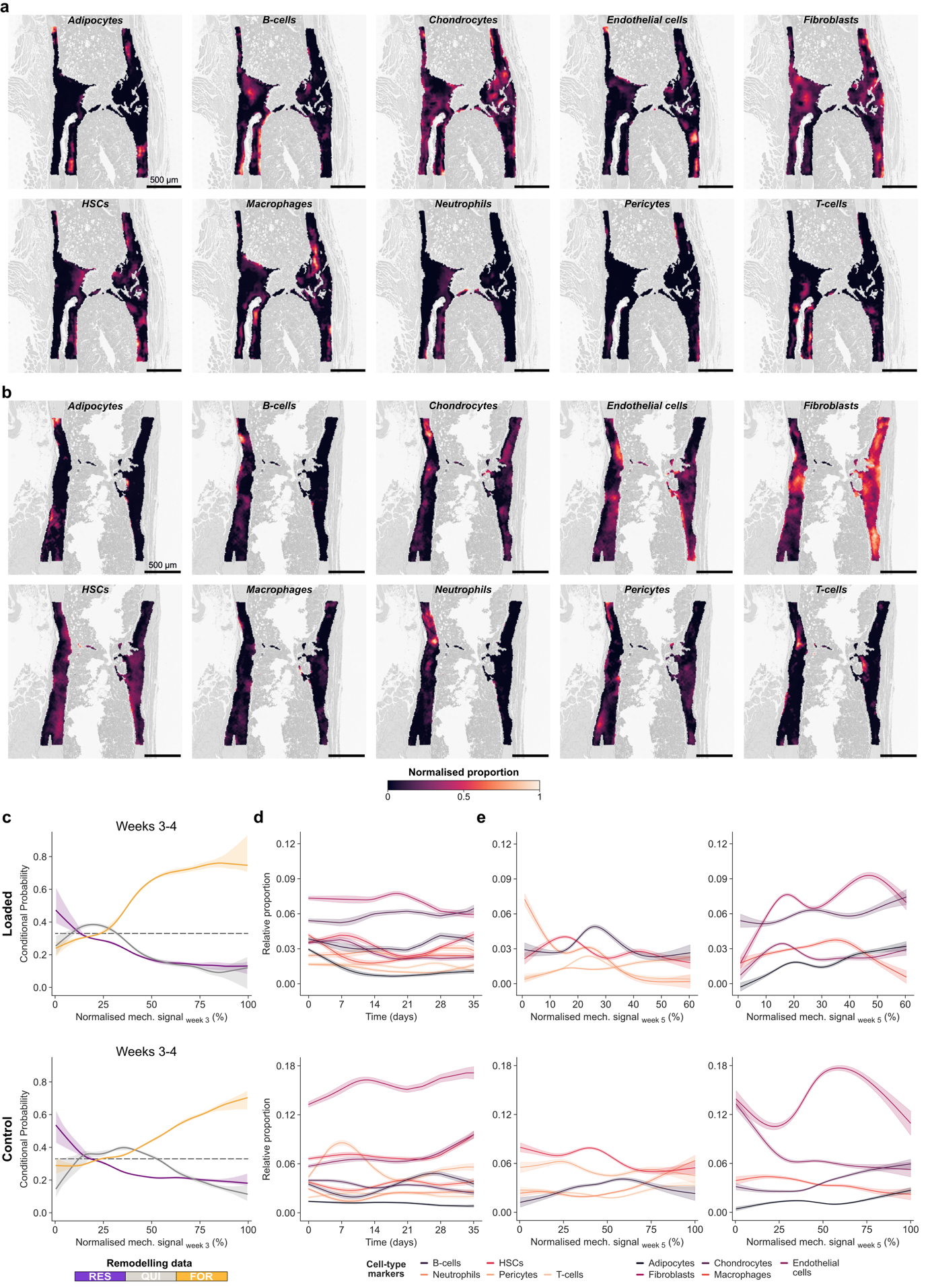

Supplementary Figure 3 | Bone mechanoregulation at the tissue and cell levels, using Starfysh cell-type relative proportions.

a, Cell-type proportions determined with Starfysh^1^ and super-resolved with iStar^2^, for the Loaded sample. Relative proportion values were normalised to the range 0-1 to support the visual comparison of individual cell-type distributions. Variability in estimated cell-type proportions is affected by sectioning artefacts and the sparsity of transcriptomic markers in certain locations. Still, our analysis focuses on high-confidence populations relevant to bone adaptation and regeneration.

b, Cell-type proportions determined with Starfysh^1^ and super-resolved with iStar^2^, for the Control sample. Values are normalised as in a.

c, Conditional probability curves connecting the surface mechanical environment as effective strain, normalised to the 99^th^ percentile, with surface (re)modelling events (FOR: formation, QUI: quiescence, RES: resorption). The plots show the mean probability line and its corresponding 95% confidence interval, per (re)modelling event for the weeks 3-4 after bootstrapping (2500 iterations) of a LOWESS operation. The dashed line at 0.33 identifies the probability of a random event for a ternary classification case.

d, Association between the relative proportion the presented in sub-panel a) with the time-points of bone formation identified with time-lapsed micro-computed tomography (micro-CT), indicating a differential distribution of cell populations across weeks. Curves show the prediction obtained with generalised linear models (GAMs) with 25 splines and 5 effective degrees of freedom, and their 95% confidence interval.

e, Association between the relative proportion of cell-types presented in sub-panel a) with the magnitude of local mechanical signals, revealing cell-type mechanoregulation trends. Curves show the prediction obtained with GAMs with 25 splines and 5 effective degrees of freedom, and their 95% confidence interval. Curves were split into two plots to aid visualisation.

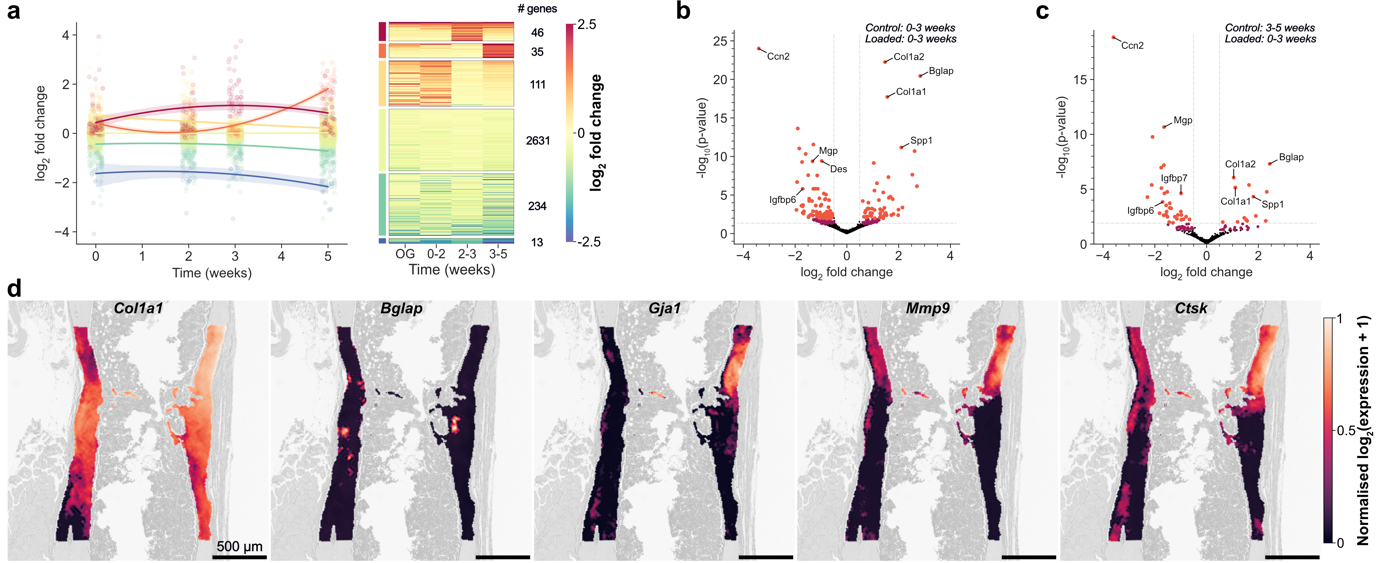

Supplementary Figure 4 | Differential gene expression analysis during bone fracture healing.

a, Agglomerative clustering of genes based on the time-lapsed log_2_-fold change in expression between the Loaded and Control samples. Number of clusters was selected by maximising the silhouette score. Values per cluster are fitted with a quadratic model. The heatmap shows the variations in log_2_-fold change for each cluster (only a maximum of 150 genes are shown).

b, Volcano plot to visualize differentially expressed genes (DEGs) for the comparison between weeks 0-3 of the Control sample with weeks 0-3 of the Loaded sample. Significance criteria: absolute log_2_-fold change > 0.5 (vertical dashed lines); False discovery rate (FDR)-adjusted p-value < 0.0125 indicated in orange (after adjusting for multiple comparisons, horizontal dashed line) and FDR-adjusted p-value < 0.05 indicated in dark pink.

c, Volcano plot to visualize DEGs for the comparison between weeks 3-5 of the Control sample with weeks 0-3 of the Loaded sample. Significance criteria and labelling follow the rules indicated in b.

d, Super-resolved spatial gene expression maps of selected biologically-relevant markers at the fracture site of the Control sample. Osteoblast markers: *Col1a1* and *Bglap*; Gap junction marker: *Gja1*; Osteoclast markers: *Mmp9 and Ctsk*.

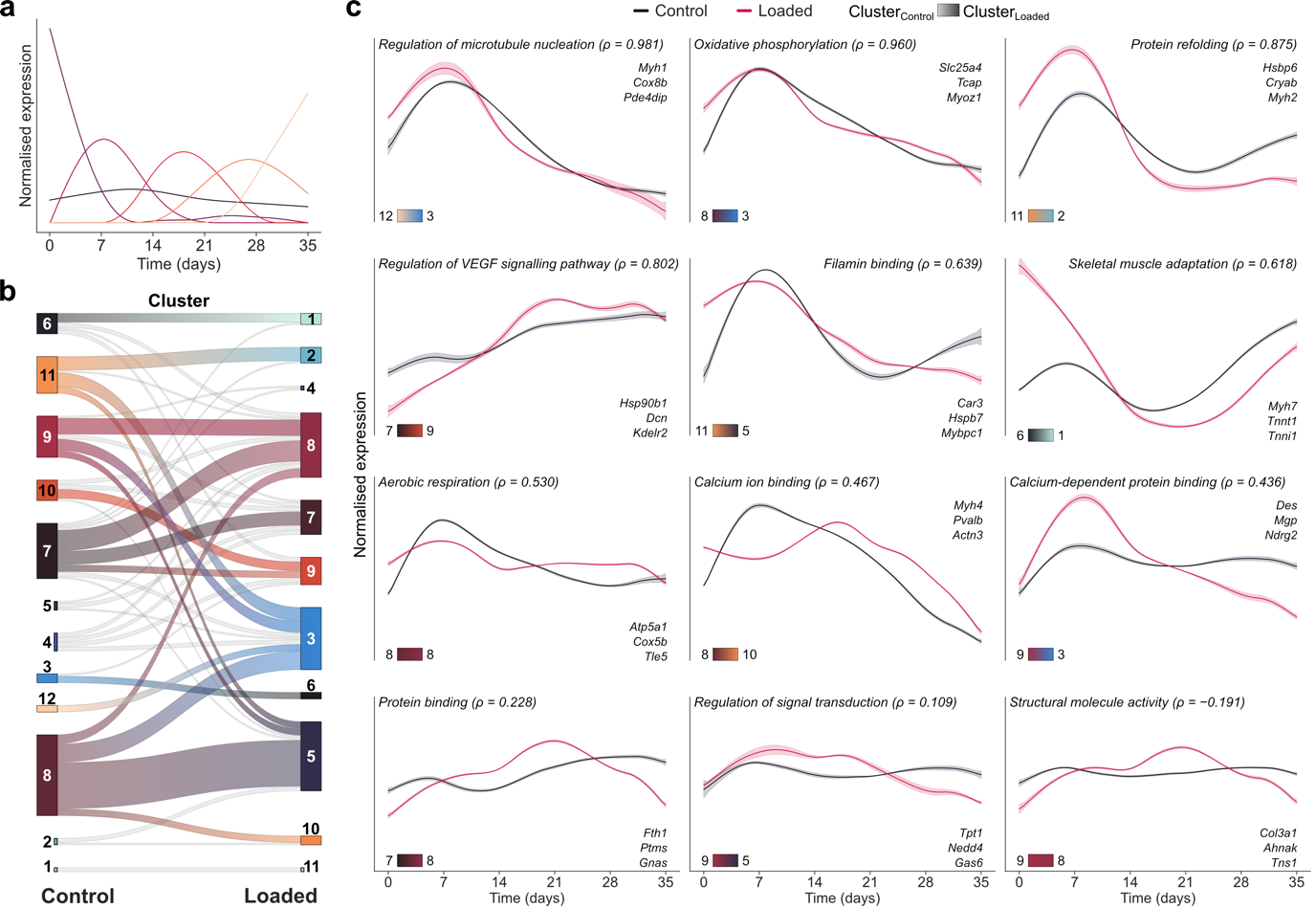

Supplementary Figure 5 | Time-lapsed gene mechanoregulation analysis during bone fracture healing in a mechanically controlled environment.

a, Components obtained after non-negative matrix factorisation (NMF) analysis of the generalised additive model (GAM) curves associating gene expression and time for spatially variable genes (SVGs) of the Control sample. GAM curves were normalised prior to NMF decomposition, to neutralise differences in expression magnitude between genes.

b, Sankey diagram associating clusters computed from NMF factors of the Control and Loaded samples (shown in Fig. 6b and Extended Fig. 3a). Coloured connections represent groups of genes (min. 3 elements) that cluster together in both samples, indicating collective changes in expression.

c, Comparison between the normalised expression of sets of genes between Control and Loaded samples, for the connections indicated in b. The top three most expressed genes are listed, as well as the cluster numbers (referring to Fig. 6c for the Loaded sample and Extended Fig. 3a for the Control sample). A functionally-relevant significant enrichment term identified with g:Profiler^3^ is indicated, determined using only the genes from each gene set. The Spearman correlation coefficient between the curves obtained for both samples is indicated as a proxy for the change in expression trends resulting from cyclic mechanical loading.

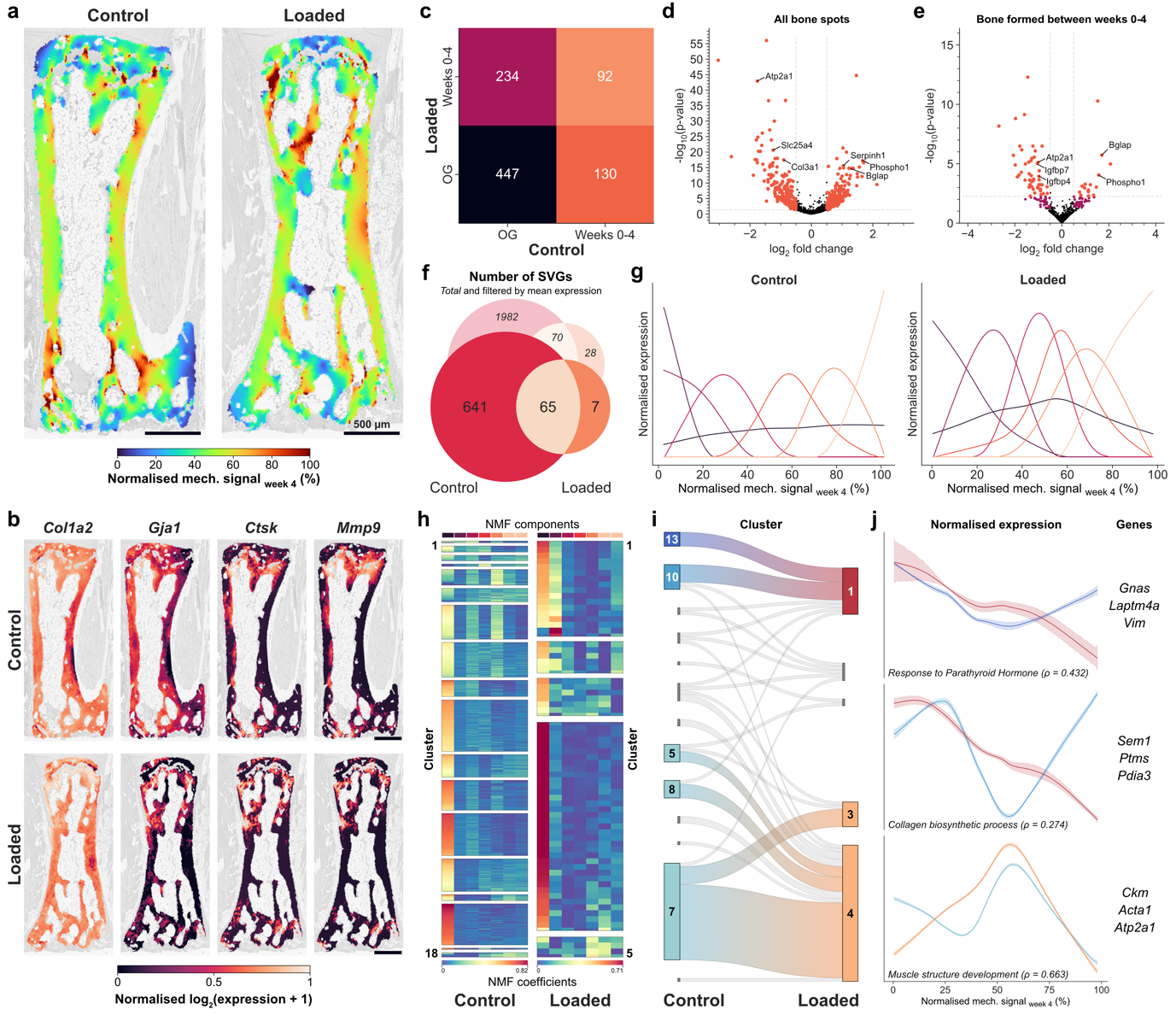

Supplementary Figure 6 | Time-lapsed gene mechanoregulation analysis during bone adaptation in a mechanically controlled environment.

a, Mechanical environment computed with micro-finite element (micro-FE) analysis on the micro-computed tomography (micro-CT) image and mapped to the 2D-3D registered spatial transcriptomics (ST) section of the Control and Loaded samples of a 6^th^ caudal vertebra from mice. Values are normalised to the 99^th^ percentile of the 3D mechanical environment. Scale bar: 500 µm.

b, Super-resolved spatial gene expression maps of selected biologically-relevant markers involved in bone adaptation. Osteoblast markers: *Col1a2*; Gap junction marker: *Gja1*; Osteoclast markers: *Ctsk* and *Mmp9*.

c, Number of differentially expressed genes (DEGs) for pairwise comparisons between bone present at the start of the experiment (OG) and bone formed during the experiment (weeks 0-4). Mechanical loading was applied to the Loaded sample between weeks 0-4. Scale bar: 500 µm.

d, Volcano plot to visualize differentially expressed genes (DEG) for the comparison between all bone spots of the Control and Loaded samples. Significance criteria: absolute log_2_ fold change > 0.5 (vertical dashed lines); FDR-adjusted p-value < 0.0125 indicated in orange (after adjusting for multiple comparisons, horizontal dashed line) and FDR-adjusted p-value < 0.05 indicated in dark pink.

e, Volcano plot to visualize DEGs for the comparison between spots of bone formed during the experiment (weeks 0-4) for the Control and Loaded samples. Significance criteria and labelling follow the rules indicated in d.

f, Number of spatially variable genes (SVGs) detected using SPARK-X^4^ for Control and Loaded samples. Only spots in bone tissue overlapping with the micro-CT registered image were considered. Values indicate the total number of genes detected (*italic*) and the number of genes with a mean unique molecular identifier (UMI) per spot above 1 (UMI counts corrected with SCTransform^5^). Only filtered genes were considered in subsequent steps.

g, Components obtained after non-negative matrix factorisation (NMF) analysis of the generalised additive model (GAM) curves associating gene expression and normalised mechanical signals for SVGs of the Loaded and Control samples. GAM curves were normalised prior to NMF decomposition, to neutralise differences in expression magnitude between genes.

h, Agglomerative clustering of all genes based on NMF coefficients. Clustering settings and results were optimised to maximise the silhouette score, yielding 18 clusters for the Control sample and 5 for the Loaded sample.

i, Sankey diagram associating clusters computed from NMF factors of the Control and Loaded samples (shown in h). Coloured connections represent groups of genes (min. 3 elements) that cluster together in both samples, indicating collective changes in expression.

j, Comparison between the normalised expression of sets of genes between Control and Loaded samples, for the connections indicated in sub-panel i). Lines are colour-coded according to the connections each one describes. The top three most expressed genes are listed. A functionally-relevant significant enrichment term identified with g:Profiler^3^ is indicated, determined using only the genes from each gene set. The Spearman correlation coefficient between the curves obtained for both samples is indicated as a proxy for the change in expression trends resulting from cyclic mechanical loading.

Supplementary Note 1

Semantic segmentation of histological data

Segmentation tools for biological applications are widely available (e.g. ZeroCostDL4Mic^6^). For this reason, Spatial µProBe provides auxiliary tools to handle histological data and preprocess image data before the multimodal registration and analysis. Briefly, we include pipelines for dataset generation (i.e. prepare training, validation and test sets by sampling patches from sections, with the option to set custom ratios between existent classes to account for imbalanced datasets, which is often the case for histological sections of bone tissue); hyperparameter optimisation with Optuna^7^ of Deep Learning segmentation models (provided by Segmentation-Models-Pytorch^8^) wrapped in a general segmentation model (“SegModel”) implemented in Pytorch-Lightning^9^; reconstruction of segmentation outputs into the original image and export as a JSON file, compatible with image viewers like QuPath^10^. For dedicated cell segmentation tasks, Spatial µProBe is also compatible with CellPose^11,12^. Dataset generation is accompanied by metadata to indicate which images were considered, and the settings of the patch size and stride, annotation class and patch sampling (exhaustive or balanced) applied. Likewise, hyperparameter optimisation with Optuna is compatible with high-performance computing platforms, enabling users to scale their hyperspace exploration to a predefined number of cores and GPUs (if available). Given the wide range of encoders available in Segmentation-Models-Pytorch, this approach expedites the identification of suitable candidates for the task at hand, especially when combined with an Early Stopping callback that interrupts an iteration when a user-defined metric stops improving for a pre-defined number of training epochs. Additionally, there is an option to have an active SQL database running to save the results of each optimisation step into a table.

Supplementary Figure SN1.1 illustrates an example of the application of this segmentation pipeline to segment bone from histological sections stained for Sclerostin (top row) and Safranin-O (bottom row), and counterstaining performed with Fast Green F7258 (Sigma-Aldrich, St. Louis, MO). A Unet-based SegModel was selected and hyperparameter optimisation was performed considering 6 variations of Mix Vision Transformer^13^ (mit-b0 to mit-b5). Loss functions were based on linear combinations of Jaccard, Dice, Focal, Lovasz, Tversky, and Soft Binary Cross-Entropy with Logits functions, available in Segmentation-Models-Pytorch. Learning rate varied between 1e^-6^ and 1e^-3^ (on a logarithmic scale). An early stopping callback monitored the validation loss, interrupting the training iteration when this parameter stopped improving by at least 0.001 after 25 epochs. Regarding the training dataset, eight sections (3 stained for Sclerostin and 5 for Safranin-O) were used to generate 1284 patches with side 224 µm (448 pixels). Since pixels assigned to bone define a severely imbalanced class, the dataset was created with the following rules: 95% of the patches consisted of foreground (i.e., “bone”), defined as patches with at least 50% of the bone mask annotation; 2.5% of patches were negative foreground (i.e., non-background pixels with tissues that are not bone), defined as patches that did not pass the previous criteria but contained 90-100% of the convex-hull of the bone annotation; background, which contained 0-10% of the bone annotation. For validation and testing sets, 2 pairs of sections (one staining each) were considered and sampled on a regular grid with the same dimensions as the training patches. Additionally, we followed a previous approach to define an upper bound on performance^11^, where we manually annotated three sections from an independent dataset twice, with the second annotation being performed on a vertically mirrored image. This established a reference Dice score of 0.983 ± 0.008 for manual annotation.

In 15 trials, the best performing model was achieved at the 3^rd^ iteration, with a validation Dice score of 0.98, considering a batch size of 26, learning rate of 5.0x10^-5^ and a loss function defined with Dice, Lovasz and Soft Binary Cross Entropy with logits. As represented in Figure SN1.1a, there are very minor misalignments in the segmentation predictions, which nearly vanish when scaling the annotation result to the resolution of in vivo micro-computed tomography (micro-CT) images, emulating the step preceding 2D-3D registration. In other words, despite variations between ground truth and predicted annotations (which can also arise based on different criteria between image annotators), the result that will effectively be used during the analysis shows a nearly perfect agreement, suitable for 2D-3D registration. Any misalignment can also be corrected manually by adjusting the output annotation with QuPath. Quantitatively, in our test set, the success of the segmentation model is supported by a receiver operating characteristic (ROC) curve (Fig. SN1.1b) with an area under the ROC-curve (AUC) of 0.985, an F1-score of 0.960, recall of 0.978 and precision of 0.944 when the maximum Intersection over Union (IoU) of 0.926 is achieved by optimising the threshold used to segment the predicted values (Fig. SN1.1c). These results correspond to a Dice score on the test set of 0.970 ± 0.004.

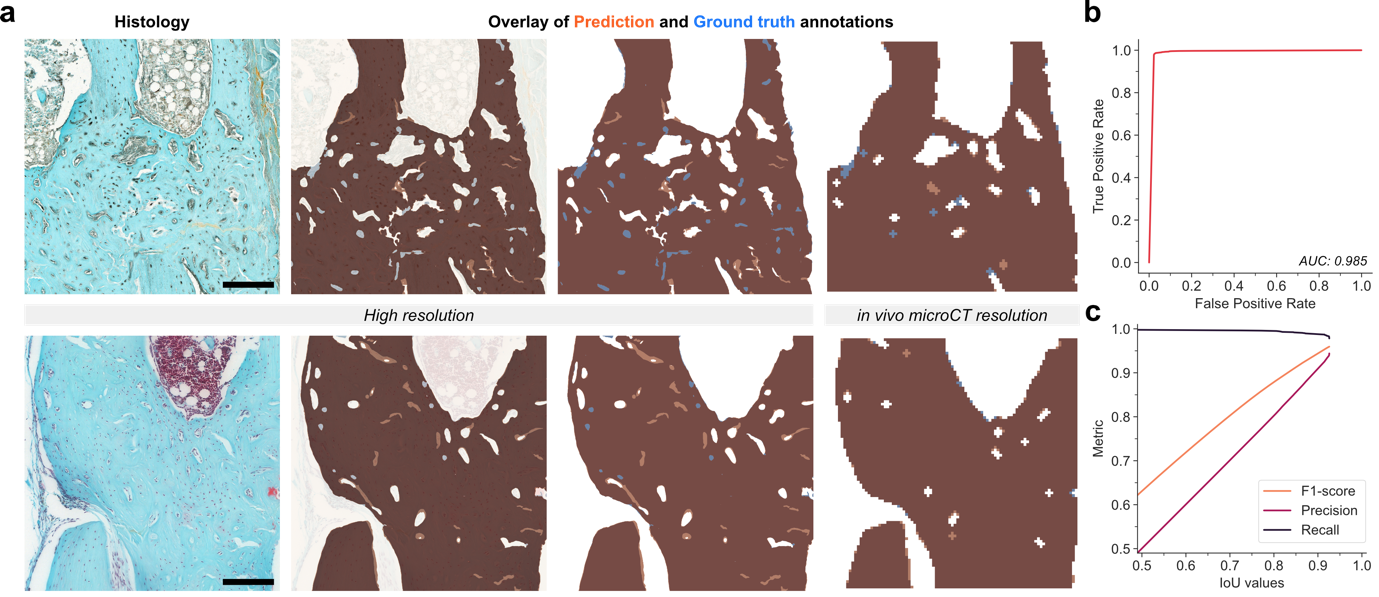

Supplementary Figure SN1.1 | Semantic segmentation of bone from histological sections of mouse femur.

a, Representative example of histological patches segmented with the best performing model identified with Optuna’s hyperparameter optimisation trials (scale bars: 1000 µm). The top row shows a patch from a section stained for Sclerostin and the bottom row represents a patch from a section stained with Safranin-O. Despite small misalignments between the predicted and ground-truth annotations, results are nearly identical when rescaling to the resolution of in vivo micro-computed tomography (micro-CT) images (voxel-size: 10.5 µm), as required for the analysis with Spatial µProBe.

b, Receiver operating characteristic curve for the best performing model identified. An area under the ROC curve (AUC) of 0.985 was achieved.

c, Association between Intersection over Union (IoU) values and relevant evaluation metrics (F1-score, recall and precision). IoU values were generated by varying the threshold used to produce a binary mask by segmenting the output predicted by the model, which was then compared with the ground-truth annotation.

Supplementary Note 2

Validation of 2D-3D registration using simulated data

2.1 Pre-selection of image similarity functions

Multimodal 2D-3D image registration in Spatial µProBe is performed on binary images from each modality, through iterative maximisation of a set of image similarity functions. Given the vast range of options available, a preliminary analysis aimed at identifying a well-performing subset for further analysis in the synthetic experiments evaluating the convergence success of the registration algorithm.

Ideal candidates should have a unique similarity maximum (with positive kurtosis around it, i.e., leptokurtic), the widest dynamic range (in the interval 0 and 1), a smooth monotonic trajectory to reach the maximum point and be uncorrelated with other options (as a proxy for the type of image features captured).

A set of 10 similarity metrics was selected at the start: Dice similarity coefficient (Di), Jaccard index (Ji), Pearson correlation coefficient (Pe), normalised root mean squared error (rMSE), normalised Mutual Information (nMI), surface Dice similarity coefficient (DS); additionally, Pe, rMSE, nMI and the structural similarity index (SSIM) functions were also computed on the Euclidean distance transform (EDT) of the images being registered (EDTPe, EDTrMSE, EDTnMI and EDTSSIM).

Following a simplified pipeline for generation of 2D sections from 3D images, 100 linearly spaced values were computed for 4 transformation parameters (translation along the x-axis and z-axis, within 30 voxels of the range of values in each axis, and rotations along phi and theta angles, between 0º and 180º). For each transformation value, the corresponding section was compared with the sections obtained with the remaining 99 transformations, and the value of the image similarity functions were computed. This process was repeated for all femur and 6^th^ caudal vertebra micro-CT images used in the main validation study. The cross-correlation between similarity functions for each transformation value and orientation was determined and averaged across all samples. Additionally, four statistical metrics were determined for each transformation value: monotonicity index (measuring how consistently the values increase or decrease), peak-to-peak range (the difference between maximum and minimum values), peak sharpness (measuring how pronounced peaks are in the data) and kurtosis (as a proxy for the shape of the data distribution).

This preliminary analysis converged on the following set of similarity functions for subsequent steps: “DS”, “nMI”, “EDTnMI” and “EDTPe”. The similarity functions “Di”, “Ji” and “Pe” were mutually correlated, while also being correlated with “EDTPe” and “nMI” (Supplementary Fig. SN2.1a and SN2.2a). The latter two were selected for subsequent analysis since they also showed favourable and complementary statistical descriptors (Supplementary Table 1 and 2). “DS” was also selected since it was mostly uncorrelated with other candidates and exhibited suitable statistical descriptors. Although “EDTnMI” and “EDTrMSE” showed similar performance, only the former was selected as it consistently displayed a smoother trend across all conditions (Supplementary Fig. SN2.1b and SN2.2b).

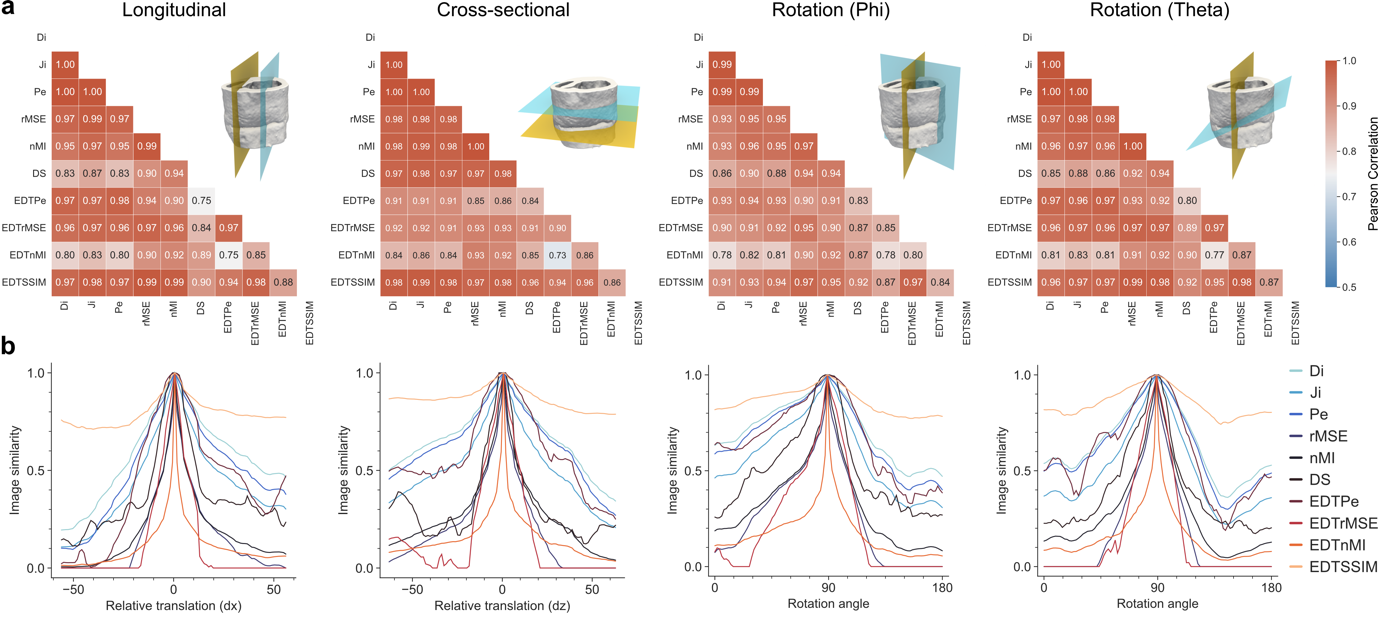

Supplementary Figure SN2.1 | Preselection of image similarity functions for 2D-3D registration, based on mouse femur samples.

a, Pearson correlation between image similarity functions, for four transformations: longitudinal translation, cross-sectional translation, rotation around phi (longitudinal axis), rotation around theta (cross-sectional axis). Values represent the average correlation between pairs of image similarity functions across four femur samples used in the synthetic study. For each sample and orientation, 100 sections were generated. The following similarity functions are considered: Dice similarity coefficient (Di), Jaccard index (Ji), Pearson correlation coefficient (Pe), normalised root mean squared error (rMSE), normalised Mutual Information (nMI), surface Dice similarity coefficient (DS); additionally, Pe, rMSE, nMI and the structural similarity index (SSIM) functions were also computed on the Euclidean distance transform (EDT) of the images being registered (EDTPe, EDTrMSE, EDTnMI and EDTSSIM). 3D visualisations show two sections illustrating the type of operation performed during section generation.

b, Representative example of the profile of the image similarity functions for four transformations shown in sub-panel a), for the median transformation value. The figure highlights the differences in monotonicity, peak uniqueness and sharpness for different functions.

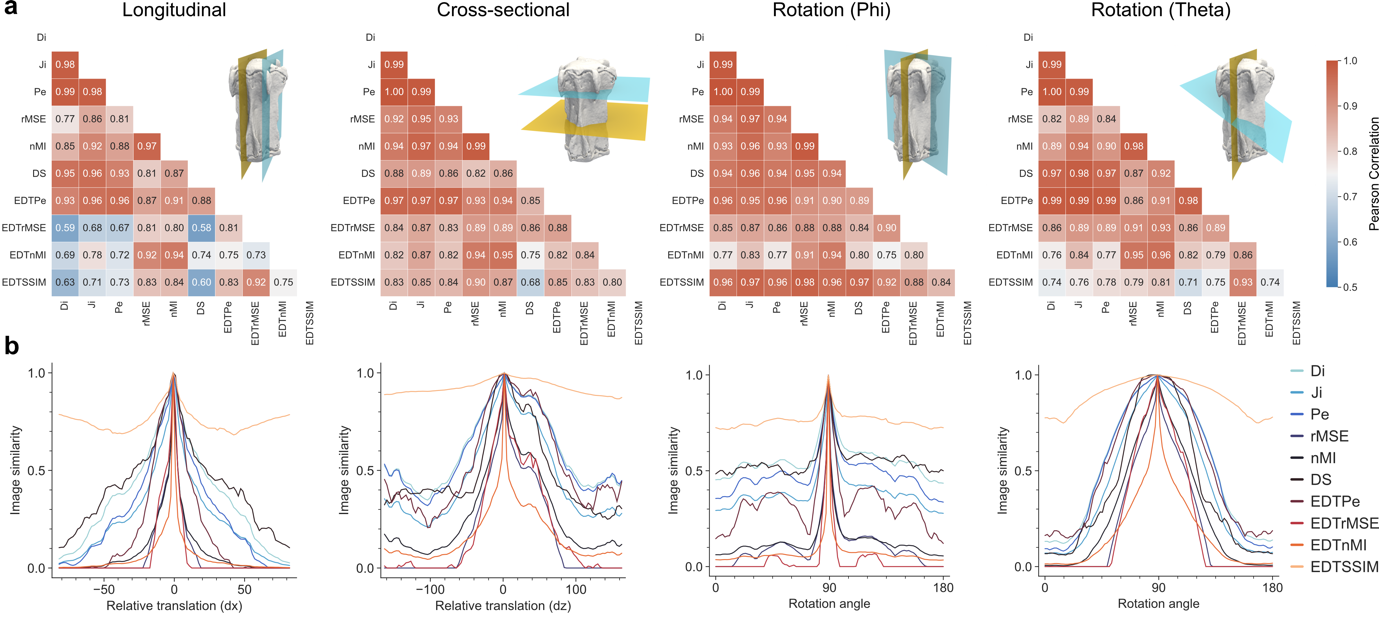

Supplementary Figure SN2.2 | Preselection of image similarity functions for 2D-3D registration, based on mouse 6^th^ caudal vertebra samples.

a, Pearson correlation between image similarity functions, for four transformations: longitudinal translation, cross-sectional translation, rotation around phi (longitudinal axis), rotation around theta (cross-sectional axis). Values represent the average correlation between pairs of image similarity functions across four 6^th^ caudal vertebra samples used in the synthetic study. For each sample and orientation, 100 sections were generated. The following similarity functions are considered: Dice similarity coefficient (Di), Jaccard index (Ji), Pearson correlation coefficient (Pe), normalised root mean squared error (rMSE), normalised Mutual Information (nMI), surface Dice similarity coefficient (DS); additionally, Pe, rMSE, nMI and the structural similarity index (SSIM) functions were also computed on the Euclidean distance transform (EDT) of the images being registered (EDTPe, EDTrMSE, EDTnMI and EDTSSIM). 3D visualisations show two sections illustrating the type of operation performed during section generation.

b, Representative example of the profile of the image similarity functions for four transformations shown in sub-panel a), for the median transformation value. The figure highlights the differences in monotonicity, peak uniqueness and sharpness for different functions.

|  | **Longitudinal** | | | |  | **Cross-sectional** | | | |
| --- | --- | --- | --- | --- | --- | --- | --- | --- | --- |
| Function | Monotonicity index | Peak-to-peak | Peak sharpness | Kurtosis |  | Monotonicity index | Peak-to-peak | Peak sharpness | Kurtosis |
| Di | 0.582 ± 0.046 | 0.744 ± 0.219 | 1.087 ± 0.057 | 2.615 ± 0.646 |  | **0.479 ± 0.035** | 0.448 ± 0.217 | 1.042 ± 0.027 | 2.247 ± 0.523 |
| Ji | 0.582 ± 0.046 | 0.838 ± 0.166 | 1.168 ± 0.108 | 3.457 ± 1.519 |  | **0.479 ± 0.035** | 0.593 ± 0.234 | 1.083 ± 0.052 | 2.497 ± 0.907 |
| Pe | **0.584 ± 0.044** | 0.849 ± 0.194 | 1.098 ± 0.059 | 2.549 ± 0.615 |  | **0.481 ± 0.032** | 0.496 ± 0.231 | 1.047 ± 0.028 | 2.252 ± 0.543 |
| rMSE | 0.367 ± 0.177 | 0.974 ± 0.053 | **1.575 ± 0.291** | **7.086 ± 4.387** |  | 0.462 ± 0.056 | **0.873 ± 0.233** | **1.346 ± 0.152** | **3.505 ± 1.160** |
| nMI | 0.581 ± 0.041 | **0.977 ± 0.041** | **1.503 ± 0.247** | 6.777 ± 3.604 |  | **0.481 ± 0.032** | 0.825 ± 0.165 | **1.269 ± 0.129** | **4.043 ± 2.045** |
| DS | 0.539 ± 0.073 | 0.884 ± 0.060 | 1.100 ± 0.052 | 4.223 ± 0.653 |  | **0.485 ± 0.058** | 0.790 ± 0.077 | 1.051 ± 0.055 | 2.749 ± 1.510 |
| EDTPe | **0.608 ± 0.053** | 0.921 ± 0.155 | 1.072 ± 0.054 | 2.622 ± 0.827 |  | 0.460 ± 0.017 | 0.517 ± 0.244 | 1.016 ± 0.010 | 2.180 ± 0.360 |
| EDTrMSE | 0.296 ± 0.177 | **1.000 ± 0.000** | 1.417 ± 0.236 | **7.041 ± 4.136** |  | 0.411 ± 0.102 | **0.874 ± 0.198** | 1.165 ± 0.083 | 2.641 ± 1.128 |
| EDTnMI | **0.588 ± 0.047** | **0.978 ± 0.025** | **2.318 ± 0.256** | **19.199 ± 5.243** |  | 0.467 ± 0.028 | **0.904 ± 0.060** | **1.897 ± 0.191** | **12.796 ± 3.502** |
| EDTSSIM | 0.547 ± 0.026 | 0.286 ± 0.046 | 1.045 ± 0.012 | 3.105 ± 0.774 |  | 0.438 ± 0.080 | 0.174 ± 0.027 | 1.019 ± 0.007 | 2.338 ± 0.499 |
|  | ***Phi*** | | | |  | ***Theta*** | | | |
| Function | Monotonicity index | Peak-to-peak | Peak sharpness | Kurtosis |  | Monotonicity index | Peak-to-peak | Peak sharpness | Kurtosis |
| Di | 0.428 ± 0.073 | 0.597 ± 0.071 | 1.077 ± 0.029 | 2.349 ± 0.772 |  | 0.480 ± 0.050 | 0.645 ± 0.225 | 1.031 ± 0.015 | 1.602 ± 0.268 |
| Ji | 0.428 ± 0.073 | 0.746 ± 0.056 | 1.150 ± 0.056 | 3.017 ± 1.223 |  | 0.480 ± 0.050 | 0.765 ± 0.187 | 1.061 ± 0.030 | 1.878 ± 0.385 |
| Pe | 0.424 ± 0.069 | 0.650 ± 0.079 | 1.086 ± 0.033 | 2.403 ± 0.864 |  | 0.476 ± 0.051 | 0.724 ± 0.210 | 1.035 ± 0.015 | 1.600 ± 0.274 |
| rMSE | 0.382 ± 0.064 | **1.000 ± 0.000** | **1.539 ± 0.153** | 5.183 ± 2.599 |  | 0.387 ± 0.117 | **0.970 ± 0.061** | **1.288 ± 0.093** | **2.901 ± 0.932** |
| nMI | 0.420 ± 0.069 | **0.931 ± 0.027** | **1.463 ± 0.154** | **5.849 ± 2.683** |  | 0.475 ± 0.050 | **0.946 ± 0.065** | **1.210 ± 0.069** | **2.908 ± 0.802** |
| DS | **0.441 ± 0.022** | 0.792 ± 0.043 | 1.139 ± 0.098 | 5.173 ± 2.758 |  | **0.497 ± 0.067** | 0.786 ± 0.052 | 1.040 ± 0.036 | 2.622 ± 0.446 |
| EDTPe | **0.481 ± 0.112** | 0.716 ± 0.164 | 1.056 ± 0.023 | 2.396 ± 0.684 |  | 0.456 ± 0.036 | 0.769 ± 0.304 | 1.015 ± 0.010 | 1.584 ± 0.304 |
| EDTrMSE | 0.363 ± 0.164 | **1.000 ± 0.000** | 1.419 ± 0.149 | **5.643 ± 2.560** |  | 0.352 ± 0.105 | **0.962 ± 0.076** | 1.163 ± 0.080 | 2.819 ± 1.081 |
| EDTnMI | **0.431 ± 0.081** | **0.951 ± 0.015** | **2.331 ± 0.232** | **21.650 ± 7.191** |  | **0.480 ± 0.047** | **0.962 ± 0.034** | **1.820 ± 0.112** | **7.878 ± 1.956** |
| EDTSSIM | 0.395 ± 0.024 | 0.226 ± 0.054 | 1.043 ± 0.014 | 3.712 ± 2.724 |  | **0.481 ± 0.044** | 0.248 ± 0.027 | 1.015 ± 0.003 | 1.918 ± 0.261 |

Supplementary Table 1 | Analytical descriptors of image similarity functions for 2D-3D registration for mouse femur samples.

Values of “Monotonicity index”, “Peak-to-peak”, “Peak sharpness” and “Kurtosis” averaged across 100 simulated transformations and four mouse femur samples, for four single-axis transformations (longitudinal, cross-sectional translations and rotations around phi/longitudinal axis and theta/cross-sectional axis). The top 3 values for each column are highlighted in bold. The image similarity functions “EDTnMI”, “nMI”, “rMSE”, “DS” show consistently high performance across multiple transformations.

|  | **Longitudinal** | | | |  | **Cross-sectional** | | | |
| --- | --- | --- | --- | --- | --- | --- | --- | --- | --- |
| Function | Monotonicity index | Peak-to-peak | Peak sharpness | Kurtosis |  | Monotonicity index | Peak-to-peak | Peak sharpness | Kurtosis |
| Di | 0.482 ± 0.033 | 0.973 ± 0.011 | 1.225 ± 0.040 | 2.730 ± 0.197 |  | 0.509 ± 0.024 | 0.665 ± 0.037 | 1.076 ± 0.014 | 1.910 ± 0.148 |
| Ji | 0.482 ± 0.033 | **0.986 ± 0.006** | 1.424 ± 0.073 | 4.747 ± 0.565 |  | 0.509 ± 0.024 | 0.798 ± 0.027 | 1.150 ± 0.027 | 2.385 ± 0.274 |
| Pe | 0.472 ± 0.025 | **1.000 ± 0.000** | 1.269 ± 0.048 | 3.139 ± 0.220 |  | **0.518 ± 0.025** | 0.692 ± 0.036 | 1.080 ± 0.015 | 1.935 ± 0.148 |
| rMSE | 0.193 ± 0.016 | **1.000 ± 0.000** | **2.312 ± 0.206** | **18.028 ± 3.585** |  | 0.352 ± 0.014 | **1.000 ± 0.000** | **1.541 ± 0.068** | **4.013 ± 0.682** |
| nMI | **0.484 ± 0.029** | **1.000 ± 0.000** | **2.265 ± 0.187** | 16.854 ± 2.925 |  | **0.515 ± 0.025** | **0.935 ± 0.012** | 1.388 ± 0.063 | 3.859 ± 0.659 |
| DS | 0.483 ± 0.033 | 0.915 ± 0.025 | 1.175 ± 0.034 | 2.633 ± 0.289 |  | 0.450 ± 0.024 | 0.728 ± 0.033 | 1.083 ± 0.034 | 3.349 ± 0.659 |
| EDTPe | 0.415 ± 0.066 | **1.000 ± 0.000** | 1.266 ± 0.058 | 4.666 ± 0.453 |  | **0.531 ± 0.028** | 0.790 ± 0.024 | 1.054 ± 0.020 | 1.983 ± 0.184 |
| EDTrMSE | 0.141 ± 0.019 | **1.000 ± 0.000** | 2.260 ± 0.230 | **16.875 ± 3.211** |  | 0.452 ± 0.039 | **1.000 ± 0.000** | **1.399 ± 0.081** | **3.980 ± 0.806** |
| EDTnMI | **0.484 ± 0.037** | **0.998 ± 0.000** | **3.465 ± 0.231** | **42.381 ± 4.634** |  | 0.511 ± 0.025 | **0.957 ± 0.006** | **2.123 ± 0.129** | **12.762 ± 2.796** |
| EDTSSIM | **0.504 ± 0.028** | 0.310 ± 0.015 | 1.126 ± 0.018 | 6.661 ± 0.605 |  | 0.488 ± 0.010 | 0.134 ± 0.003 | 1.016 ± 0.003 | 2.126 ± 0.139 |
|  | ***Phi*** | | | |  | ***Theta*** | | | |
| Function | Monotonicity index | Peak-to-peak | Peak sharpness | Kurtosis |  | Monotonicity index | Peak-to-peak | Peak sharpness | Kurtosis |
| Di | 0.487 ± 0.043 | 0.612 ± 0.034 | 1.283 ± 0.032 | 8.630 ± 0.495 |  | 0.509 ± 0.017 | 0.883 ± 0.007 | 1.025 ± 0.003 | 1.421 ± 0.036 |
| Ji | 0.487 ± 0.043 | 0.759 ± 0.026 | 1.529 ± 0.058 | 12.649 ± 0.806 |  | 0.509 ± 0.017 | 0.938 ± 0.004 | 1.051 ± 0.006 | 1.643 ± 0.078 |
| Pe | 0.493 ± 0.033 | 0.705 ± 0.030 | 1.342 ± 0.034 | 8.482 ± 0.543 |  | 0.510 ± 0.019 | 0.922 ± 0.005 | 1.027 ± 0.003 | 1.427 ± 0.036 |
| rMSE | 0.396 ± 0.083 | **1.000 ± 0.000** | **2.657 ± 0.211** | 24.226 ± 3.474 |  | 0.341 ± 0.012 | **1.000 ± 0.000** | **1.259 ± 0.018** | **2.664 ± 0.198** |
| nMI | 0.494 ± 0.028 | **0.955 ± 0.008** | 2.585 ± 0.131 | **29.142 ± 2.838** |  | **0.509 ± 0.018** | **0.996 ± 0.000** | 1.150 ± 0.016 | 2.314 ± 0.161 |
| DS | **0.540 ± 0.092** | 0.497 ± 0.023 | 1.281 ± 0.045 | 12.157 ± 1.483 |  | 0.501 ± 0.014 | 0.943 ± 0.008 | 1.015 ± 0.014 | 1.839 ± 0.026 |
| EDTPe | **0.531 ± 0.054** | 0.909 ± 0.027 | 1.500 ± 0.053 | 8.420 ± 0.380 |  | **0.510 ± 0.021** | 0.862 ± 0.016 | 1.015 ± 0.009 | 1.516 ± 0.098 |
| EDTrMSE | 0.164 ± 0.086 | **1.000 ± 0.000** | **3.595 ± 0.436** | **33.568 ± 5.465** |  | 0.318 ± 0.017 | **1.000 ± 0.000** | **1.167 ± 0.046** | **2.800 ± 0.361** |
| EDTnMI | 0.504 ± 0.020 | **0.971 ± 0.003** | **4.088 ± 0.136** | **64.777 ± 3.854** |  | **0.508 ± 0.014** | **0.989 ± 0.001** | **1.598 ± 0.071** | **4.911 ± 0.459** |
| EDTSSIM | **0.521 ± 0.029** | 0.271 ± 0.015 | 1.156 ± 0.012 | 12.254 ± 1.036 |  | 0.502 ± 0.013 | 0.251 ± 0.017 | 1.005 ± 0.001 | 1.637 ± 0.039 |

Supplementary Table 2 | Analytical descriptors of image similarity functions for 2D-3D registration for mouse 6^th^ caudal vertebra samples.

Values of “Monotonicity index”, “Peak-to-peak”, “Peak sharpness” and “Kurtosis” averaged across 100 simulated transformations and four mouse vertebra samples, for four single-axis transformations (longitudinal, cross-sectional translations and rotations around phi/longitudinal axis and theta/cross-sectional axis). The top 3 values for each column are highlighted in bold. The image similarity functions “nMI”, “EDTnMI”, “rMSE”, “EDTrMSE” and “EDTPe” show consistently high performance across multiple transformations.

2.2 Analysis of 2D-3D registration using in silico datasets

In this section, we first complement the results presented in Figure 2 and Extended Fig. 1 related to the in silico validation of the 2D-3D registration algorithm of Spatial µProBe. Next, we include the same analysis performed on mouse 6^th^ caudal vertebra micro-CT images. These vertebra micro-CT scans typically have bigger dimensions (in number of voxels) than femur images and, due to the presence of intricate trabecular architectures, provide detailed features for image registration.

Femur cross-sectional sections were generally more robust to rotational and translational offsets, sustaining excellent convergence rates until 45-60º and 450-600 µm, as opposed to longitudinal sections where convergence success starts to drop within 15-30º and 150-300 µm, respectively. Cross-sectional sections typically required more complex combinations of similarity functions to converge than longitudinal sections, except for the sample with the highest bone volume fraction (BV/TV; Supplementary Fig. SN2.3). Expectedly, non-linear deformations strongly affected the convergence success, with sections initialised with larger rotational and translational offsets from the ground-truth transformation showing lower convergence rates (Supplementary Fig. SN2.4-2.5). Success decreased sharply, from over 90% to below 10%, for sections initialised close to their target location but with increasing deformation magnitudes (Supplementary Fig. SN2.4, 0-150 µm), whereas sections with mild deformations (σ < 3) still showed acceptable convergence for translational offsets up to 450 µm (Supplementary Fig. SN2.4). Registration of cross-sectional sections was more robust when analysing images with small deformations (σ < 3) across all samples, reaching high convergence success rates even for trials initialised with translational offsets above 600 µm (Supplementary Fig. SN2.5). Highly deformed sections (σ = 5) showed virtually no success, indicating that such cases must be registered iteratively with user intervention.

Sections generated from the mouse 6^th^ caudal vertebra showed a similar performance as the femur sections (Supplementary Fig. SN2.6a-b, g-h), with cross-sectional sections showing consistently high convergence rates across all BV/TV values (Supplementary Fig. SN2.7). Attempts initialised with rotational offsets up to 30º showed higher success (> 90%) than translational offsets up to 250 µm, even for sections with non-linear deformations (Supplementary Fig. SN2.6c-f, i-l). Similar to femur samples, vertebral cross-sectional sections typically required more complex combinations of similarity functions to converge than longitudinal sections. Notably, a single similarity function (DS) was sufficient to achieve the highest convergence rates for challenging conditions with high deformations or initialisation offsets (Supplementary Fig. SN2.7, SN2.8). When paired with EDTnMI, this similarity function showed the best performance for longitudinal and cross-sectional sections initialised with translational offsets above 250 µm and 750 µm, respectively (Supplementary Fig. SN2.8, SN2.9). For smaller translational offsets (< 250 µm), across all rotational offsets, EDTnMI and nMI were particularly successful for mild deformations (σ < 1).

Overall, this analysis showed that unsupervised Iterative Assisted Registration (IterAR) can recover the location of a 2D section in its 3D image, with confident success for moderate translational (< 250 µm) and rotational (< 30º) initialisation offsets. Furthermore, successful results were observed for both mouse femur and caudal vertebra, highlighting the versatility of this approach and independence from image structure. Finally, registration trials driven by the combination of EDTnMI/nMI and DS image similarity functions were particularly successful, supporting its use for real applications.

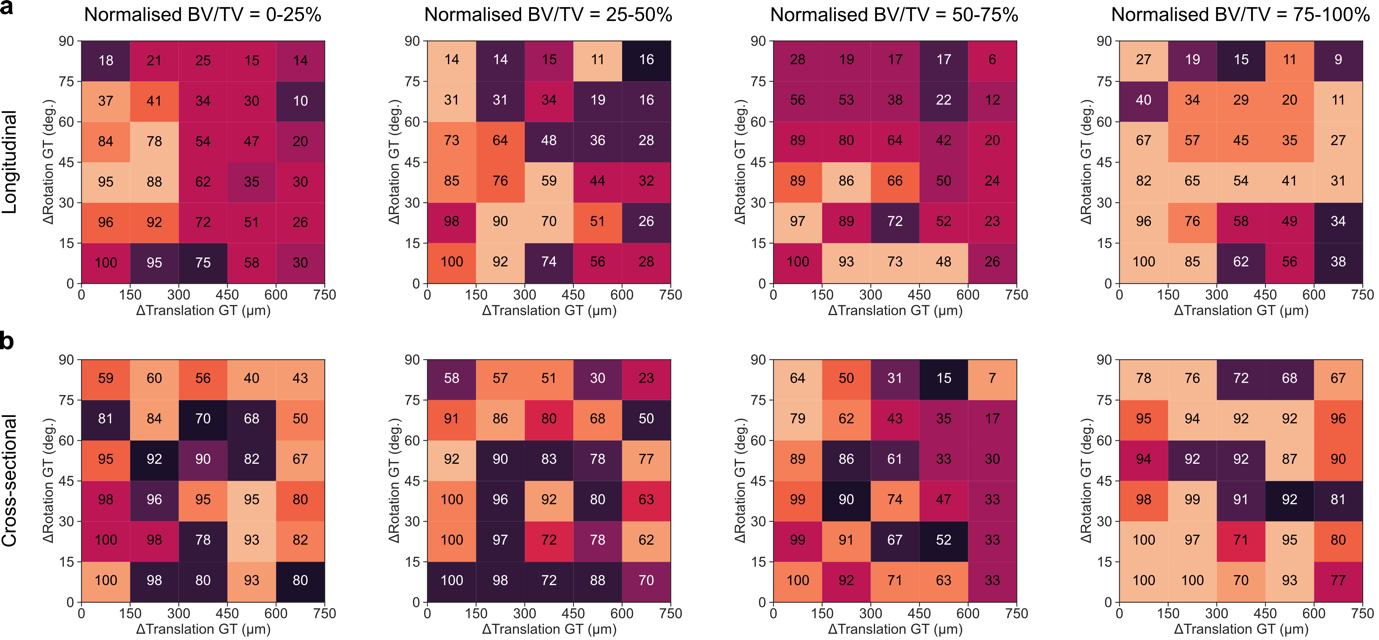

Supplementary Figure SN2.3 | Convergence success rate of the 2D-3D registration algorithm for the analysis of sections generated with rigid transformations from mouse femur samples.

a, Convergence success rates (%) for sections initialised at different translational and rotational offsets from the ground-truth (GT) transformation, for sections generated along the longitudinal orientation across images with increasing bone volume fraction (BV/TV). The background colours of each bin indicate the simplest combination of image similarity functions that achieved the score indicated (remaining combinations not shown). Colour legend provided at the end of this section.

b, Convergence success rates (%) for sections initialised at different translational and rotational offsets from the ground-truth transformation, for sections generated along the cross-sectional orientation. The same colour code rule was applied as in sub-panel a).

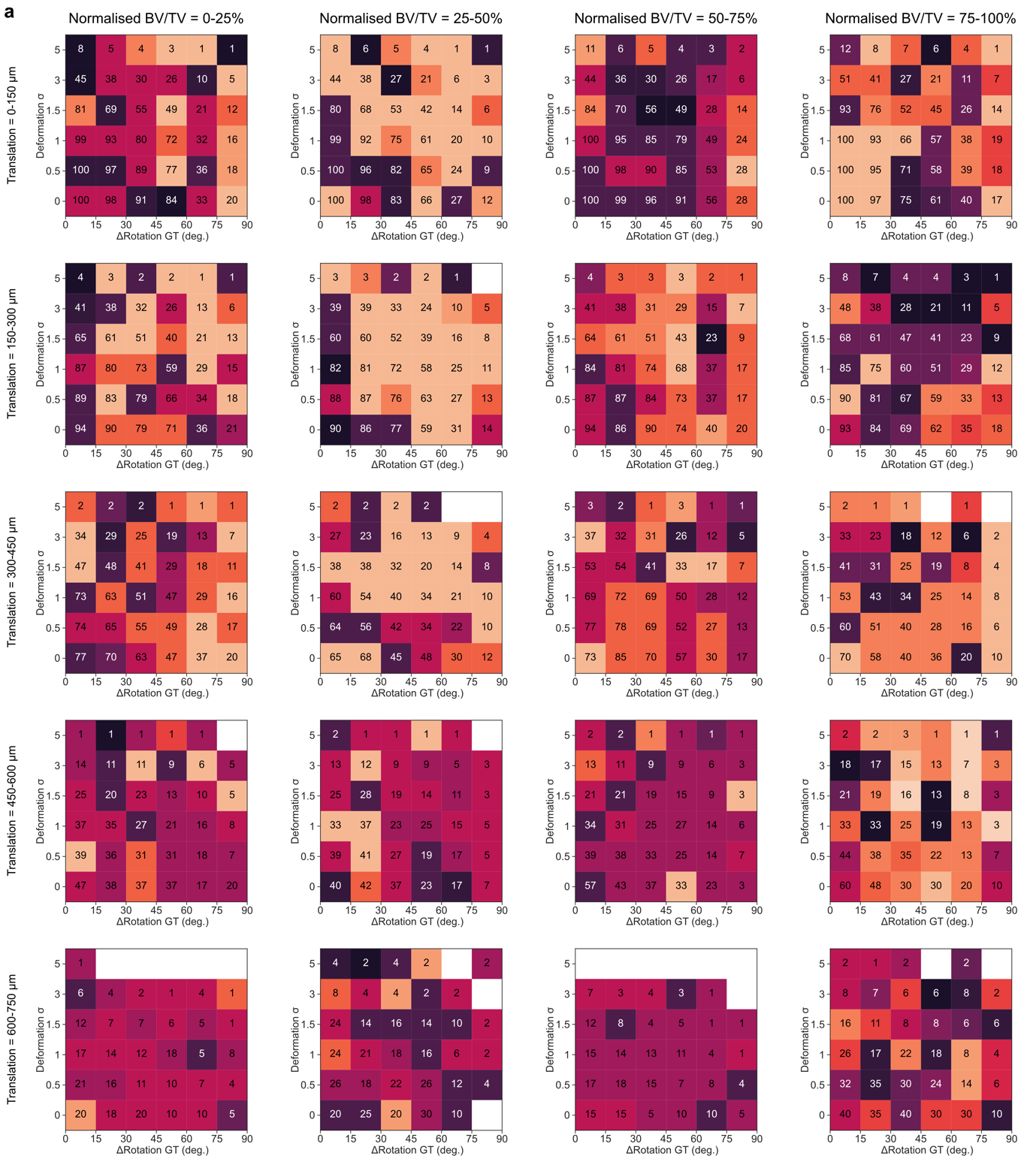

Supplementary Figure SN2.4 | Convergence success rate of the 2D-3D registration algorithm for the analysis of sections generated with rigid and non-linear transformations from mouse femur samples (longitudinal orientation).

a, Convergence success rates (%) across varying deformation settings (σ) and rotational offsets, for longitudinal sections, and including synthetically generated non-linear deformations with increasing magnitude (increasing σ). Each row indicates sections initialised with increasing translational offsets from the ground-truth (GT) transformation. Colour code of each bin as in SN2.3. White boxes represent no successful results (0%). Columns refer to images with increasing bone volume fraction (BV/TV), normalised in quartiles across all samples (see Methods).

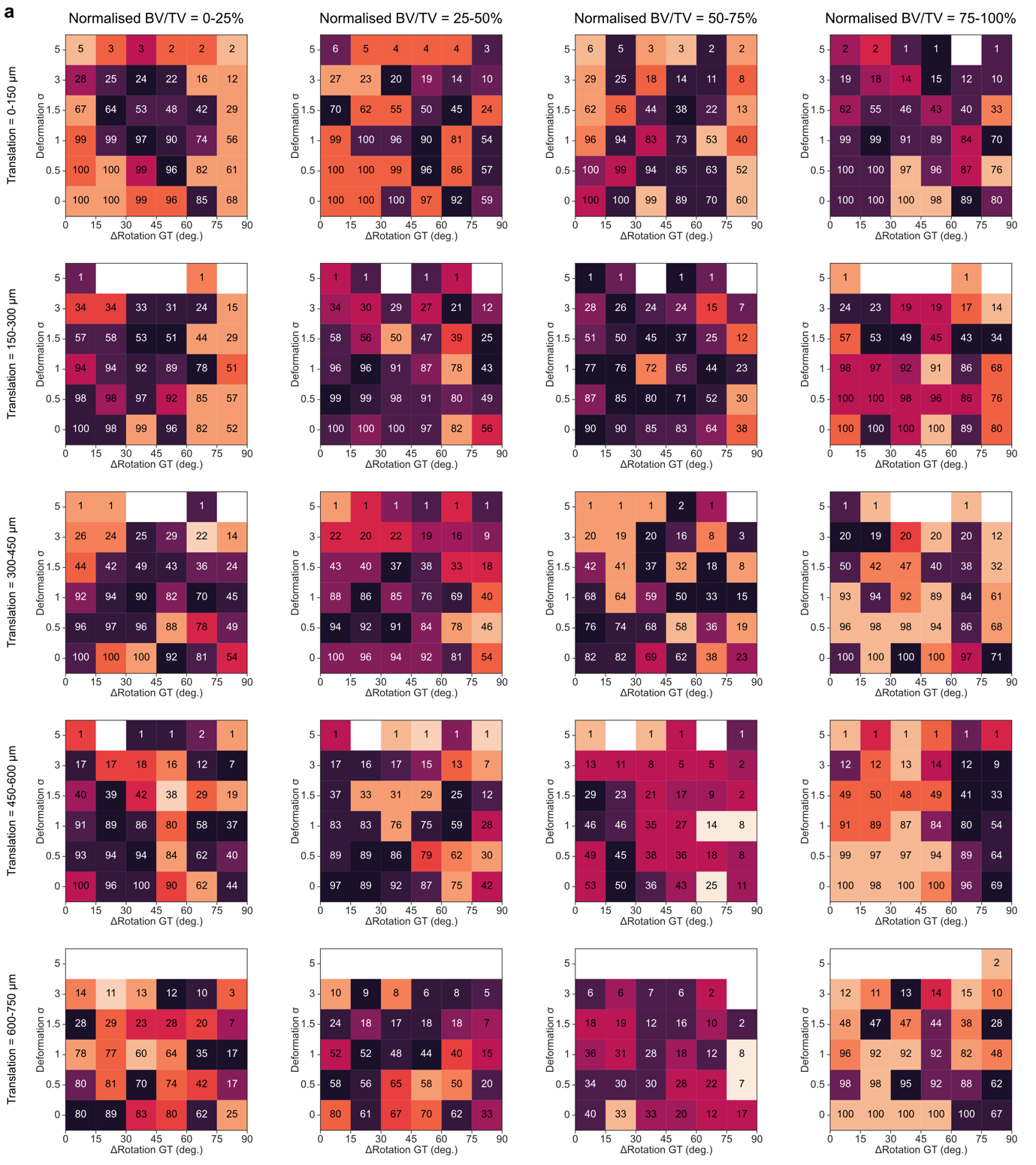

Supplementary Figure SN2.5 | Convergence success rate of the 2D-3D registration algorithm for the analysis of sections generated with rigid and non-linear transformations from mouse femur samples (cross-sectional orientation).

a, Convergence success rates (%) across varying deformation settings (σ) and rotational offsets, for sections generated along the cross-sectional orientation and including synthetically generated non-linear deformations with increasing magnitude (increasing σ). Each row indicates sections initialised with increasing translational offsets from the ground-truth (GT) transformation. Colour code of each bin as in SN2.3. Columns refer to images with increasing bone volume fraction (BV/TV), normalised in quartiles across all samples (see Methods).

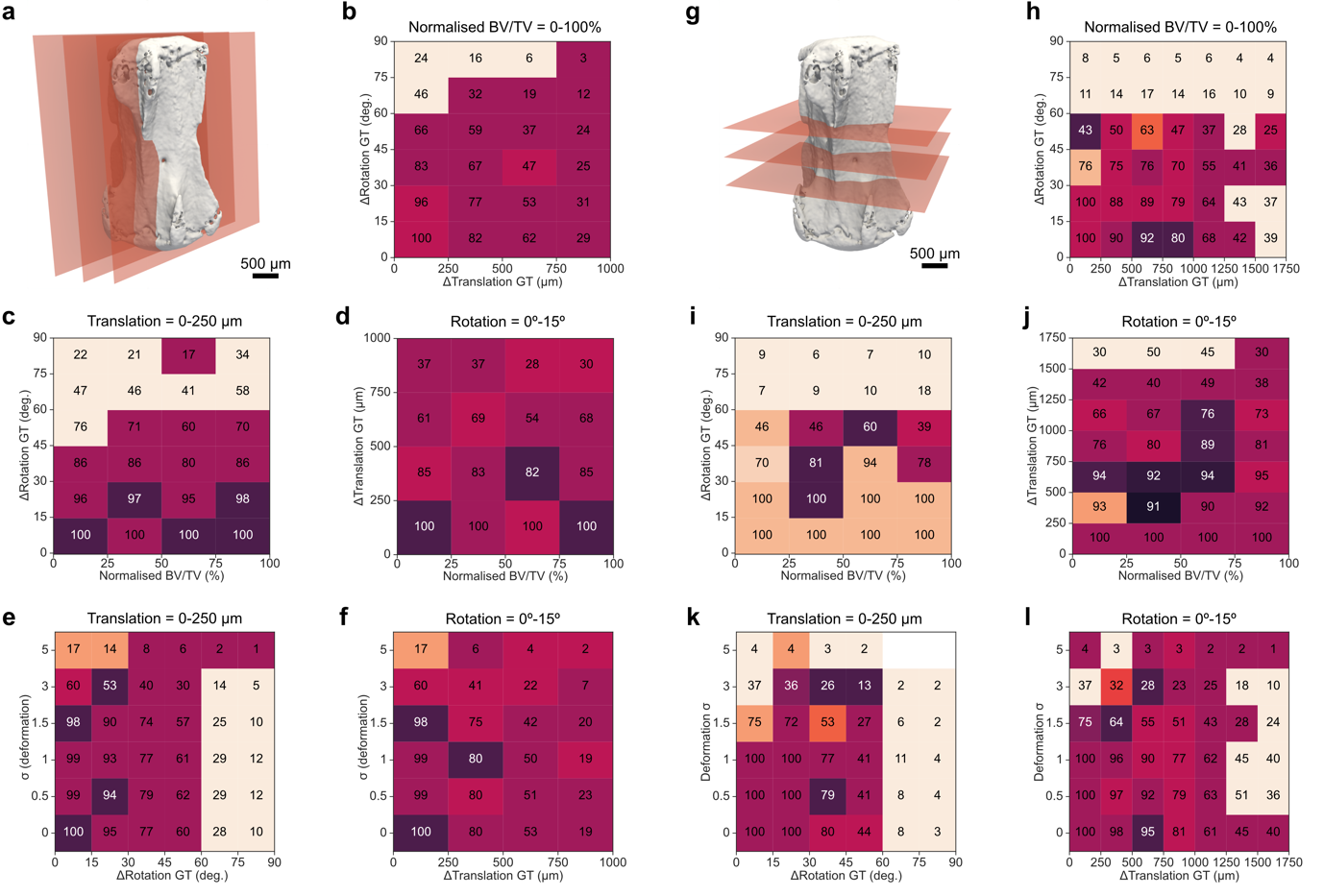

Supplementary Figure SN2.6 | Validation of 2D-3D registration in Spatial µProBe using sections generated from mouse 6^th^ caudal vertebra samples.

a, Illustration of the generation of synthetic datasets of 2D sections from 3D micro-computed tomography (micro-CT) images of the 6^th^ caudal vertebra, for longitudinal sections.

b, Convergence success rates (%) across all samples analysed, binned in intervals of 250 µm and 15º of translational and rotational initialisation offsets to the ground-truth (GT) transformation, for sections generated along the longitudinal orientation. The background colours of each bin indicate the simplest combination of image similarity functions that achieved the score indicated (remaining combinations not shown). The average of the images across all the normalised bone volume fraction (BV/TV) values is presented (0-100%).

c, Convergence success rates (%) for sections initialised with a maximum of 250 µm translational offset to the ground-truth transformation, for sections generated along the longitudinal orientation. Scores are binned in intervals of 15º rotational offset to the ground-truth transformation in the y-axis, and with BV/TV across the samples considered in the x-axis (one sample per column). Colour code of each bin as in sub-panel b).

d, Convergence success rates (%) for sections initialised with a maximum of 15º rotational offset to the ground-truth transformation, for sections generated along the longitudinal orientation. Scores are binned in intervals of 250 µm translational offset to the ground-truth transformation in the y-axis, and with the normalised BV/TV across the samples considered in the x-axis (one sample per column). Colour code of each bin as in sub-panel b).

e, Convergence success rates (%) across varying deformation settings (σ) and rotational offsets, for sections initialised with a translation within 250 μm of the ground-truth transformation, generated along the longitudinal orientation. Colour code of each bin as in sub-panel b).

f, Convergence success rates (%) across varying deformation settings (σ) and rotational offsets, for sections initialised with a 15º rotational offset to the ground-truth transformation, generated along the longitudinal orientation. Colour code of each bin as in sub-panel b).

g, h, i, j, k, l show the corresponding results as in a, b, c, d, e, f, respectively, for 2D sections from 3D micro-CT images of the 6^th^ caudal vertebra along the cross-sectional orientation.

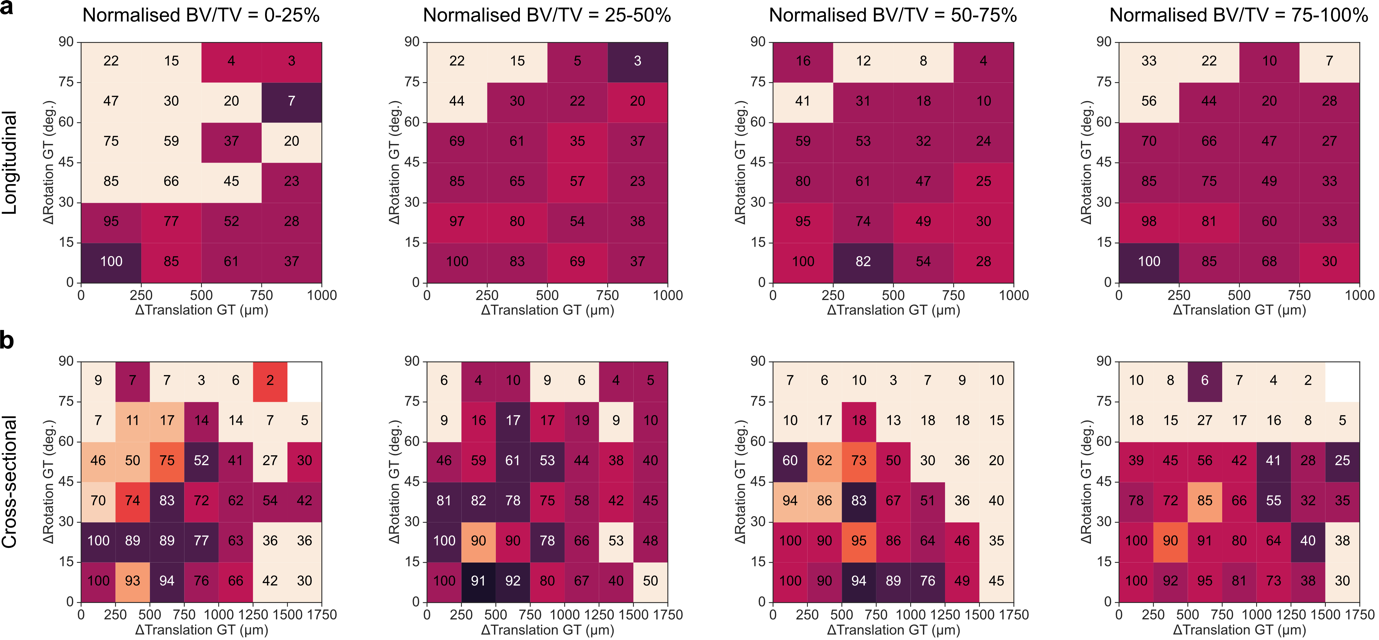

Supplementary Figure SN2.7 | Convergence success rate of the 2D-3D registration algorithm for the analysis of sections generated with rigid transformations from mouse 6^th^ caudal vertebra samples.

a, Convergence success rates (%) for sections initialised at different translational and rotational offsets from the ground-truth (GT) transformation, for sections generated along the longitudinal orientation. The background colours of each bin indicate the simplest combination of image similarity functions that achieved the score indicated (remaining combinations not shown). Colour legend provided at the end of this section. Columns refer to images with increasing bone volume fraction (BV/TV), normalised in quartiles across all samples (see Methods).

b, Convergence success rates (%) for sections initialised at different translational and rotational offsets from the ground-truth transformation, for cross-sectional sections. The same colour code rule was applied as in sub-panel a).

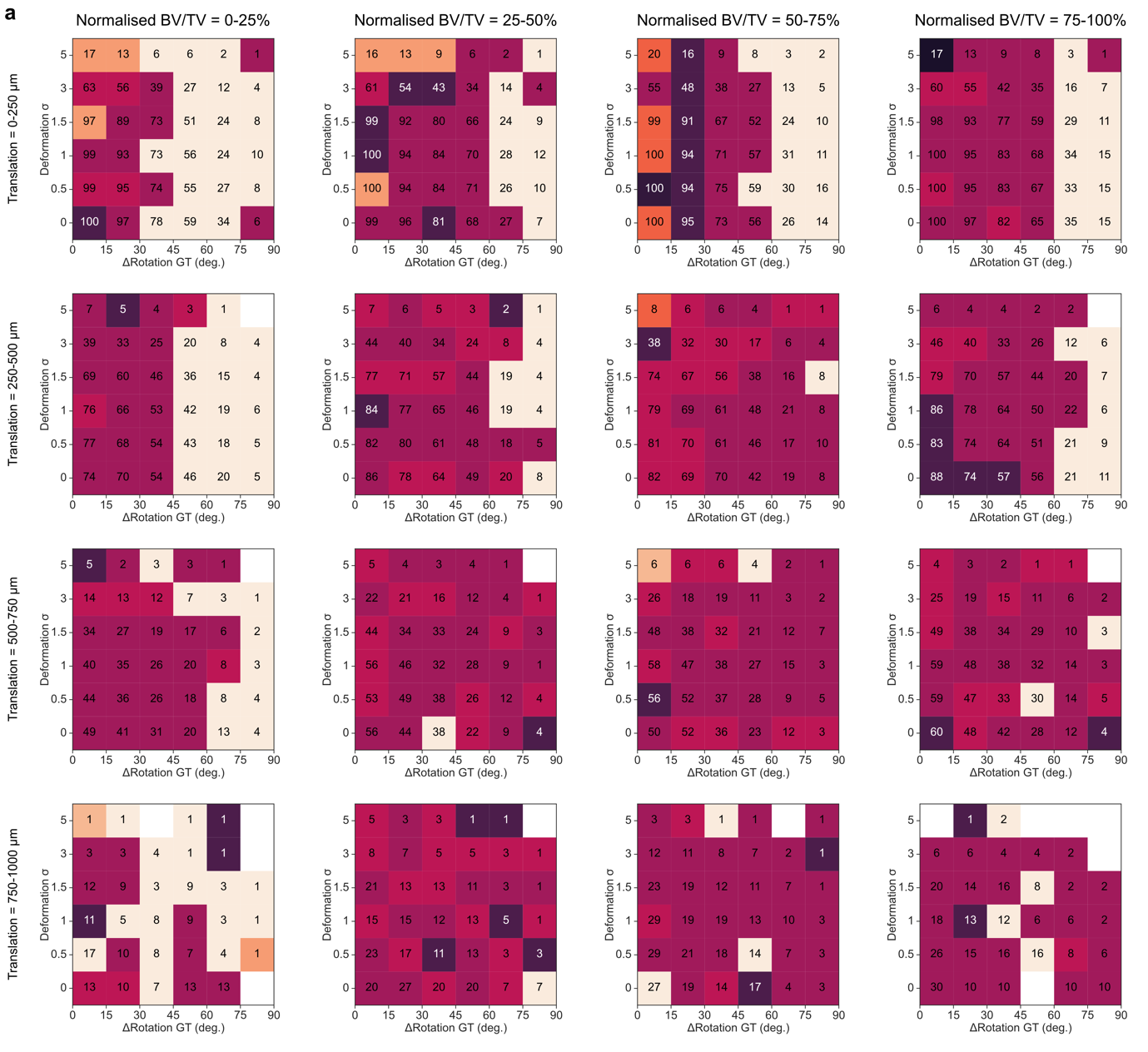

Supplementary Figure SN2.8 | Convergence success rate of the 2D-3D registration algorithm for the analysis of sections generated with rigid and non-linear transformations from mouse 6^th^ caudal vertebra samples (longitudinal orientation).

a, Convergence success rates (%) across varying deformation settings (σ) and rotational offsets, for sections generated along the longitudinal orientation and including synthetically generated non-linear deformations with increasing magnitude (increasing σ). Each row indicates sections initialised with increasing translational offsets from the ground-truth (GT) transformation. Colour code of each bin as in SN2.3. Columns refer to images with increasing bone volume fraction (BV/TV), normalised in quartiles across all samples (see Methods).

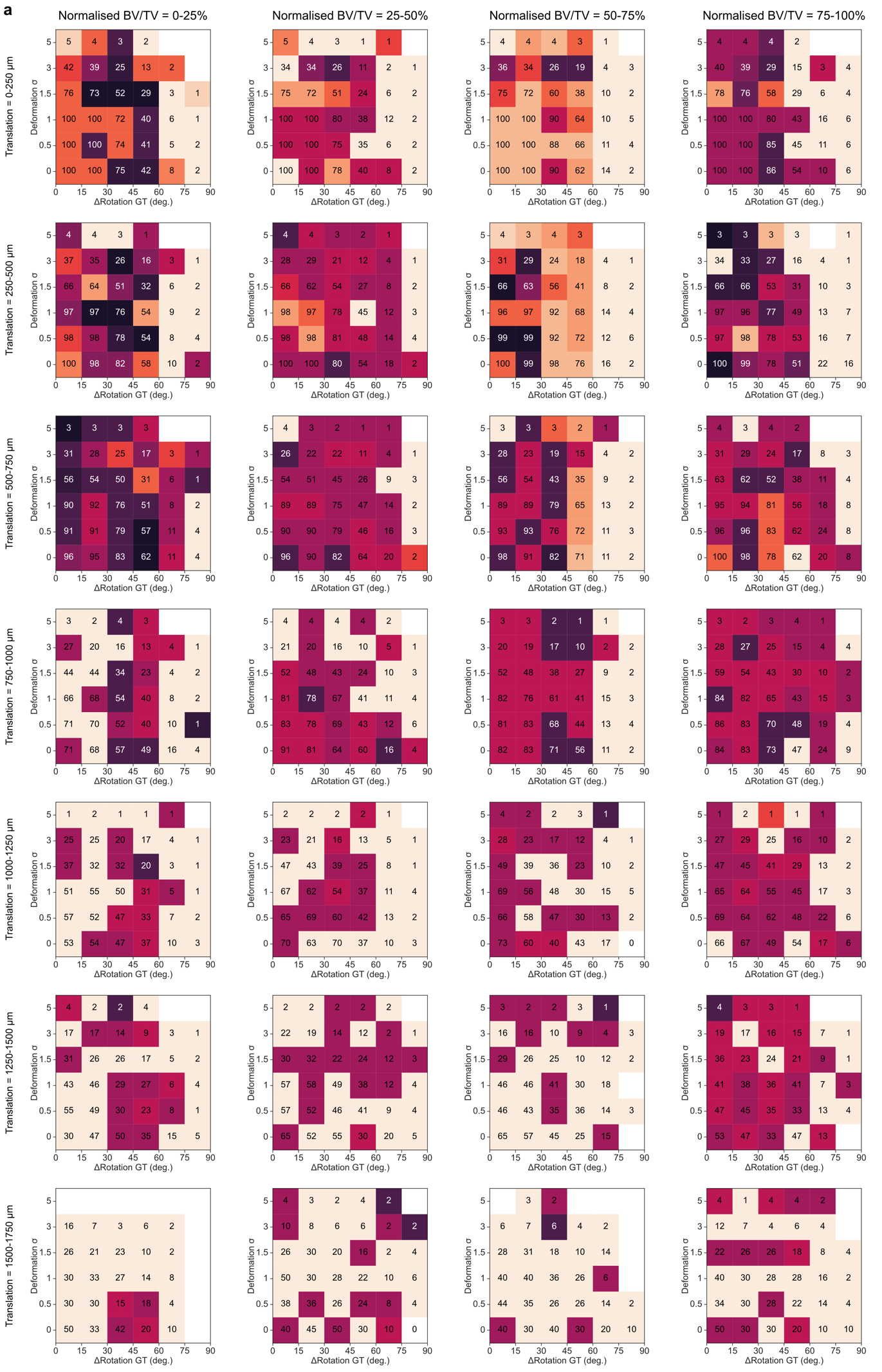

Supplementary Figure SN2.9 | Convergence success rate of the 2D-3D registration algorithm for the analysis of sections generated with rigid and non-linear transformations from mouse 6^th^ caudal vertebra samples (cross-sectional orientation).

a, Convergence success rates (%) across varying deformation settings (σ) and rotational offsets, for sections generated along the cross-sectional orientation and including synthetically generated non-linear deformations with increasing magnitude (increasing σ). Each row indicates sections initialised with increasing translational offsets from the ground-truth (GT) transformation. Colour code of each bin as in SN2.3. Columns refer to images with increasing bone volume fraction (BV/TV), normalised in quartiles across all samples (see Methods).

Colour legend for Supplementary Figures SN2.3 to SN2.9:

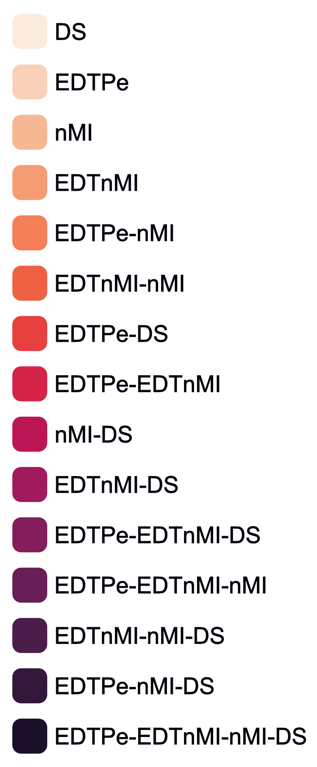

Supplementary Note 3

Super-resolution of Visium spatial transcriptomics data

Spatial µProBe leverages recent advances in super-resolution of 10x Genomics Visium spatial transcriptomics (ST) data to predict gene expression at the resolution of in vivo micro-CT scans (10.5 µm), from the standard resolution of 55 µm diameter spots (with 100 µm centre-to-centre distance) in one capture area. Specifically, we used iStar^2^ and we evaluated the feasibility of this tool to predict gene expression data in our bone fracture healing dataset, while conserving the original magnitude of the gene expression data for subsequent analysis. Since 3D datasets (e.g. spatially heterogeneous mechanical signal from micro-finite element analysis) are produced at the micro-CT resolution, super-resolved ST data can effectively be linked to all values of the 2D section obtained from the registered 3D dataset, rather than being limited to the sparse spot locations of the standard Visium ST data.

We compared the performance of super-resolved (Super-res) with standard Visium (Visium-res) ST data across three categories relevant to the mechanoregulation analysis performed in our work: correlation between gene expression modelled with generalised additive models (GAMs) using the Super-res data (versus Visium-res) and bone formation time-points or local mechanical signals obtained from 3D datasets; accuracy (measured with the mean absolute error) of GAMs produced with Super-res and Visium-res data; and, the width of the 95% confidence interval of the average expression predicted with the GAMs from Super-res and Visium-res data. For each sample (Control and Loaded, see Methods), the analysis was performed on spatially variable genes (SVGs) identified with SPARK-X^4^. Additionally, the same pipeline was considered for relative cell-type proportions predicted with Starfysh^1^, for both samples.

Our analysis showed that GAM curves associating gene expression with local mechanical signals are highly correlated for both Control and Loaded samples (Supplementary Fig. SN3.1a, SN3.1e and SN3.2a, SN3.2e), with 89% of the SVGs identified showing a Spearman correlation above 0.5 between GAMs obtained with Super-res and Visium-res, compared to 67% and 75% for GAMs associating gene expression and time of Control and Loaded samples, respectively. Genes with lower correlations were predominantly associated with lower mean expression values, suggesting that stricter filtering of SVGs based on expression should improve these outcomes. Here, we opted for a relatively lenient filter to favour the exploratory potential of this approach. When comparing the mean absolute error (MAE) between Super-res and Visium-res GAM models, most genes show comparable values (Supplementary Fig. SN3.1b, SN3.1f and SN3.2b, SN3.2f), with a small subset showing an increased MAE for Super-res GAMs. This result is plausible since MAE was computed with respect to the raw Visium data for both types of GAM models. Given that Super-res expression data can produce small shifts in the original distribution of values, a GAM created from Super-res data is likely to exhibit a higher MAE against Visium data when compared with a GAM modelled with Visium data (which is predicting the same data it was created with). Notably, a clear advantage of super-resolution is the improved precision and confidence in the trends obtained with GAMs, as shown by significantly narrower 95% confidence intervals, while retaining a strong agreement in the trends displayed by GAM predicted curves (Supplementary Fig. SN3.1d, SN3.1h and SN3.2d, SN3.2h).

Comparably, super-resolution of relative cell-type proportions estimated with Starfysh exhibit similar performance with 67% and 87% of the cell-types analysed showing a correlation above 0.5 for the Control and Loaded samples, respectively, for GAMs associating relative proportion and mechanical signals. For GAMs linking relative proportion and time, these values improve to 87% and 93% for Control and Loaded samples, respectively. In this case, normalised MAE values were predominantly higher for the GAMs produced at Visium resolution, with super-resolution also maintaining improved precision, as shown in the significantly narrower 95% confidence intervals of the predictions produced by the GAM (Supplementary Fig. SN3.3, SN3.4).

The input relative proportions estimated with Starfysh are bounded between 0 and 1, which coincides with the prediction range of iStar. Therefore, no post-processing scaling is required, which explains the improved performance of super-resolution for this application. Comparably, since the input gene expression data presented above shows considerably different ranges, iStar super-resolution predictions must account for this increased variability during training which increases the prediction error and deviations to ground-truth Visium expression data. As recommended by the authors of iStar^2^, highly expressed and variable genes should be prioritised to improve signal-to-noise ratio during model training. Nonetheless, as mentioned above, we opted for highlighting the exploratory potential of our analysis with more permissive gene selection criteria, which can still be further optimised in combination with more comprehensive ST datasets provided as input. The results showed that GAMs produced from super-resolution data provide insightful representations of gene expression data and relative proportion of cell-types, with quantitative advantages over analysis performed at standard Visium resolution.

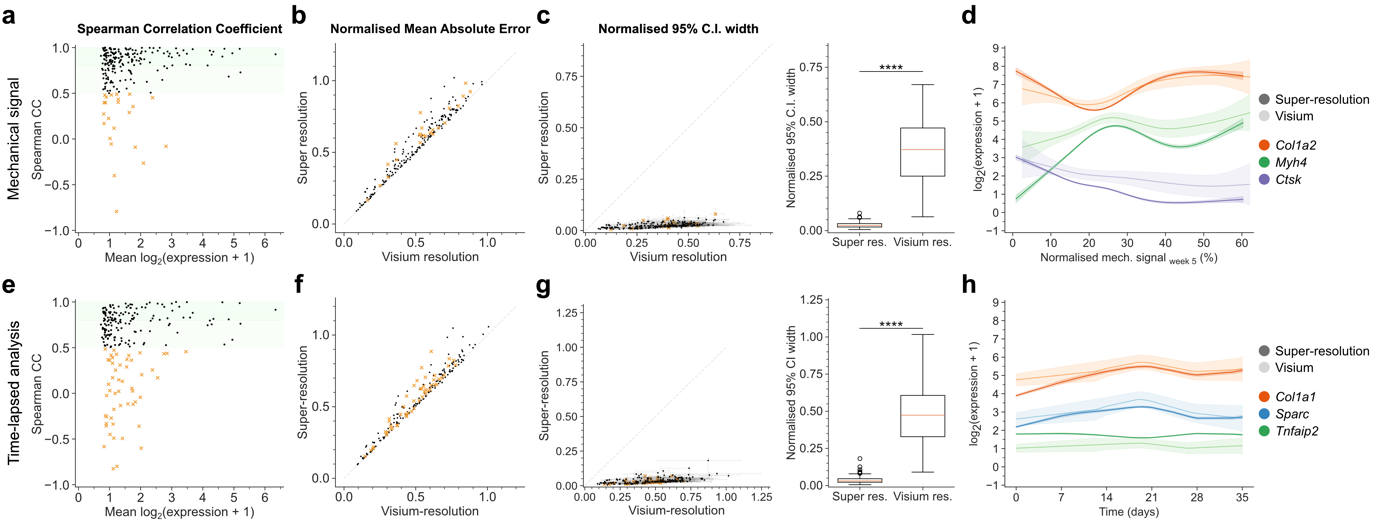

Supplementary Figure SN3.1 | Evaluation of iStar super-resolution performance on ST data from the Loaded sample.

a, Spearman correlation coefficient (CC) between the predicted expression value obtained with a generalised additive model (GAM) of gene expression as a function of mechanical signals, when using super-resolution gene expression and Visium data. Data plotted as a function of the mean log_2_(expression + 1). Only spatially variable genes (SVGs) identified with SPARK-X were considered. Two increasingly darker shades of green in the background indicate correlations above 0.5 and 0.8 respectively. Genes with a correlation above 0.5 are indicated with a black dot, while those with a correlation below 0.5 are indicated with an orange cross.

b, Mean absolute error (MAE) normalised by the mean expression (from Visium data) for all SVGs between GAMs created with super-resolution and Visium data. Genes with a correlation above 0.5 are indicated with a black dot, while those with a correlation below 0.5 are indicated with an orange cross.

c, Median and inter-quartile range of the 95% confidence interval (C.I.) normalised by the predicted expression at each point, for all SVGs, for GAMs created with super-resolution and Visium data. Genes with a correlation below 0.5 identified in a are marked with an orange cross. The same data is represented as a box-plot on the right (****, p < 0.0001).

d, Examples of GAMs produced with super-resolution and Visium data for selected markers. GAMs created with super-resolution reproduce the trend of the GAMs from Visium data while achieving much improved precision.

e, f, g and h show the same analysis as a, b, c and but for GAMs associating gene expression data across time, estimated from time-lapsed micro-CT images.

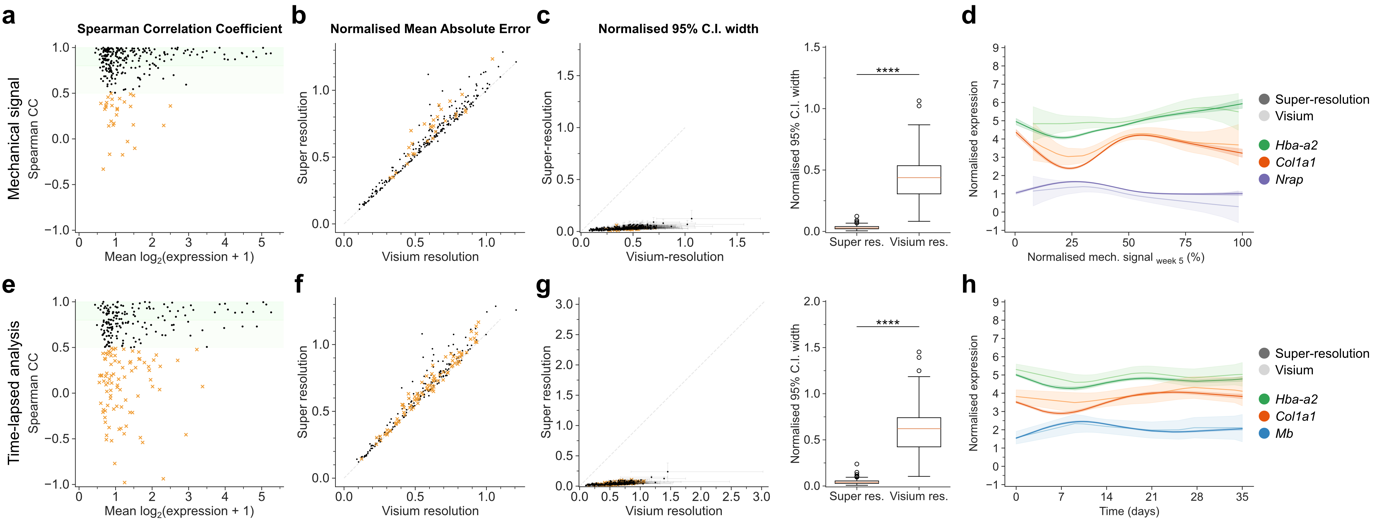

Supplementary Figure SN3.2 | Evaluation of iStar super-resolution performance on ST data from the Control sample.

a, Spearman correlation coefficient (CC) between the predicted expression values obtained with a generalised additive model (GAM) of gene expression as a function of mechanical signals, when using super-resolution gene expression and Visium data. Data plotted as a function of the mean log_2_(expression + 1). Only spatially variable genes (SVGs) identified with SPARK-X were considered. Two increasingly darker shades of green indicate correlations above 0.5 and 0.8 respectively. Genes with a correlation below 0.5 are indicated with an orange cross.

b, Mean absolute error (MAE) normalised by the mean expression (from Visium data) for all SVGs between GAMs created with super-resolution and Visium data. Genes with a correlation below 0.5 identified in sub-panel a) are marked with an orange cross.

c, Median and inter-quartile range of the 95% confidence interval (C.I.) normalised by the predicted expression at each point, for all SVGs, for GAMs created with super-resolution and Visium data. Genes with a correlation below 0.5 identified in sub-panel a) are marked with an orange cross. The same data is represented as a box-plot on the right (****, p < 0.0001).

d, Examples of GAMs produced with super-resolution and Visium data for selected markers. GAMs created with super-resolution reproduce the trend of the GAMs from Visium data while achieving much improved precision.

e, f, g and h show the same analysis as a, b, c and but for GAMs associating gene expression data across time, estimated from time-lapsed micro-computed tomography (micro-CT) images.

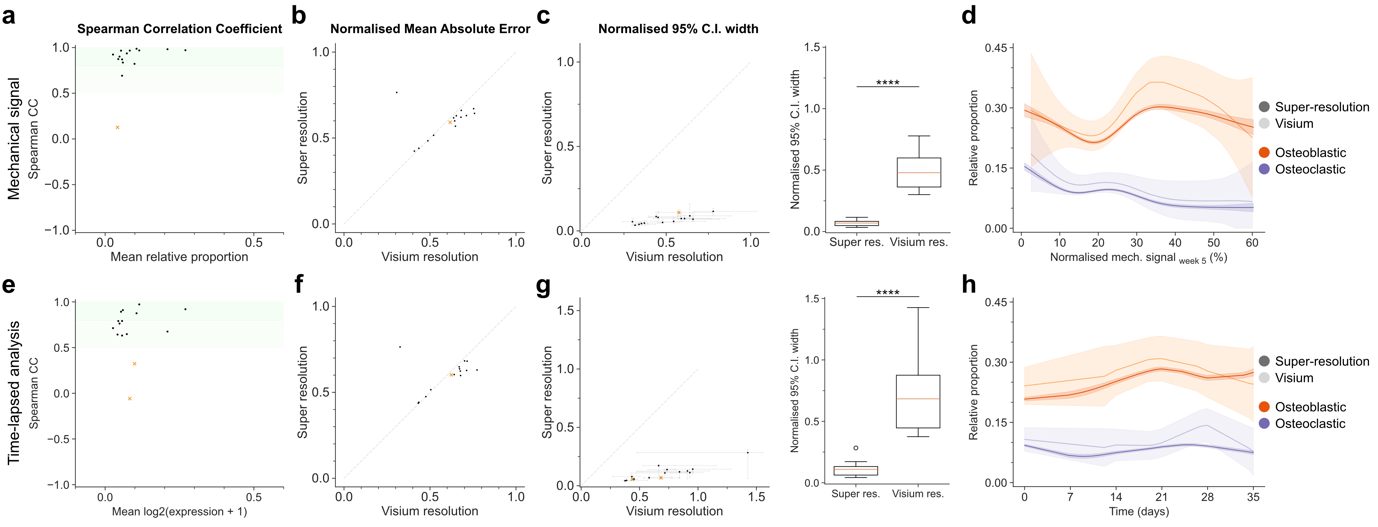

Supplementary Figure SN3.3 | Evaluation of iStar super-resolution of Starfysh cell-type relative proportions estimated from ST data of the Loaded sample.

a, Spearman correlation coefficient (CC) between the predicted cell-type relative proportions obtained with a GAM associating cell-type relative proportion as a function of mechanical signals, when using super-resolution and Visium resolution data of the output produced by Starfysh. Data plotted as a function of the mean relative proportion of each cell type. Two increasingly darker shades of green indicate correlations above 0.5 and 0.8 respectively. Cell types with a correlation below 0.5 are indicated with an orange cross.

b, Mean absolute error (MAE) normalised by the mean relative proportion (from Visium data) for all cell-types between generalised additive models (GAMs) created with super-resolution and Visium resolution data obtained from Starfysh. Cell types with a correlation below 0.5 are indicated with an orange cross.

c, Median and inter-quartile range of the 95% confidence interval normalised by the predicted relative proportion, for all cell types analysed, for GAMs created with super-resolution and Visium data produced by Starfysh. Cell types with a correlation below 0.5 are indicated with an orange cross. The same data is represented as a box-plot on the right (****, p < 0.0001).

d, Examples of GAMs produced with super-resolution and Visium data for selected cell types.

e, f, g and h show the same analysis as a, b, c and but for GAMs associating relative proportion of cell-types data across time, estimated from time-lapsed micro-computed tomography (micro-CT) images.

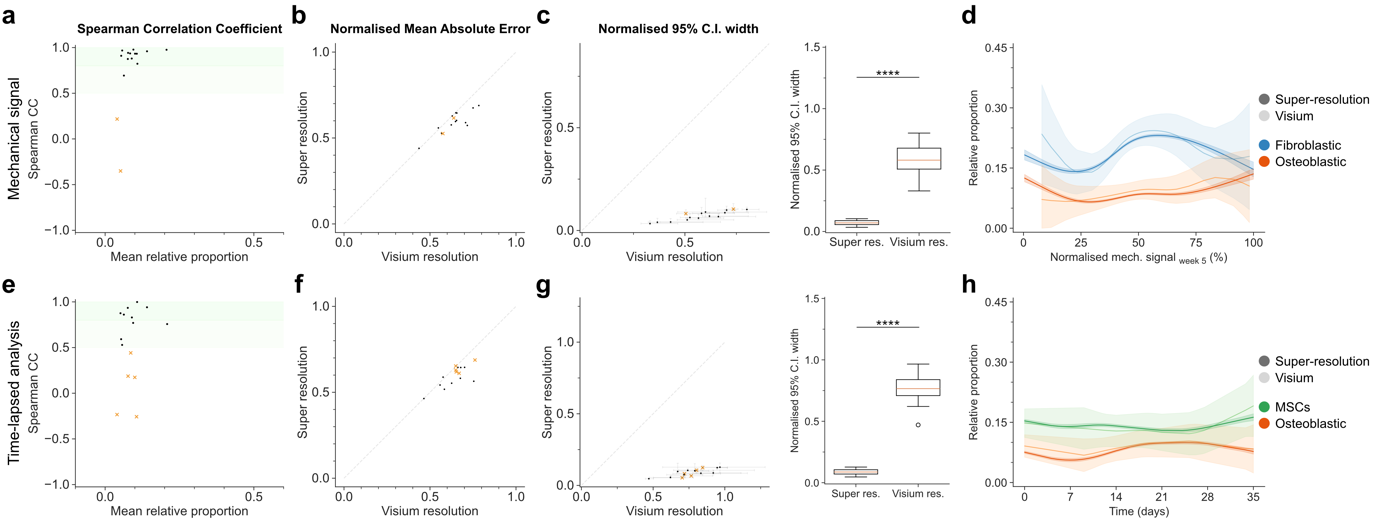

Supplementary Figure SN3.4 | Evaluation of iStar super-resolution of Starfysh cell-type relative proportions estimated from ST data of the Control sample.

a, Spearman correlation coefficient (CC) between the predicted cell-type relative proportions obtained with a generalised additive model (GAM) of gene expression as a function of mechanical signals, when super-resolution and Visium resolution of the output produced by Starfysh. Data plotted as a function of the mean relative proportion of each cell type. Two increasingly darker shades of green indicate correlations above 0.5 and 0.8 respectively. Cell types with a correlation below 0.5 are indicated with an orange cross.

b, Mean absolute error normalised by the mean relative proportion (from Visium data) for all cell-types between GAMs created with super-resolution and Visium resolution data obtained from Starfysh. Cell types with a correlation below 0.5 are indicated with an orange cross.

c, Median and inter-quartile range of the 95% confidence interval (C.I.) normalised by the predicted relative proportion, for all cell types analysed, for GAMs created with super-resolution and Visium data produced by Starfysh. Cell types with a correlation below 0.5 are indicated with an orange cross. The same data is represented as a box-plot on the right (****, p < 0.0001).

d, Examples of GAMs produced with super-resolution and Visium data for selected cell types.

e, f, g and h show the same analysis as a, b, c and but for GAMs associating relative proportion of cell-types data across time, estimated from time-lapsed micro-computed tomography (micro-CT) images.
